# Integrative phenotypic and genomic analyses reveal strain-dependent responses to acute ozone exposure and their associations with airway macrophage transcriptional activity

**DOI:** 10.1101/2021.01.29.428733

**Authors:** Adelaide Tovar, Wesley L. Crouse, Gregory J. Smith, Joseph M. Thomas, Benjamin P. Keith, Kathryn M. McFadden, Timothy P. Moran, Terrence S. Furey, Samir N. P. Kelada

## Abstract

Acute ozone (O_3_) exposure is associated with multiple adverse cardiorespiratory outcomes, the severity of which varies across human populations and rodent models from diverse genetic backgrounds. However, molecular determinants of response, including biomarkers that distinguish which individuals will develop more severe injury and inflammation (*i.e.*, high responders), are poorly characterized. Here, we exposed adult, female and male mice from 6 strains, including 5 Collaborative Cross (CC) strains, to filtered air (FA) or 2 ppm O_3_ for 3 hours, and measured several inflammatory and injury parameters 21 hours later. Additionally, we collected airway macrophages and performed RNA-seq analysis to investigate influences of strain, treatment, and strain-by-treatment interactions on gene expression as well as transcriptional correlates of lung phenotypes. Animals exposed to O_3_ developed airway neutrophilia and lung injury, with varying degrees of severity. We identified many genes that were altered by O_3_ exposure across all strains, and examination of genes whose expression was influenced by strain-by-treatment interactions revealed prominent differences in response between the CC017/Unc and CC003/Unc strains, which were low- and high-responders, respectively (as measured by cellular inflammation and injury). Further investigation of this contrast indicated that baseline gene expression differences likely contribute to their divergent post-O_3_ exposure transcriptional responses. We also observed alterations in chromatin accessibility that differed by strain and with strain-by-treatment interactions, lending further plausibility that baseline differences can modulate post-exposure responses. Together, these results suggest that aspects of the respiratory response to O_3_ exposure may be mediated through altered airway macrophage transcriptional signatures, and further confirms the importance of gene-by-environment interactions in mediating differential responsiveness to environmental agents.

## Introduction

For decades, ambient ozone (O_3_) exposure has been recognized as a hazard to human health. Acute O_3_ exposure causes decrements in pulmonary function, tissue injury and inflammation in healthy individuals, and these effects are more severe in individuals living with respiratory disease. Moreover, emerging evidence links long-term O_3_ exposure to development of chronic diseases and disorders including asthma (1–3) and allergic rhinitis (4). As such, identifying those with enhanced risk and establishing the mechanisms by which O_3_ exposure promotes and exacerbates disease are important public health objectives.

Controlled exposure studies performed as early as the 1980s have demonstrated that O_3_-induced respiratory responses are highly reproducible within individuals (including when exposures are separated by long time intervals (5)) and yet quite variable across a population, even after participant characteristics such as sex, age, and smoking and disease statuses are matched (5–8). Genetic polymorphisms that interact with O_3_ exposure (“gene-environment interaction” or “GxE”) likely contribute to observed inter-individual variation (9, 10), a notion supported both by studies measuring O_3_ responses among genetically variable inbred rodent strains (11–16) and through demonstration of the effects of natural genetic variation on responses to other environmental stimuli (17–22). Estimating individual-level causal effects of toxicant exposures remains challenging and often impossible to achieve in human studies. Although this hurdle can be overcome with the use of animal models, most studies examining effects of O_3_ exposure have used a single common inbred mouse strain C57BL/6J, limiting their power to make broader claims about core drivers of O_3_-induced responses across genetically diverse individuals or uncover the biology governing these divergent responses.

To address this research gap, we exposed a panel of Collaborative Cross (CC) recombinant inbred mouse strains that span a broad phenotypic range of responses to acute O_3_ exposure and performed genomic analyses of airway macrophages, a key cell type known to mount dynamic responses to O_3_ (23–28). The CC is a mouse genetic reference population derived from eight founder strains, whose genomes collectively harbor around 40 million single-nucleotide variants (SNVs) (29, 30), thus capturing more than 90% of the genetic variation present in all inbred mouse strains (31). This resource has been used to identify novel candidate genes and variants associated with drug-induced adverse effects (32, 33), responses to pollutants (34), and respiratory innate immunity (35–37). Additionally, the CC has been used to study transcriptional correlates of disease features (38–40). Thus, this population is appropriate for understanding genetic contributions to complex disease phenotypes and dissecting causal mechanisms of O_3_ toxicity.

Here, we exploit response heterogeneity within inbred mouse strains in conjunction with inflammatory and injury phenotyping and genomic profiling of airway macrophages to: (1) define a core set of transcriptional responses triggered by O_3_ exposure that are shared, regardless of genetic background; (2) use co-expression analysis along with molecular phenotyping to identify modules of co-regulated genes and their correlations with relevant traits; and (3) identify responses that are unique to specific genetic backgrounds. With respect to the last goal, we describe responses that are both qualitatively different (*i.e.*, which genes and pathways) and quantitatively different (*i.e.*, direction and magnitude of effect), focusing on contrasts between a low- and high-responding strain (CC017/Unc and CC003/Unc, respectively).

## Materials and Methods

### Animals

All experiments described here were approved by the University of North Carolina Institutional Committee on Animal Use and Care (IACUC). Adult (10-12 weeks of age) female and male mice were used for the experiments described here. C57BL6/J mice were purchased from the Jackson Laboratory (Bar Harbor, Maine) and acclimated at the University of North Carolina at Chapel Hill for 3 weeks prior to exposure. Collaborative Cross (CC) strains CC003/Unc, CC017/Unc, CC025/GeniUnc, CC039/Unc, and CC059/TauUnc were obtained from the UNC Systems Genetics Core Facility as weanlings and aged on investigator’s racks prior to exposure. In the text, CC strain names are referred to without their corresponding laboratory code (*e.g.*, CC003/Unc is abbreviated as CC003). All animals were housed in groups of 1-5 on ALPHA-Dri bedding (Shepard) with *ad libitum* food (Envigo 2929) and water, under standard 12-hour light/dark cycles.

### Ozone exposure

Animals were exposed to filtered air (FA) or 2 ppm ozone (O_3_), as described previously (23, 41). In brief, mice were exposed in individual wire-mesh chambers without access to food or water, then returned to their home cages and allowed to recover for 21 hours before necropsy. Animals were exposed in sex- and strain-matched pairs, where one mouse was exposed to O_3_ and the other to FA (n = 3 per sex/treatment/strain except CC0039 where n = 4 females exposed to O_3_ and 1 male exposed to FA and CC003 where n = 1 female and 2 males exposed to FA). Exposures were performed across three successive days, with one matched pair per sex and strain represented in each batch.

### Phenotyping

#### Lung phenotyping

Twenty-one hours after cessation of exposure, mice were anesthetized (2 g/kg urethane) and sacrificed by inferior vena cava/abdominal aorta exsanguination. Bronchoalveolar lavage (BAL) was collected by instilling the lungs with phosphate-buffered saline containing cOmplete protease inhibitor cocktail (Roche) (0.5 mL x 1, 1 mL x 1). The BAL fluid was centrifuged for 10 minutes at 400 × g, and supernatant from the first fraction was saved and stored at −80C for biochemical analysis. Pellets from both BAL fractions were pooled, resuspended in red blood cell lysis buffer, and centrifuged once more at 400 × g for 10 minutes. Pellets were resuspended in 500 μL of HBSS. From this suspension, 400 μL was used for isolating airway macrophages, while the remainder was used for counting total numbers of viable cells and preparing cytocentrifuge slides for differential cell counts.

#### Airway macrophage (AM) isolation

The BAL cell suspension was plated with FBS-containing RPMI-1640 in a 24-well non-treated plate and left to rest in a tissue culture incubator for four hours. Non-adherent cells were removed using a vacuum line. The enriched fraction of airway macrophages was removed using Accutase (Gibco) and slow-frozen in two equal aliquots of freezing medium (90% FBS and 10% DMSO). In a separate exposure using only female C57BL/6J mice, we confirmed that this method results in an enriched fraction of AMs devoid of infiltrating monocytes and macrophages (~80%, Supplemental Figure 1).

**Figure 1.**
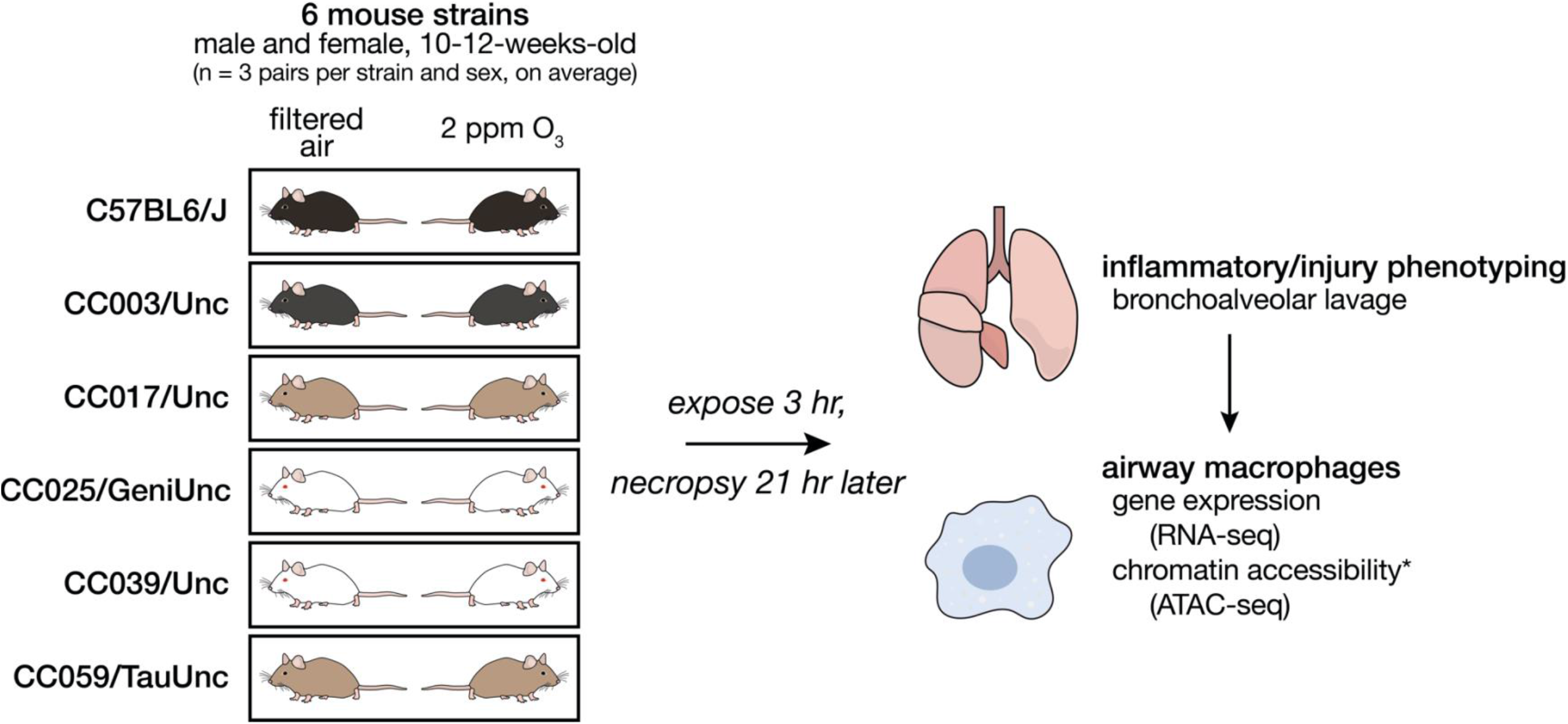
Study design. We exposed adult male and female mice from 6 strains in sex- and age-matched pairs to filtered air (FA) or 2 ppm ozone (O_3_) for 3 hours and sacrificed 21 hours later. We performed bronchoalveolar lavage (BAL) to collect cells and protein from the airspaces for quantitative inflammatory and injury phenotyping. We also collected airway macrophages by adherence from BAL cells from each mouse and performed gene expression by RNA-seq and, in CC003 and CC017, profiled chromatin accessibility using assay for transposase-accessible chromatin using sequencing (ATAC-seq). (n = 3 per sex/treatment/strain except CC0039 where n = 4 females exposed to O_3_ and 1 male exposed to FA and CC003 where n = 1 female and 2 males exposed to FA)

#### Protein measurement and cytokine profiling

Total BAL protein was quantified with the Qubit Total Protein Quantification kit and the Qubit 2.0 fluorometer (Thermo Scientific). BAL cytokines were profiled using a pre-mixed MILLIPLEX protein immunoassay (Millipore) that was read on a Bio-Plex 200 multiplex suspension array system (Bio-Rad). A panel of 15 cytokines was measured in total: eotaxin-1/CCL11, G-CSF, GM-CSF, IL-10, IL-12p70, IL-1β, IL-6, IP-10/CXCL10, KC/CXCL1, LIX, MCP-1/CCL2, MIP-1α/CCL3, MIP-1β/CCL4, MIP-2/CXCL2, and TNFα. G-CSF, GM-CSF, IL-12p70, IL-1β, MCP-1, MIP-1α, MIP-2, and TNFα were undetected in > 33% of samples and excluded from further analysis.

### Statistical analysis of phenotype data

For inflammatory cell counts, injury measurements, and cytokine profiling presented in Figures 1–3 and Supplemental Figures 2-3, raw data (Supplemental Table 1) were transformed where appropriate to resolve heteroscedasticity and ensure normality. Analysis of variance (ANOVA), likelihood-ratio and t-tests were performed in R (version 4.0.3), and results of testing were deemed significant if *p*-values were < 0.05.

**Figure 2.**
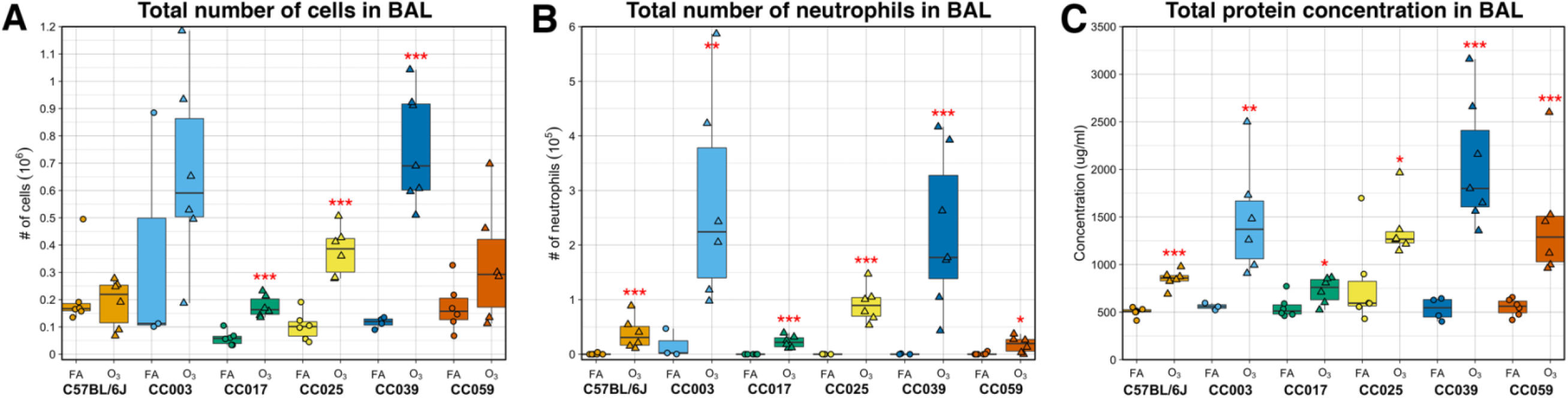
O_3_ exposure causes variable inflammation and injury across 6 strains of mice. Total cellular inflammation, neutrophilia, and total protein concentration, a metric of lung injury, were measured in the BAL fluid. (A) Total cell count (10^6^), (B) neutrophil count (10^5^), and (C) total protein concentration (μg/mL). Data are presented as box-and-whiskers plots, which display the distribution from the minimum, first quartile, median, third quartile, and maximum. Individual data points are overlaid, with circles representing FA exposed mice and triangles representing O_3_ exposed mice. All measures had significant strain-by-treatment interaction effects, assessed using a likelihood-ratio test (A: *p* < 0.0001, B: *p* < 0.05, C: *p* < 0.0001). (n = 3 per sex/treatment/strain except CC0039 where n = 4 females exposed to O_3_ and 1 male exposed to FA and CC003 where n = 1 female and 2 males exposed to FA; * *p* < 0.05, ** *p* < 0.005, *** *p* < 0.0005, for within-strain contrasts (t-tests) between FA and O_3_)

**Figure 3.**
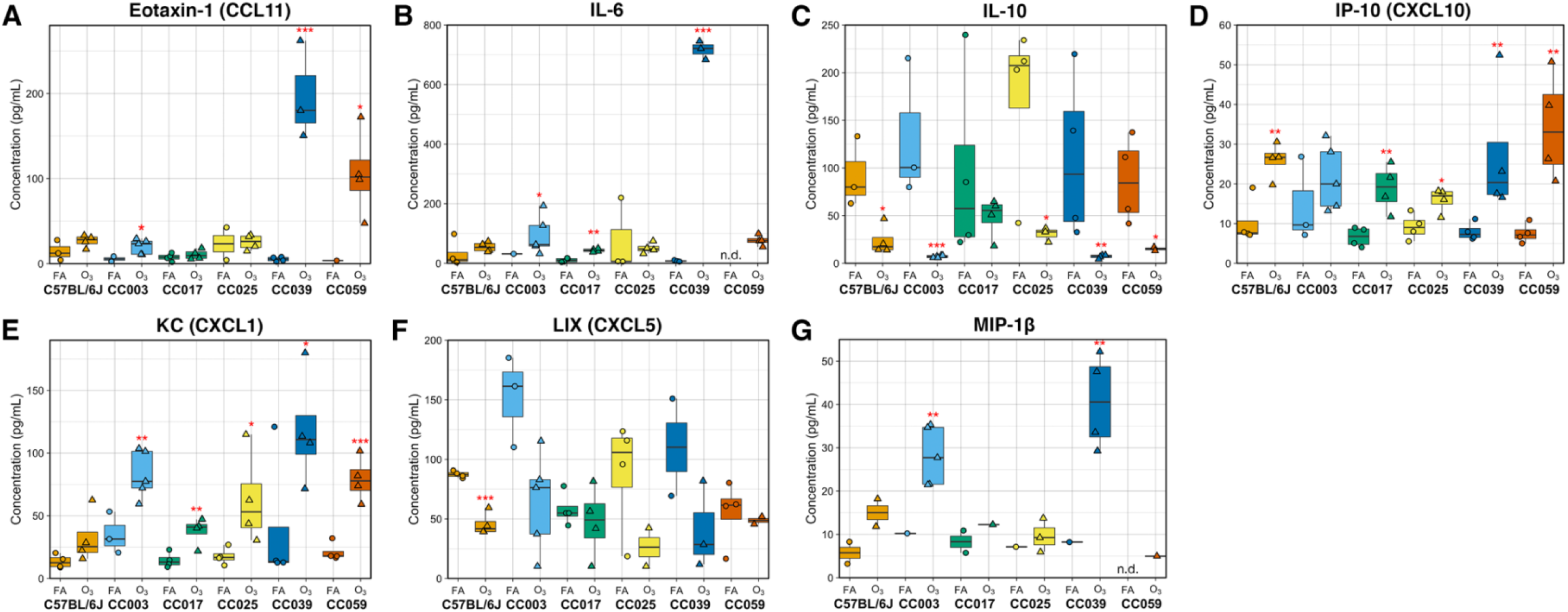
Respiratory cytokine responses are altered by O_3_ exposure and vary by strain. Molecular markers of inflammation were measured using a multiplex cytokine detection kit. (A) Eotaxin-1 (CCL11), (B) KC (CXCL1), (C) IL-6, (D) IL-10, (E) IP-10 (CXCL10), and (F) LIX (CXCL5). Data are presented as box-and-whiskers plots, which display the distribution from the minimum, first quartile, median, third quartile, and maximum. Individual data points are overlaid, with circles representing FA exposed mice and triangles representing O_3_ exposed mice. Points below the assay limit of detection were excluded from analysis. Eotaxin-1, KC, and IL-6 also displayed significant strain-by-treatment interactions assessed by a likelihood-ratio test (A: *p* < 1 x 10^−7^, B: *p* < 0.005, C: *p* < 0.0005). (n = 2 per sex/strain/treatment for all strains except CC0039 where n = 3 females and 1 male per treatment; * *p* < 0.05, ** *p* < 0.005, *** *p* < 0.0005 for within strain contrasts (t-tests) between FA vs. O_3_)

**Table 1.** Summary of differences between CC017 and CC003 in O_3_-induced phenotypes and airway macrophage molecular profiles.

|  | CC017 | CC003 |
| --- | --- | --- |
| **Inflammation (total BAL cells, neutrophils)** | low | high |
| **Injury (total BAL protein, albumin)** | low | high |
| **Cytokines** | Muted changes overall; maintained high IL-10 production post-ozone exposure and LIX was unchanged | Significant decrease in IL-10 production, large increase in production of pro-inflammatory cytokines |
| **Baseline gene expression pathways** | TGF-beta signaling; clotting and complement cascade; phagocytic functions | Glutathione metabolism; pathogen responses; canonical inflammatory signaling cascades |
| **Ozone-induced gene expression pathways** | Cell cycle progression and cellular proliferation; canonical inflammatory signaling cascades | Adipogenesis; TNF-alpha signaling via NF-kB; other canonical inflammatory signaling cascades |
| **Transcription factors associated with enriched motifs** | Lower expression of most TFs at baseline, and many remain unchanged or only minimally changed by ozone exposure | Higher expression of most TFs at baseline, and higher expression of all TFs after ozone exposure |

### RNA isolation and sequencing

One aliquot of frozen AMs was gently thawed, spun down at 400 x g for 5 minutes, and washed once with cold PBS to remove freezing medium. AM pellet was lysed directly with 350 μL of Buffer RLT, and total RNA was isolated using the Qiagen RNeasy Micro Kit. Stranded libraries were prepared using 10 ng of total RNA with the Ovation SoLo RNA-seq Library Preparation Kit (NuGEN). Four library pools were created (one matched pair per strain and treatment represented per pool) which were sequenced across four lanes on an Illumina HiSeq 4000 to generate 50-bp, single-end reads.

### ATAC-seq library preparation

Frozen AMs were thawed as for RNA isolation, and ATAC-seq libraries were prepared as described previously (42, 43) with minor modifications. AM pellets were gently resuspended in lysis buffer (10mM Tris-HCl, pH 7.5, 10 mM NaCl, 3mM MgCl_2_, 0.1% NP-40 in nuclease-free H_2_O) and nuclei were pelleted by centrifugation at 500 x g for 30 min in a refrigerated swinging bucket centrifuge. The supernatant was gently discarded, nuclei were resuspended in a 25 μL transposition reaction containing 2 μL of Tn5 transposase, and the mixture was incubated at 37°C in a thermomixer set to 1000 rpm for 1 hour. Transposed DNA fragments were purified using a Qiagen MinElute Reaction Cleanup Kit. PCR amplification was performed to add indexing primers and increase library yield. Final libraries were purified using Agencourt AMPure XP magnetic beads and quantified via Qubit dsDNA HS Assay Kit. Libraries were sequenced on an Illumina HiSeq 4000 to generate 50-bp, paired-end reads.

### Gene expression analysis

#### Sequence alignment and transcript quantification

Sequenced reads were aligned to the C57BL/6J reference genome (mm9) or Collaborative Cross strain-specific pseudogenomes (Build 37, http://csbio.unc.edu/CCstatus/index.py?run=Pseudo) using STAR (44) and quantified using RSEM (45).

#### Differential expression analysis

To estimate the treatment effects, strain effects, and strain-by-treatment interactive effects on gene expression, a series of models were fit using DESeq2 (46).

The effect of treatment of gene expression was estimated using the following model,

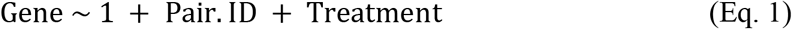

where Pair. ID is a factor that indicates sex- and strain-matched pairs, to accommodate the experimental design. Treatment coefficients representing the “T effect,” were evaluated within the DESeq2 framework using the Wald test, with false-discovery rate (FDR) correction (α = 0.05).

The influence of strain on gene expression was assessed outside of the paired framework used in Equation 1, with the following model,

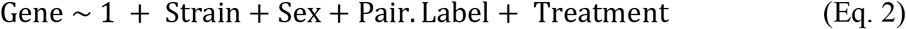

where Pair. Label describes batch. Strain coefficients representing the “G effect” were tested for significance using the likelihood-ratio test, against a reduced model less the Strain term, subject to FDR correction (α = 0.05).

Finally, “GxT effect” coefficients were fit in the paired framework using an extension of Eq. 1,

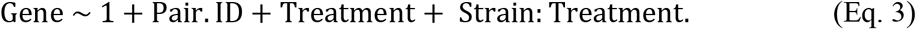

where Strain: Treatment coefficients were tested for significance using the likelihood-ratio test, using the following process. For each gene, a set of six GxT coefficients was computed: the first was based on the differential expression for a given strain compared to the mean differential expression across the remaining five strains, and the other five were determined by repeating this procedure individually comparing against each of the other strains. This process was applied for all genes, generating a matrix of GxT coefficients. All GxT coefficients were tested for significance using the Wald test, and subject to joint shrinkage using r/ashr (47) and FDR correction (α = 0.05).

#### Weighted gene co-expression network analysis (WGCNA)

Network co-expression analysis was performed using r/WGNCA (48). Expression data for all genes with number of reads greater than total number of samples (N = 46) were included (13,594 genes) and data were subjected to the variance-stabilizing transformation. One consensus network was generated for samples from both FA and O_3_-exposed mice using the recommended WGCNA approach for automatic network construction and module detection. We assumed a signed correlation network and used biweight midcorrelations to compute the correlations. We selected a soft threshold of twelve based on a 0.9 threshold for the scale free topology index. For settings related to dendrogram cutting and module merging, we used the recommendations in the tutorial (49). After module assignment, we examined the correlation of the eigengene (first principal component) for each of these modules with inflammatory, injury, and cytokine metrics measured across the panel of strains. Modules were then prioritized based on the number and significance of these correlations, and gene set enrichment was performed.

#### Gene set enrichment

Lists of genes were examined for putative biological significance using r/enrichR (50), with databases listed in Supplemental Table 2.

### ATAC-seq analysis

AM ATAC-seq data from CC003 and CC017 mice were analyzed with the following methods and procedures.

#### Sequence alignment and pre-processing

The PEPATAC (51) pipeline with default parameters was used to perform adapter trimming, alignment, and peak calling for ATAC-seq reads, which were aligned to the appropriate pseudogenome. Uniquely mapped reads were converted to mm9 coordinates (excluding those overlapping the mm9 blocklist) to allow for cross-strain comparisons. The union set of peaks were called from each sample, and divided into 300-bp overlapping windows which were further subdivided based on their location (proximal: within 5 kb of nearest transcription start site (TSS); distal: beyond 5 kb of nearest TSS).

#### Differential accessibility analysis

Similar models were used to perform differential accessibility analysis as described for differential expression analysis above, where Gene is replaced with number of mapped reads within 300-bp overlapping windows, with a covariate included to account for differences in data quality. Following differential accessibility analysis, significant neighboring windows within 200 bp were merged (FDR < 0.05). Motif analysis was performed using HOMER (52), focusing on enrichment of known motifs.

## Results

### Cellular and molecular markers of O_3_-induced inflammation and injury vary by strain

To investigate the influence of genetic background on O_3_-induced respiratory responses, we exposed adult, female and male mice from 1 classical inbred and 5 Collaborative Cross (CC) strains in matched pairs to filtered air (FA) or 2 ppm O_3_ for 3 hours (n = 3 per sex/treatment/strain except CC039 where n = 4 females exposed to O_3_ and 1 male exposed to FA and CC003 where n = 1 female and 2 males exposed to FA; Figure 1). Relative to FA-exposed control mice, O_3_ exposure caused a significant increase in the total number of bronchoalveolar lavage (BAL) cells in three of six strains, CC017, CC025, and CC039 (Figure 2A). Across all strains, neutrophils were significantly increased in mice exposed to O_3_ relative to their FA counterparts (Figure 2B). Three-way analysis of variance (ANOVA) with strain, treatment, and sex as factors revealed a significant main effect of strain and treatment. Sex was not a significant predictor for either cellular measurement. We used a likelihood-ratio test (LRT) to compare model fit when including an interaction term to the main effects model, which indicated significant strain-by-treatment interactions for both total cells and neutrophils (*p* < 0.0001 and *p* < 0.01, respectively). Similar effects were seen when considering proportion of neutrophilic infiltrate (*i.e.*, percentage of neutrophils) in the BAL (Supplemental Figure 2). CC003 and CC039 were the highest responders as judged by the total number and proportion of neutrophils in the BAL, while CC017 had the lowest O_3_-induced inflammation and neutrophilia.

In addition to cellular inflammation, we assessed lung injury induced by acute O_3_ exposure, as reflected by the concentration of protein in BAL. After FA exposure, all strains had low levels of total protein in the lung lavage fluid (Figure 2C); however, the increased levels of BAL protein varied by strain after O_3_ exposure. As with inflammation, we detected a significant main effect of strain and treatment, with no significant effect of sex on total protein concentration. Similar to the results for inflammation, there was a highly significant strain-by-treatment interaction (*p* < 0.0001). Because total protein is a non-specific marker of serum protein leakage and lung injury, we measured serum albumin by ELISA in a subset of these mice to confirm our findings. Overall, the strain order of the post-O_3_ exposure effect was similar between total protein and serum albumin measures of lung injury; however, in the FA samples, there was greater variability between strains revealed by the albumin assay (Supplemental Figure 3). Interestingly, the rank order of strain effects was not strictly the same when comparing BAL cellular inflammation and total protein, though CC003 and CC039 remained high responders while CC017 had the lowest increase in total protein concentration after O_3_ exposure of the strains surveyed.

We also measured a panel of hallmark O_3_-responsive cytokines in the BAL to further characterize the molecular aspects of airway inflammatory responses (Figure 3). One strain, CC039, had markedly high concentrations of eotaxin-1 (CCL11) and IL-6 in response to O_3_ exposure in comparison to the rest of the strains (Figure 3A & B). The chemokine IP-10 (CXCL10) was highly upregulated in O_3_-exposed animals in 5 of the 6 strains, but was only slightly altered in CC003 (Figure 3D). As we observed in a previous study (23), the anti-inflammatory cytokine IL-10 was decreased upon O_3_ exposure in a strain-dependent manner (Figure 3D). Similarly, levels of LIX (CXCL5) tended to be lower in O_3_-exposed mice compared to FA controls, though this effect was only significant in C57BL/6J. Notably, for nearly all cellular and biochemical phenotypes measured, CC strains displayed responses that were both more and less pronounced than the classical inbred strain used (C57BL/6J).

Finally, we calculated pairwise correlations between these phenotypic traits (Supplemental Figure 4). As expected, levels of the anti-inflammatory cytokine IL-10 were negatively correlated with most other inflammatory phenotypes excluding concentrations of LIX, for which there was a positive correlation. Otherwise, nearly all other phenotypes were significantly positively correlated with one another, with correlation coefficients in the range of 0.48-0.85. Some traits were more strongly correlated, including IL-6 and eotaxin-1/CCL11 concentrations, while others were less closely linked. These set of correlations, along with the shuffling of strain order of effect when comparing cellular inflammation to total protein concentration, suggest that responses to O_3_ exposure do not follow a simple linear relationship and are subject to different types of genetic regulation.

### A core set of O_3_-responsive transcripts are shared across the panel of strains, and group into co-expression modules correlated with relevant phenotypes

To comprehensively characterize the gene expression alterations induced by O_3_ exposure in an important target cell population, we performed RNA-seq analysis in airway macrophages (AM) isolated from a subset of samples from each of the strains (n = 2 per sex, treatment, and strain). Principal components analysis (PCA) of the top 500 most variably expressed genes revealed that samples clearly separate by strain (PC1) and treatment (PC2), and these PCs account for roughly 33% of variance within the data (Figure 4). Intriguingly, samples only begin to separate by sex at PC9, which accounts for a mere 2.62% of total variance within this subset of genes, suggesting that gene expression in this context is modestly influenced by sex (Supplemental Figure 5).

**Figure 4.**
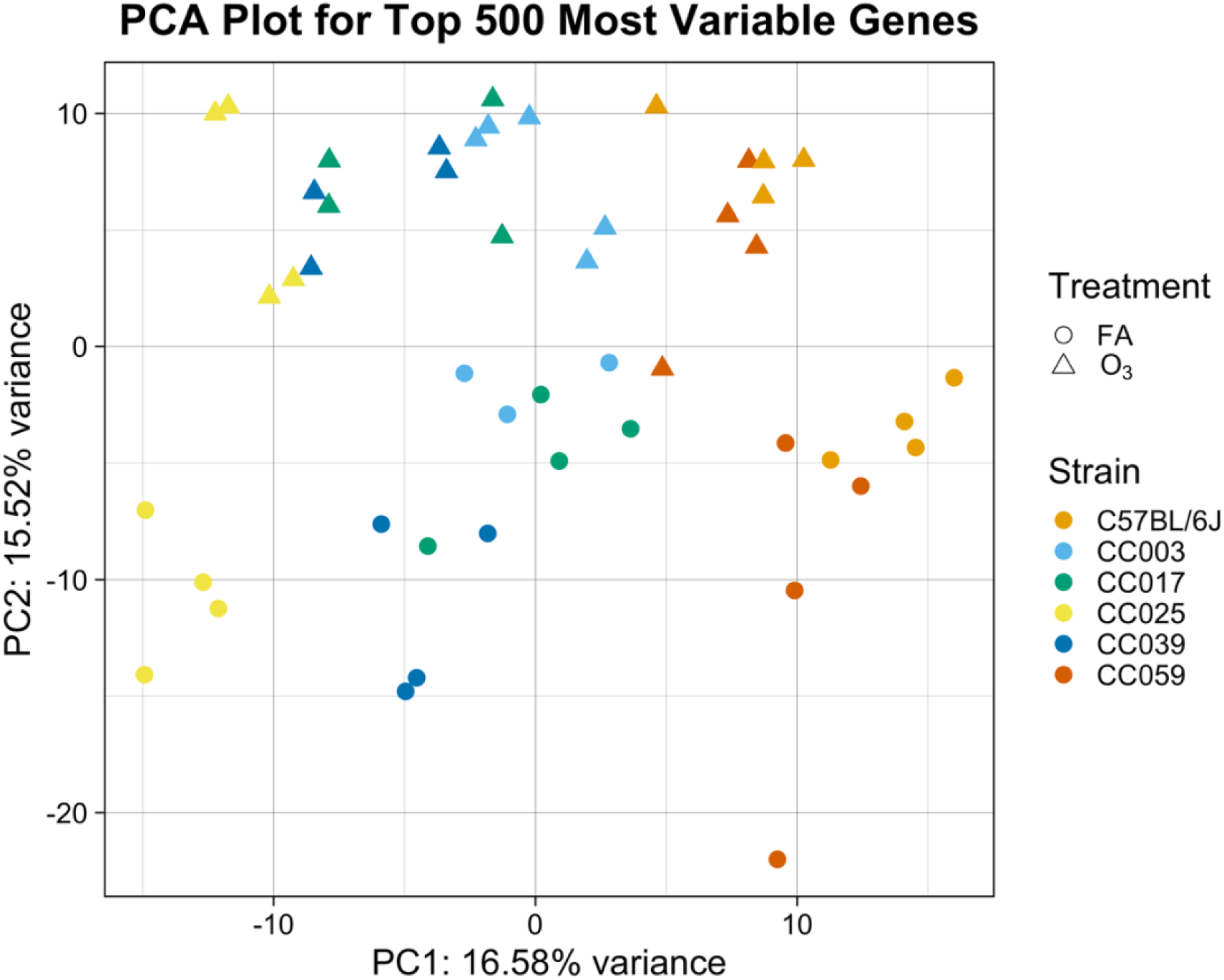
Airway macrophage (AM) gene expression varies by strain and O_3_ exposure. RNA-seq was performed on a subset of mice exposed to FA or 2 ppm O_3_. Principal components analysis (PCA) of the top 500 most variably expressed genes reveals separation of samples by strain (PC1) and treatment group (PC2). (n = 2 per sex/strain/treatment, except CC003 where n = 1 female and 2 males per treatment and CC039 where n = 3 females and 1 male per treatment)

We were first interested in identifying genes whose expression is altered by O_3_ exposure (treatment effect or “T effect”), without explicit consideration of genetic background. To define the set of T effect transcripts, we used a paired framework within DESeq2 to regress out the effects of strain and batch, and to extract the treatment effect. We identified 2,761 O_3_-responsive genes with an FDR < 0.05, 773 of which had an absolute log_2_ fold change > 1. These results are depicted in Figure 5, and all significantly differentially expressed genes (DEGs) are reported in Supplemental Table 3. Four DEGs that we previously identified in a meta-analysis, *Thbs1*, *Lcn2*, *S100a9*, and *Cdk1*, exhibited sizable changes (log_2_(fold change): ~1.6 - 5.5) in gene expression (23). Other significant DEGs included *Scgb1a1*, *Cyp1b1*, and *Mmp9*. *Scgb1a1* is predominantly produced by club cells but was recently implicated in alveolar macrophage immune responses (55) and has been proposed as a biomarker of O_3_-induced lung injury in humans (56), while *Cyp1b1* is a well-known detoxification enzyme regulated by activity of the aryl hydrocarbon receptor and *Mmp9* is an extensively characterized matrix metalloprotease associated with proinflammatory macrophage activity (57, 58).

**Figure 5.**
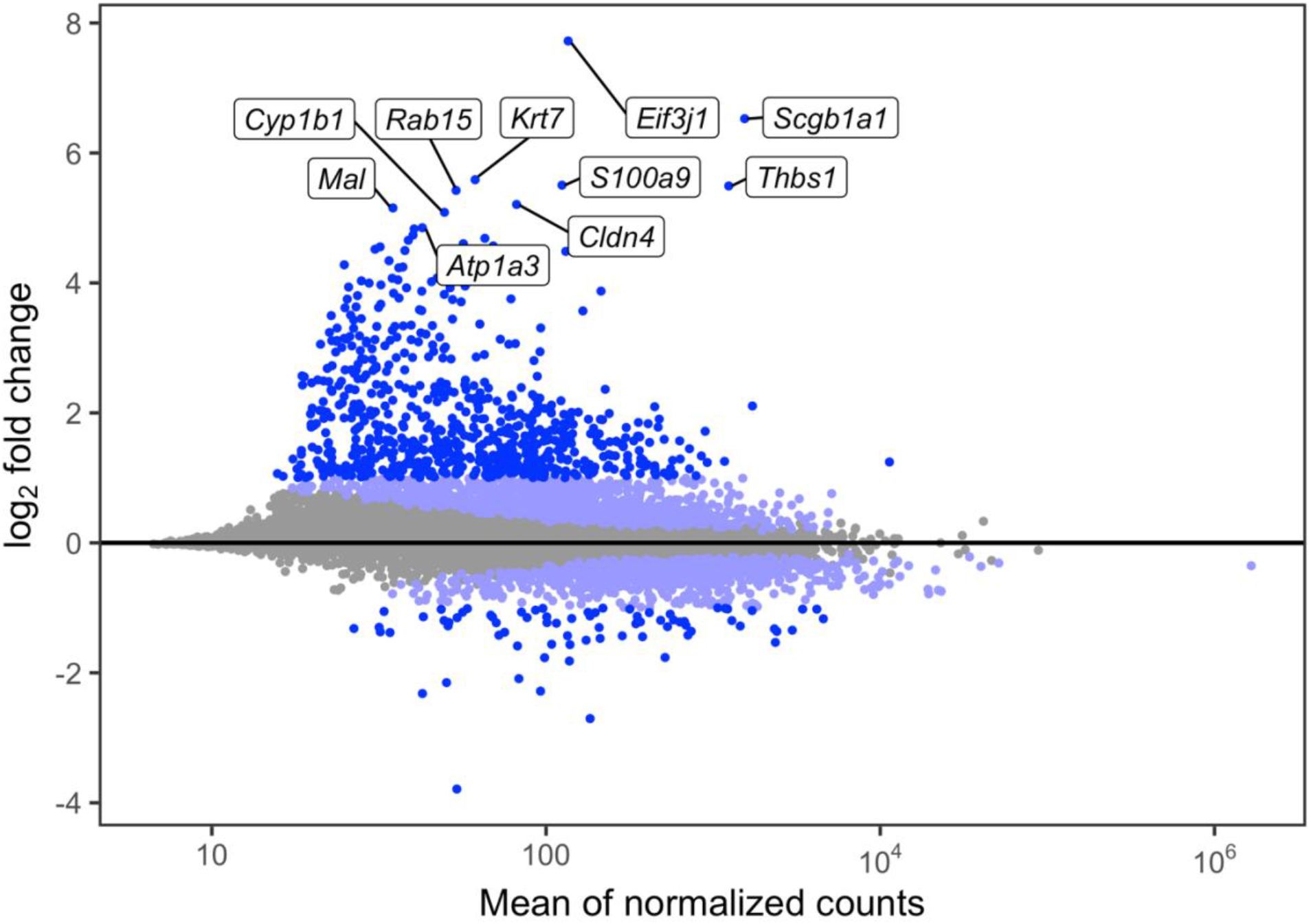
O_3_ exposure causes characteristic changes in airway macrophage (AM) gene expression. The union set of the light blue and royal blue dots are all significantly differentially expressed genes (2,761; FDR < 0.05). The royal blue dots alone represent those with absolute log_2_ fold change > 1 (773). The top 10 most highly differentially expressed genes are labeled.

Next, we were interested in relating our gene expression data to the phenotypic data collected on the same mice to identify networks and pathways associated with relevant traits. Thus, we performed weighted gene co-expression network analysis (WGCNA; (48)) to jointly cluster the expression data from both FA- and O_3_-exposed mice into co-expression modules. Using this expression set, we grouped 4,373 genes into 18 modules containing a variable number of genes, ranging from 51 to 1,462 (Supplemental Table 3). The remaining 9,221 transcripts were collected in an unassigned module (“grey60”).

Five of the 18 modules displayed strong module-trait relationships that can be grouped into two broad categories: three modules (“red,” “brown,” “blue”) that are positively correlated with pro-inflammatory cytokine secretion, lung injury, and neutrophilia and the remaining two modules (“pink,” “midnightblue”) exhibiting the inverse relationship with those traits (Supplemental Figure 6). For these five modules, we examined pathway annotations associated with the assigned lists of genes (Supplemental Table 4). The largest of the five modules (“blue,” 742 genes) was positively associated with pro-inflammatory cytokine secretion, and enriched for pathways involved in cell cycle progression, DNA replication and repair. Another large module within the first category (“brown,” 352 genes) was enriched for several pathways including those generally associated with inflammation such as signaling induced by canonical regulators (*e.g.*, TNFα, IL-6, TGF-β) and responses to immune stimuli like lipopolysaccharide (LPS), salmonella, and influenza A. The genes within this module were also enriched for terms associated with protein modification and processing in the endoplasmic reticulum. The final module within this category (“red,” 188 genes) was enriched for pathways associated with oxidative phosphorylation and other mitochondrial metabolic processes, as well as fatty acid metabolism and thermogenesis. A small module falling within the second category (“pink,” 131 genes) only displayed biological associations with one term (“focal adhesion”), though we discovered many genes involved in lipid processing and metabolism after manual inspection of this gene set including *Alox5ap* (5-lipoxygenase-activating protein), *Fabp1* (fatty acid-binding protein 1), *Adipor1* (adiponectin receptor 1), and *Abcg1* (ATP-binding cassette sub-family G member 1). Finally, the second module displaying inverse correlation with pro-inflammatory phenotypes (“midnightblue,” 66 genes) was associated with several RNA polymerase II-mediated processes and a few inflammatory signaling pathways including TNFα signaling via NF-κB.

### Strain background modifies basal and O_3_-induced AM gene expression

After establishing a set of O_3_-responsive transcripts that were differentially expressed across all strains, we sought to explore how genetic background influences transcriptomic responses to O_3_ exposure, and examine whether the transcriptome could give insights into biological signatures that distinguish low- and high-responding strains. Specifically, we used a series of modeling approaches to identify genes whose expression is influenced by strain background (“G effect”) and genes for which there are interactive strain-by-treatment effects on expression (“GxT effect”).

First, we identified genes with significant G effects, accounting for batch, sex, and treatment (Supplemental Table 3). Overall, there were 3,787 genes (28% of examined transcripts) for which strain was significantly associated with expression (FDR < 0.05), a level that is comparable to a previously published study examining the influence of genetic and environmental effects on peritoneal macrophage gene expression using the Hybrid Mouse Diversity Panel (5,726 out of 12,980 measured; 44.1%) (17). Additionally, this figure is slightly greater than the total number of T effect genes (2,761) defined in the previous section. Next, we identified genes with significant GxT effects (Supplemental Table 3). Rather than testing for interactive effects between strain and treatment jointly across all strains or by examining all 15 possible pairwise strain contrasts, we estimated GxT effects for each gene by comparing differential expression (O_3_ vs. FA) in one strain against the mean differential expression of all other strains (*i.e.*, each coefficient is the strain-specific relative response for a given gene). This procedure was performed iteratively for all genes across all strains, resulting in a 13,594 x 6 matrix of coefficients. In total, 545 genes (4% of examined transcripts) had one or more significant GxT effect (FDR < 0.05). Within this list were numerous cytokine genes (*Ccl2*, *Ccl6*, *Ccl7*, *Ccl9*, *Cxcl3*, *Il18*, *Slpi*) and scavenger receptors (*Cd36*, *Colec12*, *Marco*, *Msr1*), classes of genes that have been previously implicated in O_3_ responses (59–64) (Figure 6). Interestingly, there were also 245 genes with both significant G and GxT effects on gene expression, including the genes visualized in Figure 6.

**Figure 6.**
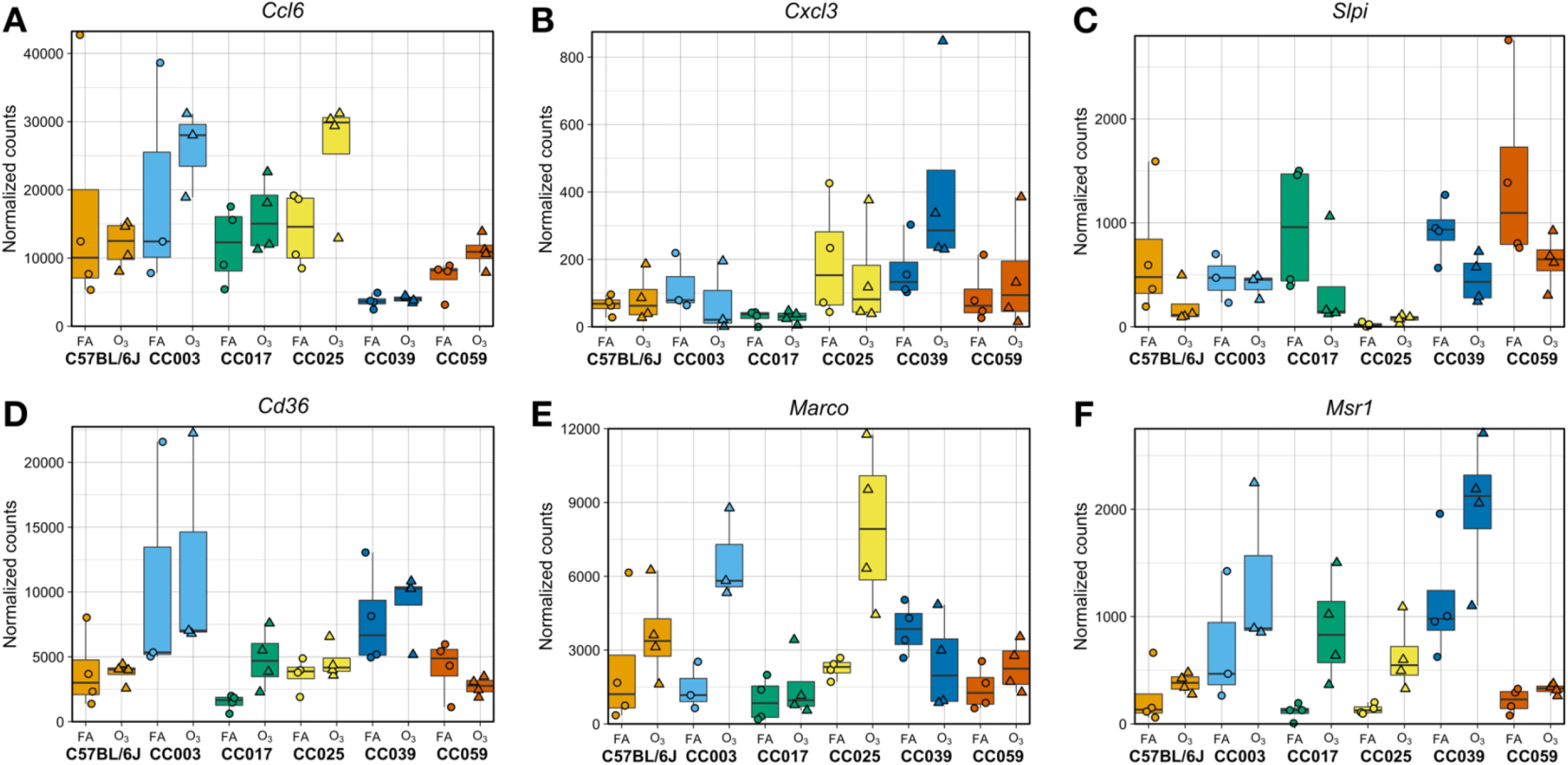
Numerous genes display strain-by-treatment (“GxT”) effects on expression, including those with putative and known roles in O_3_ responses. (A-C) represent immune response genes, while (D-F) are scavenger receptors. Some genes display GxT effects across all strains (*e.g.*, *Marco*, *Msr1*), while others are selectively observed in only some strains (*e.g.*, *Ccl6*, *Cxcl3*, *Slpi*). (n = 2 per treatment/sex/strain; all genes have one or more significant GxT effects on expression [FDR < 0.05])

To identify general patterns of response and guide the choice of specific strains on which to focus downstream analyses, we visualized the matrix of GxT coefficients using a PCA biplot, where each gene is an observation (point) and each strain is a loading vector (line) (Figure 7). Genes clustered along these loading vectors contribute most to the corresponding strain’s overall gene expression response, where positive coefficients indicate that the strain had higher relative change in expression induced by O_3_ exposure than the others, and vice versa for negative coefficients. Strikingly, in alignment with their divergent inflammatory and injury phenotypes, loadings for CC017, CC003, and CC039 were highest among the six strains. CC017 (a low responder) is nearly orthogonal to both CC003 and CC039 (high responders), while the latter two strains are in opposing directions. Within the set of coefficients for CC017, *Slc40a1*, *Plxdc2*, and *Pla2g7* were among genes with greatest positive contributions to this strain’s loading while *Vcan* and *Cd74* contributed negatively. The former group of genes are known to be involved in macrophage behaviors including iron homeostasis, IL-10 production via pigment epithelium-derived factor (PEDF), and suppression of pro-inflammatory platelet factors (65–67), and the latter set are associated with leukocyte adhesion/migration and neutrophil recruitment (68, 69). Included in CC003’s set of genes with positive coefficients were *Card11, Rxra*, and *Lrch4*, though many more genes with negative coefficients contributed to its loading including known anti-inflammatory markers associated with M2 macrophage polarization such as *Hif1a*, *Mmp12*, and *Clec4a2*. Finally, the important immune-related genes *Ccl9*, *Arg1*, and *Cd86* contributed positively to CC039’s loading, while *Marco* and *Osbpl3* were in the group of genes negatively influencing this strain’s loading. *Marco* is a scavenger receptor whose expression is displayed in Figure 6E, while *Osbpl3* is a member of the oxysterol-binding protein family; both may be involved in regulating clearance of O_3_-generated reaction products (59, 70, 71).

**Figure 7.**
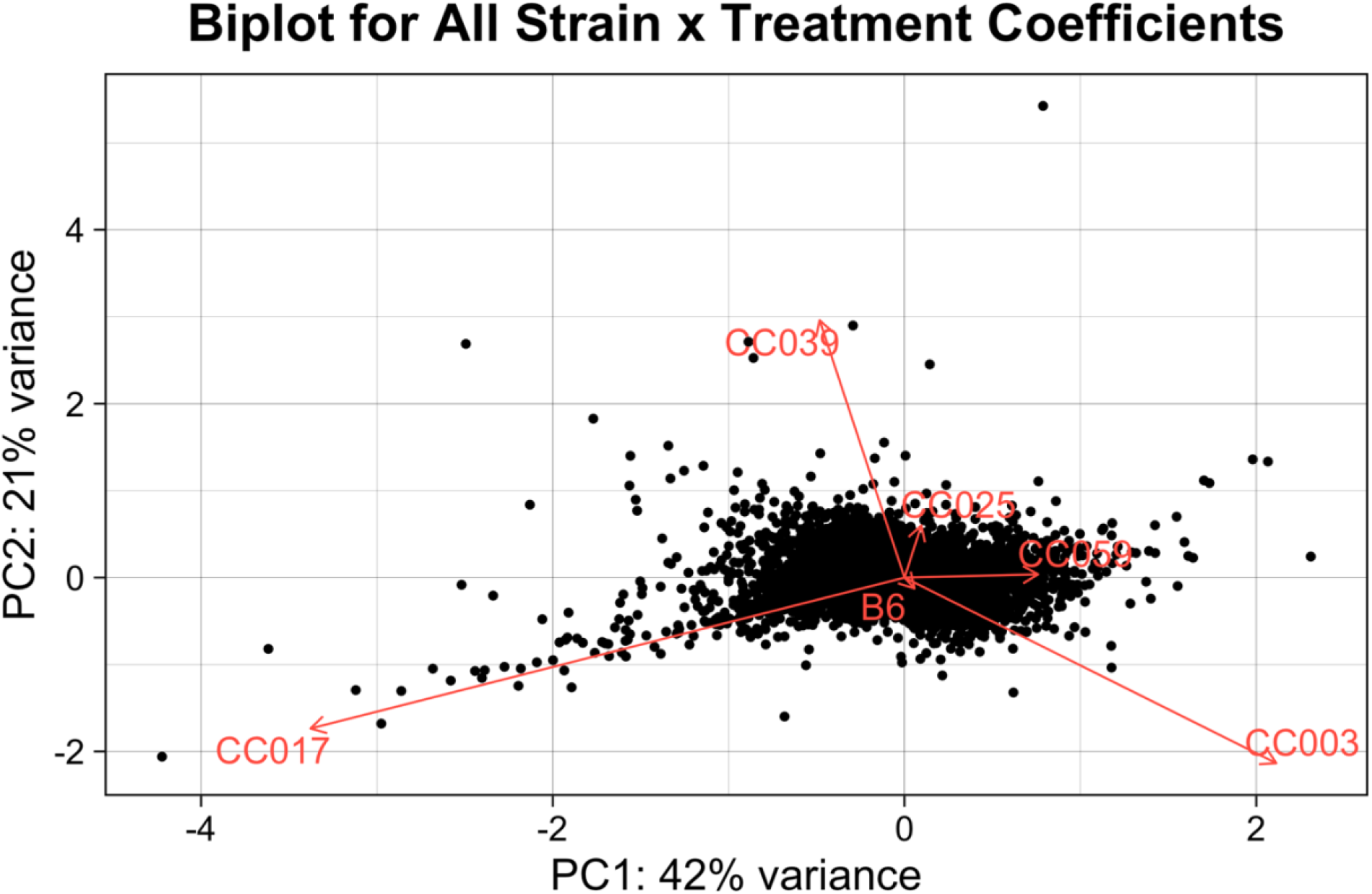
Strain-by-treatment (GxT) coefficients for gene expression separate strains by their gene expression responses. PCA was performed with the matrix of GxT coefficients. Each point on the graph represents the first two principal components for a single gene (that has six coefficients, one per strain), where the loadings represent the characteristic for each strain.

Because CC003, CC017, and CC039 displayed such distinct responses compared to the other strains, we sought to identify GxT effects between these pairwise contrasts. Comparing genes whose expression was differentially altered by O_3_ exposure in CC017 versus CC003, we identified 1,091 significant GxT DEGs, while in CC039 versus CC017, only 148 genes were significantly DE, and in CC039 versus CC003, 373 genes were significantly DE (FDR < 0.05). Because the first contrast (CC017 versus CC003) was most marked with both the largest number of DEGs identified and the inclusion of a low and high responding strain (in terms of BAL neutrophilia and total protein concentration), we chose to perform further analysis with this pair of strains.

### CC003 displays enhanced induction of pro-inflammatory pathways compared to CC017, which may be mediated by divergent basal gene expression profiles

The GxT genes defined in the previous section represent one potential explanation for why CC003 and CC017 face divergent phenotypic consequences after O_3_ exposure. Gene-environment interactions in gene expression can manifest through multiple modes, which can be visualized as reaction norms (Figure 8); these response patterns have been explored previously in great detail (17, 72). The classical example arises when two genotypes express similar levels of transcript in one condition and diverge upon introduction to a new condition, (*i.e.*, differences are revealed by changing exposures; Figure 8C). Alternatively, two genotypes may have different expression at baseline (FA controls) and further exhibit distinct degrees (slopes) of change in response to exposure (Figure 8D-F).

**Figure 8.**
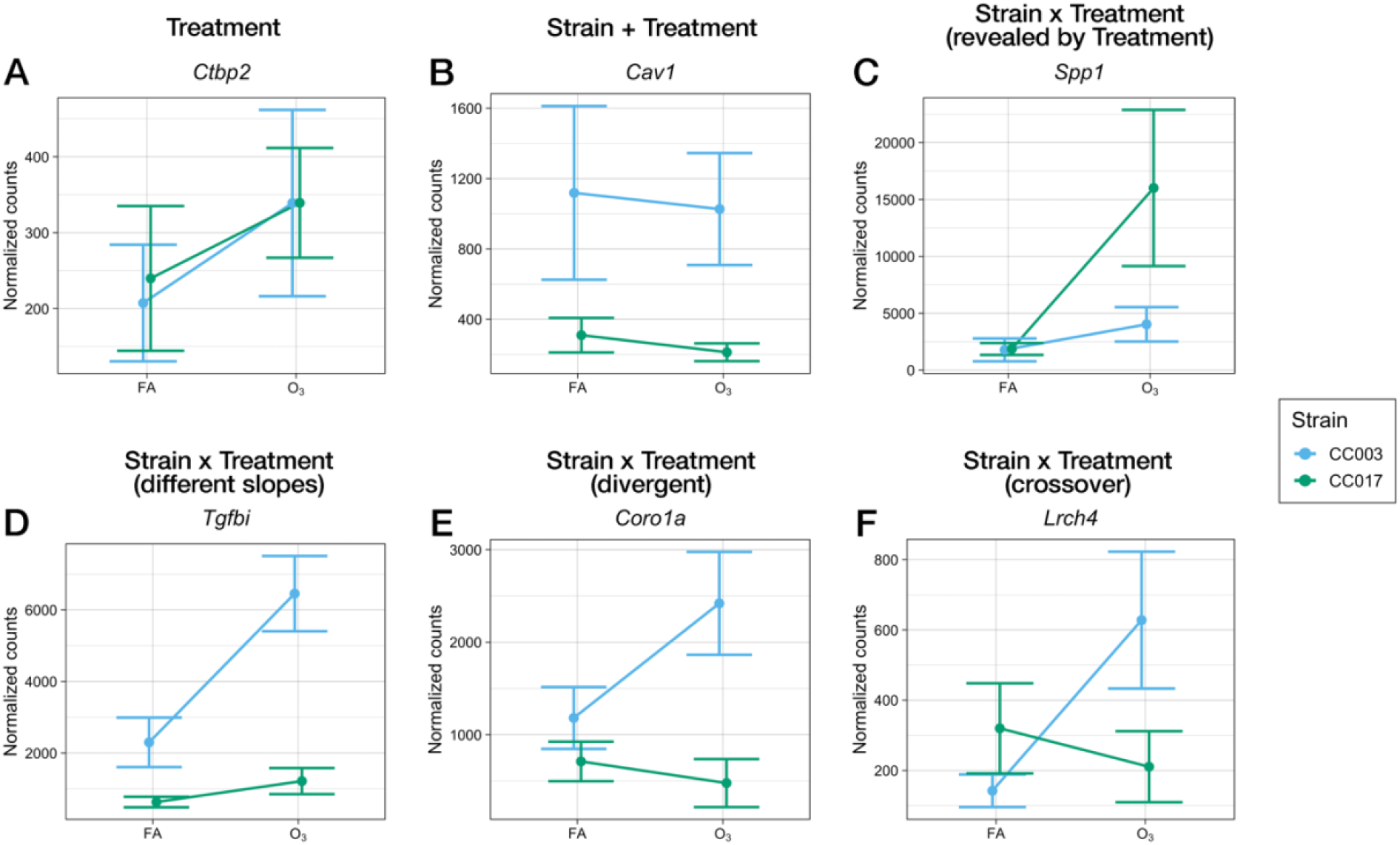
Examples of effects of treatment and strain on gene expression between CC003 and CC017, illustrated by reaction norms. In each plot, a gene is visualized that depicts the labeled effect on gene expression. (A) Treatment effects are seen for *Ctbp2* (C-terminal-binding protein 2), where both strains have increased expression after O_3_ exposure but not significantly different expression from one another in either condition. (B) Additive (but not interactive, noting the parallel slopes) strain and treatment effects are seen, where CC003 expresses *Cav1* (caveolin-1) more highly at baseline than CC017, which is retained after O_3_ exposure. (C) Strain-by-treatment interactive effects are seen for *Spp1* (osteopontin), where both strains express similar levels at baseline and CC017 expresses much greater levels after O_3_ exposure than CC003. (D) Baseline expression of *Tgfbi* (transforming growth factor-beta-induced) differs between CC003 and CC017 and increases in both strains after O_3_ exposure, with greater magnitude in CC003 (i.e.*, along different slopes, but in the same direction)*. (E) Baseline expression of *Coro1a* (coronin 1A) differs between CC003 and CC017 and decreases in CC017 after O_3_ exposure while increasing in CC003. (F) Expression of *Lrch4* (leucine-rich repeat and calponin homology domain-containing protein 4) is higher in CC017 than in CC003 at baseline, but increases with O_3_ exposure in CC003 to surpass CC0017’s original level. CC017 expression of this gene is decreased, to a similar level seen in CC003 at baseline.

To explore the biological functions associated with genes DE between CC017 and CC003 and characterize how their responses to O_3_ differ, we performed pathway analysis with the set of of 1,091 GxT DEGs described above (partitioned by whether O_3_-induced expression was enhanced in CC017 versus CC003; Supplemental Tables 5 & 6). Within the set of 806 genes with greater O_3_-induced upregulation in CC017 compared to CC003 (764 with log_2_ fold change > 1, CC017 versus CC003), pathways associated with cell cycle-related targets of E2F (*Ctcf*, *Brca1*, *Brca2*, *Dck*) and the inflammatory response (*Il18*, *Hif1a*, *Klf6*, *Ccl2*, *Ccl7*, *Il7r*) were overrepresented. Conversely, for the 285 genes with greater expression after O_3_ exposure in CC003 compared to CC017 (256 with log_2_ fold change < −1, CC017 versus CC003), pathways involved in adipogenesis (*Stat5a*, *Apoe*), IL-2/-5/-7/-9 signaling (*Map2k2*, *Cdk4*, *Ptpn6*, *Eif3b*, *E2f1*), and TNFα via NF-κB (*Irs2*, *Junb*) were enriched.

An alternative, and not mutually exclusive, explanation is that basal gene expression (*i.e.*, in FA-exposed mice) is a determinant of the contrasting responses between CC003 vs CC017, which we have observed in other studies (73). These baseline differences can result in both additive changes in gene expression (*i.e.,* different intercepts, but the same slope; Figure 8B), as well as genotype-environment interactions in expression (Figure 8D-F). We evaluated baseline differences by examining genes with significant G effect (extracted from the modeling framework used to identify genes with significant GxT effects; here termed G' effect). Between CC017 and CC003, 1,187 genes with significant G' effects were identified, 386 of which were more highly expressed in CC017 and 801 more highly expressed in CC003 (Supplemental Table 7). Next, we performed pathway analysis, to gain insight into the types of biological activities implicated by these 1187 genes*, i.e.*, pathways that that differ between the strains at baseline (Supplemental Table 8). Pathways enriched within the set of G' effect genes more highly expressed in CC017 at baseline included those associated with TGF-β signaling (*Serpine1*, *Furin*, *Klf10*), complement and coagulation/clotting cascade (*Plau, Vwf*, *F7*, *F10*, *Cfb*), and phagocytic pathways (*Tlr2*, *Itgb5*, *Scarb1*). Conversely, pathways associated with genes more highly expressed at baseline in CC003 included glutathione metabolism (*Gstm1*, *Anpep*, *Mgst2*), responses to a variety of pathogens (*e.g.*, influenza A, *Bordetella pertussis*, *Legionella*, and *Salmonella*), and numerous canonical inflammatory pathways (*e.g.*, signaling via IL-2/STAT5, IL-17, IL-6, and Toll-like receptors). Additionally, a few pathways were enriched in both sets of genes, including TNFα signaling via NF-κB, xenobiotic metabolism, and oxidative stress response. Taken together, these results suggest that higher expression of canonical inflammatory pathways in CC003 mice prime or accelerate their responses to O_3_ compared to CC017.

### AM chromatin accessibility is influenced by strain and strain-by-treatment, but not treatment effects, when comparing CC0017 and CC003

One prominent mechanism by which genetic variation alters gene expression is through modifying the degree of chromatin accessibility indicative of changes to regulatory activity and organization of the transcriptional landscape. Hence, we were interested in examining chromatin state differences between CC003 and CC017 using the same statistical analysis frameworks we used for gene expression. To achieve this goal, we generated genome-wide chromatin accessibility profiles using ATAC-seq for a subset of these mice (n = 2 FA- and 3 O_3_-exposed per strain) and generated lists of differentially accessible regions (DARs) that were significantly influenced by strain, treatment, and strain-by-treatment interaction. We identified 2,030 DARs with significant G effects (FDR < 0.05), 963 of which were more accessible in CC017 and 1067 of which more accessible in CC003. In contrast to the strong effects of O_3_ exposure on transcriptional responses in these strains, we identified no DARs with significant T effects. Finally, we identified 414 regions with significant GxT effects on accessibility (FDR < 0.05), 191 of which were more accessible in response to O_3_ in CC017 and 223 more accessible in response to O_3_ in CC003. Overall, these results imply that the baseline chromatin landscape between the two strains is more divergent than that which is induced by O_3_ exposure.

We examined both sets of DARs (G and GxT) for enrichment of transcription factor motifs using HOMER (52), subset by their position relative to transcription start sites (proximal: < 5 kb; distal: > 5 kb) and their relative accessibility (more accessible in CC017 or CC003). This analysis revealed motifs for transcription factors in the AP-1/ATF family including AP-1, FRA1, FRA2, BATF, and JunB, enriched in distal CC017-accessible sites, both in the set of G and GxT effect DARs. Many of these motifs were also enriched in distal CC003-accessible sites for both G and GxT effect DARs. We also inspected gene expression of these transcription factors to examine whether they exhibited differential expression between the strains at baseline and/or after O_3_ exposure (Supplemental Figure 7). We discovered that *Fos* had a significant G’ effect (Supplemental Figure 7A; adj. *p* = 1.1 × 10^−4^), while *Jun* had suggestive influence of a GxT effect (Supplemental Figure 7B; adj. *p* = 0.11) and *Junb* had significant influence of both G’ (adj. *p* = 2.4 × 10^−2^) and GxT effects (adj. *p* = 7.7 × 10^−3^; Supplemental Figure 7C). Expression of *Fosl1,* the gene encoding FRA1, was also subject to significant G' effect (Supplemental Figure 7D; adj. *p* = 1.8 × 10^−2^), though its overall expression was much lower than that of the other factors. In all cases, expression of the given factor was higher in CC003. Distal regions that were more accessible in CC003 also contained a preponderance of motifs for ETS family transcription factors such as ELF3, ELF4, ELF5, and PU.1. Upon examining expression of the gene encoding PU.1 (*Spi1*), we found near-significant (adj. *p* = 0.056) GxT effect on expression between CC017 and CC003 (Supplemental Figure 7E).

Proximal CC003-accessible regions were also enriched for ETS factor motifs in both the set of G and GxT effect DARs, though the latter results were distinguished by the presence of many IRF-family factors including IRF1, IRF3, and IRF8. Expression of *Irf8* was significantly influenced by G' effect (adj. *p* = 4.4 x 10^−6^), along with modest GxT effect (adj. *p* = 0.073), while the remaining IRF factors were not influenced by strain or treatment effects (Supplemental Figure 7F). Finally, the lists of enriched motifs for proximal CC017-accessible regions were highly distinct between the G effect and GxT effect DARs. In the former set, numerous zinc-finger proteins were enriched in the list of predicted transcription factors, including some with highly similar binding motifs (*e.g.*, Sp1, Sp2, KLF4, KLF6). Within this set of factors, we determined that *Klf6* expression was influenced by significant G' (adj. *p* = 4.5 x 10^−4^) and GxT effects (adj. *p* = 4.1 × 10^−3^; Supplemental Figure 7G). In the latter set, there was a wide diversity of predicted motifs represented including for many transcription factors present in the other sets of DARs, as well as for NRF1 (nuclear respiratory factor 1) and BHLHE41, which have important roles in regulating cellular growth and renewal (74), and CEBPB which an important macrophage transcription factor (75, 76). *Cebpb* gene expression was subject to both G’ (adj. *p* = 0.019) and GxT effects (adj. *p* = 0.025; Supplemental Figure 7H).

Together, these results indicate that CC003’s enhanced responsiveness to O_3_ exposure may be mediated by heightened pro-inflammatory AM transcriptional activity, which was characterized by increased enrichment of canonical immune signaling pathways both at baseline and after O_3_ exposure (Table 1). Conversely, CC017 may mount a dampened response to O_3_ exposure that is coordinated by maintaining anti-inflammatory cytokine secretion and AMs that favor cellular proliferation and repair processes upon O_3_ exposure with minimal induction of pro-inflammatory transcriptional activity. Lastly, this analysis suggests that a small set of factors may have important roles in shaping baseline alterations in chromatin organization between CC017 and CC003, thereby poising their responses to O_3_ exposure.

## Discussion

While the adverse health consequences of O_3_ exposure have been appreciated and studied for decades, thorough characterization of the underlying molecular mechanisms and how they develop differentially among individuals is lacking. The use of integrative genomics and high-dimensional phenotyping in genetically diverse animal populations such as the Collaborative Cross (CC) mouse genetic reference panel stands to identify universal biomarkers of effect as well as unique biomarkers of susceptibility.

Here we employed a small panel of CC strains to explore the range of inflammatory, injury, and airway macrophage (AM) genomic responses elicited by acute O_3_ exposure. We used statistical approaches to partition the influences of strain, treatment, and strain-by-treatment interactions on variation in gene expression, which enabled (1) selection of broadly defined O_3_-responsive transcripts; (2) identification of genes whose expression levels are subject to unique genetic and genotype-by-environment interactions; and (3) joint use of these criteria for agnostic selection of CC003 and CC017 as strains of interest for closer scrutiny. We also extended these statistical procedures to examine chromatin accessibility data, an additional, crucial aspect of gene regulation.

Though select studies have examined O_3_ responses in diverse mouse strains, most studies have relied on the use of C57BL/6J to draw broad conclusions. Our results demonstrate that immediate effects of exposure, including inflammatory cell recruitment (namely, neutrophilia), lung injury, and cytokine and chemokine secretion are highly variable across genetic backgrounds, corroborating earlier studies (11, 14, 77). Moreover, while these responses are largely correlated, there is some evidence of regulatory decoupling or separate mechanisms acting on independent time scales, as evidenced by the fact that not all correlations were strong. We also demonstrated that O_3_ exposure causes a characteristic change in AM gene expression shared across multiple genetic backgrounds, and that this signature can be organized into co-expression modules. These modules are enriched for biologically significant pathways bearing relevance to O_3_ responses, including canonical immune signaling pathways and cell cycle progression, and have significant correlations with other aspects of response including neutrophilia and cytokine secretion, suggesting that AM gene expression is causally related to these traits. An important future extension of this work will be to incorporate information collected from additional cell types, as AMs are not the lone driver of phenotypic outcomes. For instance, we have previously shown that acute O_3_ exposure causes noticeable changes in the transcriptome of cells in the conducting airways, in both unique and similar patterns as in AMs (23); thus, profiling their involvement, perhaps with a combination of genomic modalities that enable precise characterization of individual cells or sorted populations will enable description of how diverse cellular signals collectively result in a unified O_3_ response.

This study also revealed widespread effects of strain and strain-by-O_3_ exposure interactions on gene expression. Genes for which we identified a significant effect of strain on expression (termed “G effect” genes) can effectively be considered expression quantitative trait loci (eQTL), albeit with unknown location, though precedent indicates they are more likely local (“*cis*”) than distal (“*trans*”) (17, 78). This statement can be extended to the “GxT” effect genes as well; however, previous studies in *C. elegans* (79), yeast (72), and various cell culture systems (80–83) have shown that context-specific eQTL are often associated with distal signals, in addition to local ones. While we found evidence of genes with these effects when including all strains, the most striking differences were between CC017/Unc and CC003/Unc, a low- and high-responding strain, respectively, as designated by airway neutrophilia and injury. Of note, these strains also exhibit statistically significant differences in BAL neutrophilia and injury after exposure to 1 ppm for 3 hours, indicating that similar genetic architecture may contribute to responses along the concentration-response curve (Supplemental Figure 8). Though these strains have been included in previously published studies (84, 85), minimal experimental evidence exists to indicate specific functional differences between the two that might account for their divergent responses to O_3_ exposure. In a study that screened severity of Zika virus infections across 35 CC strains, CC017/Unc was among the strains with low plasma viral load (two days post-infection), whereas CC003/Unc had twice as many plasma viral copies (84). While neither strain went on to develop clinical features of disease and had similar plasma viral load six days post-infection, baseline differences in immune responsiveness may account for early differences in viral load and differences observed in our study. Additionally, a study examining variation in liver toxicokinetics of trichloroethylene (TCE) across 50 CC strains demonstrated that CC017/Unc had much lower metabolism of TCE to trichloroacetic acid (TCA, the primary driver of TCE toxicity) than CC003/Unc (86). They concluded that differential metabolism is linked to PPARα activity, in a direct relationship. This pathway was not directly implicated in any of the results presented here, but may represent an interesting target for future studies.

While not the focus of the study presented here, an interesting observation is the finding that two strains can arrive at convergent phenotypic profiles traversing a distinct set of transcriptional paths. Specifically, we observed that CC003/Unc and CC039/Unc both mounted strong responses to O_3_ exposure in the form of high airway neutrophilia, lung injury, and elevated cytokine secretion; however, each possessed unique DEGs due to O_3_ (*i.e.*, GxT effect genes). As discussed previously, AM gene expression is only one component of O_3_ responsiveness, thus measuring gene expression in other compartments would verify the extent to which these two strains develop distinct responses at the molecular level. Furthermore, whether these two strains follow similar resolution trajectories after acute O_3_ exposure or are equally sensitive to long-term effects of O_3_ exposure remain open questions. Nevertheless, this observation motivates inclusion of molecular parameters alongside higher-order phenotypes in studies involving genetically diverse individuals (whether humans or model organisms), as the latter alone could obscure the underlying causes.

Across a subset of two CC strains, we see influences of genetic variation on chromatin architecture, specifically at the level of chromatin accessibility. Indeed, these findings have been borne out in integrative studies jointly profiling chromatin accessibility and gene expression across genetically diverse individuals (78, 87, 88). Though specific variants or even regulatory elements are unlikely to be conserved from mouse to human, general principles governing how they influence gene expression and downstream trait variation are highly similar. Moreover, the number of joint genomic and phenotypic analyses and level of experimental precision that can be achieved using the CC or other genetically diverse mouse populations exceeds that which is currently possible in large-scale human genetic studies, leading to more robust discoveries. Therefore, using systems-level analyses that integrate many layers of regulatory information across genetically diverse mouse strains, including ATAC-, ChIP-, and RNA-seq, will be highly valuable for understanding how natural genetic variation modulates or disrupts the regulatory mechanisms underpinning O_3_ responses.

Finally, we did not observe a significant influence of sex on inflammatory, injury, or cytokine secretion in response to O_3_ exposure. Additionally, the most variably expressed genes were only modestly influenced by sex (separation by sex in principal component 9, which accounted for 2.62% total variance). This contrasts with previous studies stating that O_3_ responses vary by sex; however, there is not a consensus stating which sex is more susceptible to O_3_ exposure. Epidemiologic studies have indicated that women have higher rates of hospitalization and mortality following increases in ambient O_3_ levels (89–91), while experimental studies have indicated that female mice display increased pro-inflammatory gene expression profiles and secretion of cytokines and chemokines in response to acute O_3_ exposure (92–94). Otherwise, results regarding sex differences in other indicators of O_3_ response, including those measured in the study presented here (airway inflammation and injury), have been conflicting (95–97). Future work that examines this question more closely, perhaps in a wider set of CC strains, with additional molecular parameters, or at different time points will be able to conclusively determine whether sex might interact with genotype to result in divergent outcomes.

In conclusion, we have shown that respiratory responses to O_3_ exposure are highly variable in a small subset of inbred mouse strains from the CC. A primary future goal will be to identify specific regions of the genome that contribute to inter-strain differences in responsiveness (*i.e.*, map QTL), including a centralized model of response that encompasses DNA variation and chromatin architecture through to phenotypic outcomes.

## Supporting information

HOMER Motif Analyses

## Acknowledgements

The authors would like to acknowledge the assistance of Daniel Vargas and Jessica Bustamante (technical support), Carlton Anderson (UNC Advanced Analytics Core, Luminex processing), the UNC High-Throughput Sequencing Facility (library preparation, RNA- and ATAC-seq), and Courtney Nesline and the UNC Division of Comparative Medicine. This research was funded by NIH Grants ES024965 and ES024965-S1 to S.N.P.K., a UNC Center for Environmental Health and Susceptibility Pilot Project Award to T.S.F. and S.N.P.K. (P30ES010126), T32 training grants to G.J.S. and W.L.C. (ES007126-35), a Leon and Bertha Golberg Postdoctoral Fellowship from the UNC Curriculum in Toxicology and Environmental Medicine to G.J.S., and UNC Dissertation Completion Fellowships to A.T. and B.P.K.

