## Supplemental Figures 1-8 for "Integrative phenotypic and genomic analyses reveal strain-dependent responses to acute ozone exposure and their associations with airway macrophage transcriptional activity"

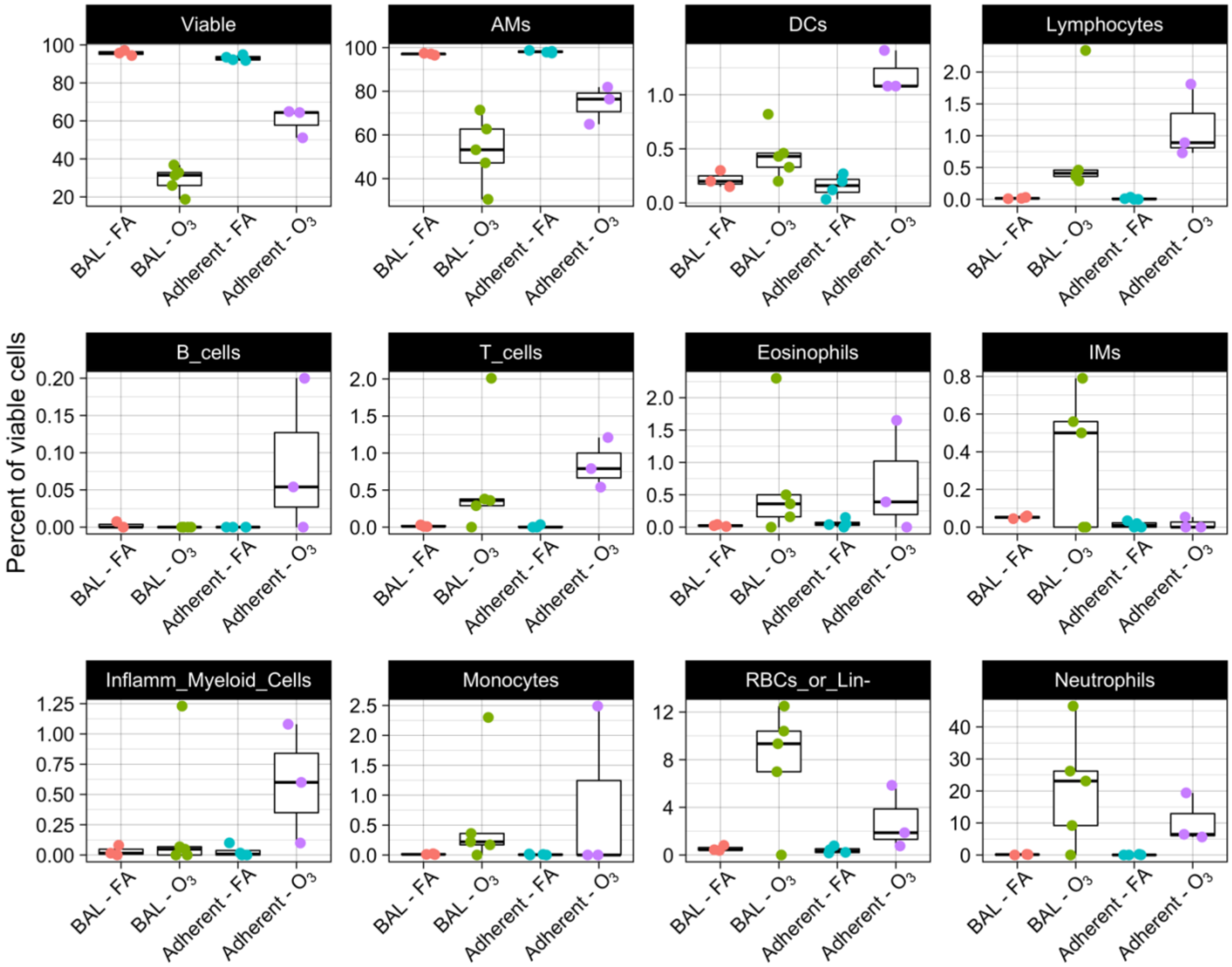


**Supplemental Figure 1. Most cells collected by adherence on plastic culture dishes are alveolar macrophages, as assessed by flow cytometry.** Adult, female C57BL/6J mice were exposed to filtered air (FA) or 2 ppm ozone ($O_3$) for 3 hours and bronchoalveolar lavage (BAL) was collected 21 hours later. Half of the BAL cells were left untouched, while the remainder were plated on untreated culture dishes and allowed to adhere for 2 hours in a 37°C tissue culture incubator. Cells were washed and collected, and both sets (BAL or adherent) were stained and data were acquired using flow cytometry to discriminate between different populations of airway cells. **(**AMs: alveolar macrophages; DCs: dendritic cells; IMs: interstitial macrophages; Inflamm_Myeloid_Cells: inflammatory myeloid cells; RBCs_or_Lin-: red blood cells or lineage-negative cells)

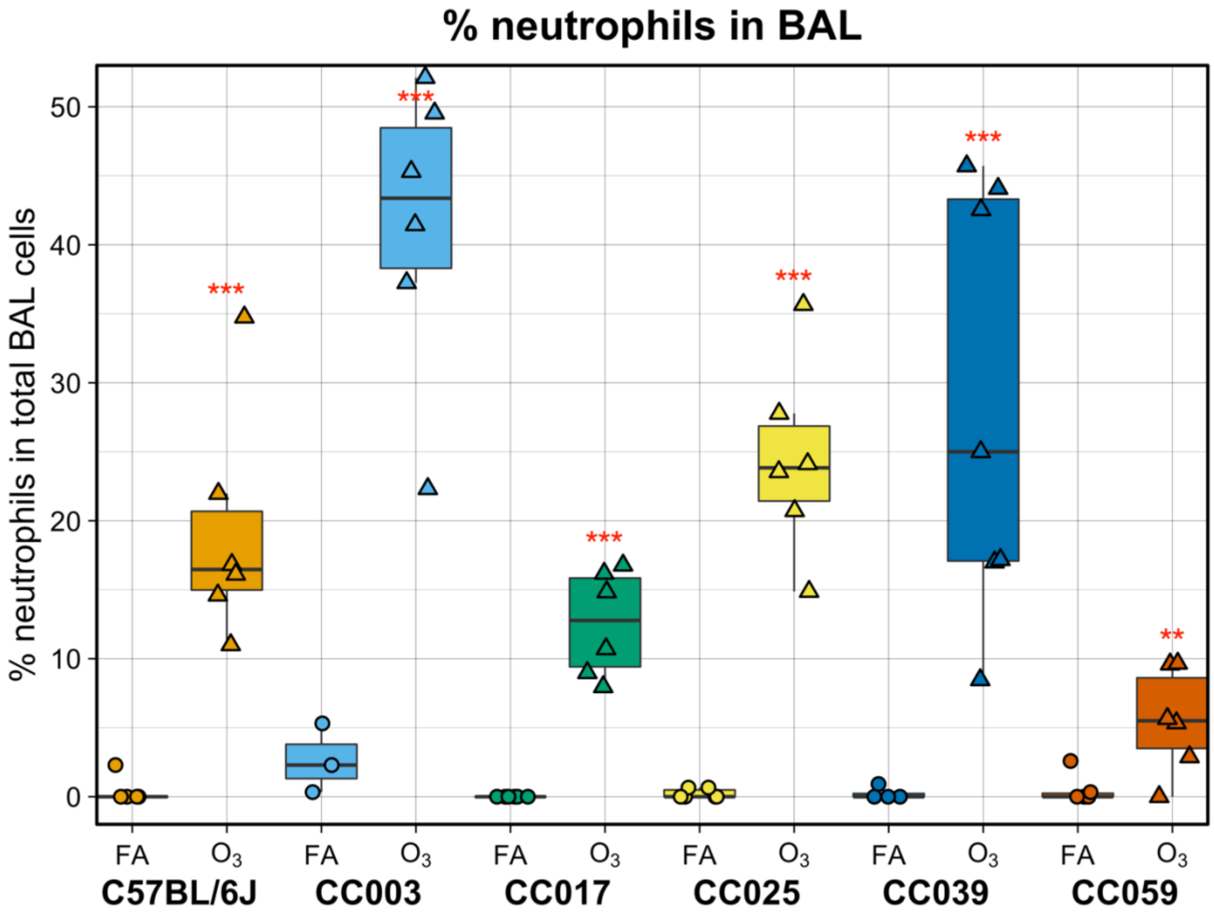


**Supplemental Figure 2. $O_3$ exposure results in increased proportion of neutrophils in the airway cellular infiltrate.** Data are presented as box-and-whiskers plots, which display the distribution from the minimum, first quartile, median, third quartile, and maximum. Individual data points are overlaid, with circles representing FA exposed mice and triangles representing $O_3$ exposed mice. This measurement had a significant strain-by-treatment interaction effect, assessed using a likelihood-ratio test ($p < 0.0005$). (n = 3 per sex/treatment/strain except CC0039 where n = 4 females exposed to $O_3$ and 1 male exposed to FA and CC003 where n = 1 female and 2 males exposed to FA; ** $p < 0.005$, *** $p < 0.0005$, for within-strain contrasts (t-tests) between FA and $O_3$)

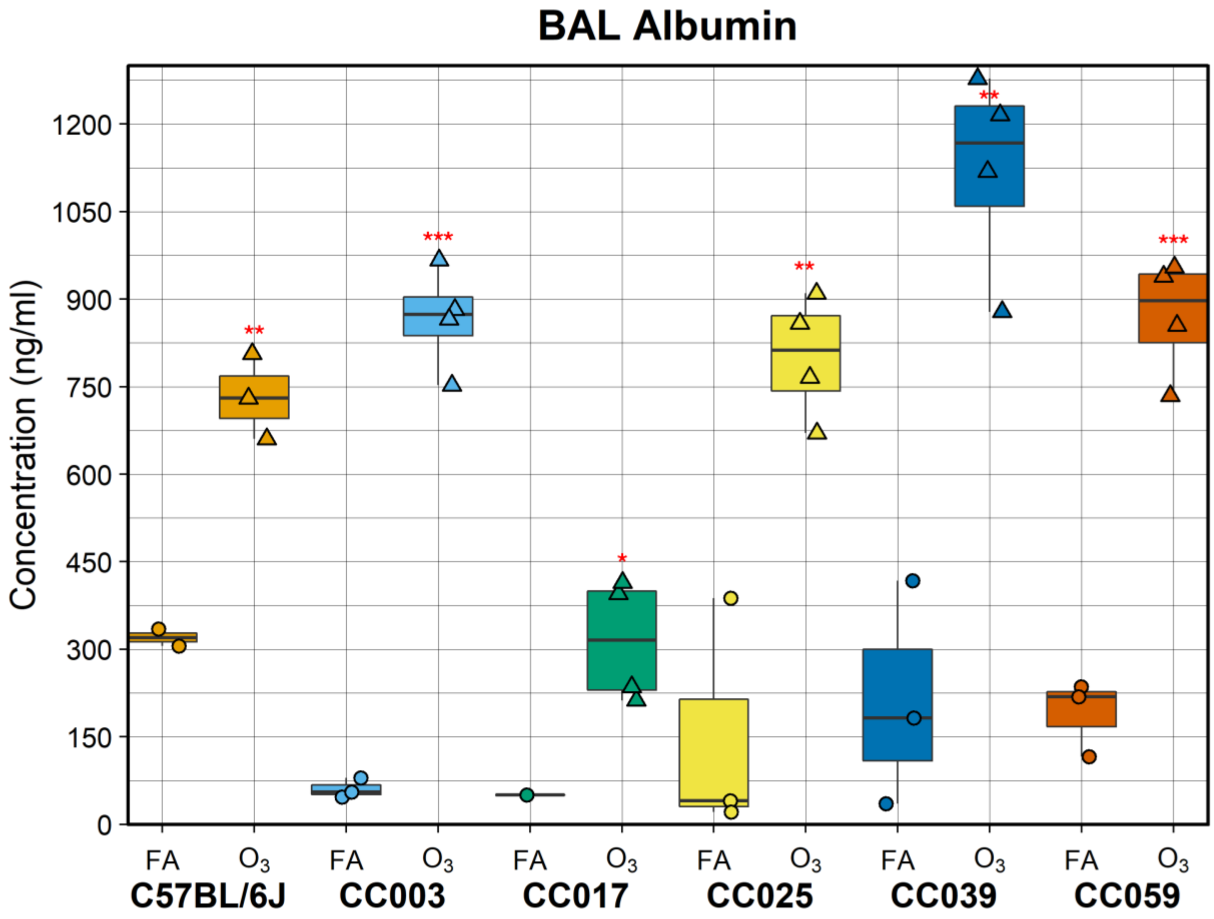


**Supplemental Figure 3. Mice exposed to $O_3$ have increased BAL albumin, confirming results of total protein concentration measurements.** Albumin was measured in a subset of BAL samples by ELISA, as a more specific metric of $O_3$-induced lung permeability. This metric was significantly influenced by a strain-by-treatment interaction effects, assessed using a likelihood-ratio test ($p < 0.05$). (n = 2-3 and 3-4 per sex/strain for FA and $O_3$, respectively; * $p < 0.05$, ** $p < 0.005$, *** $p < 0.0005$, for within-strain contrasts (t-tests) between FA and $O_3$)

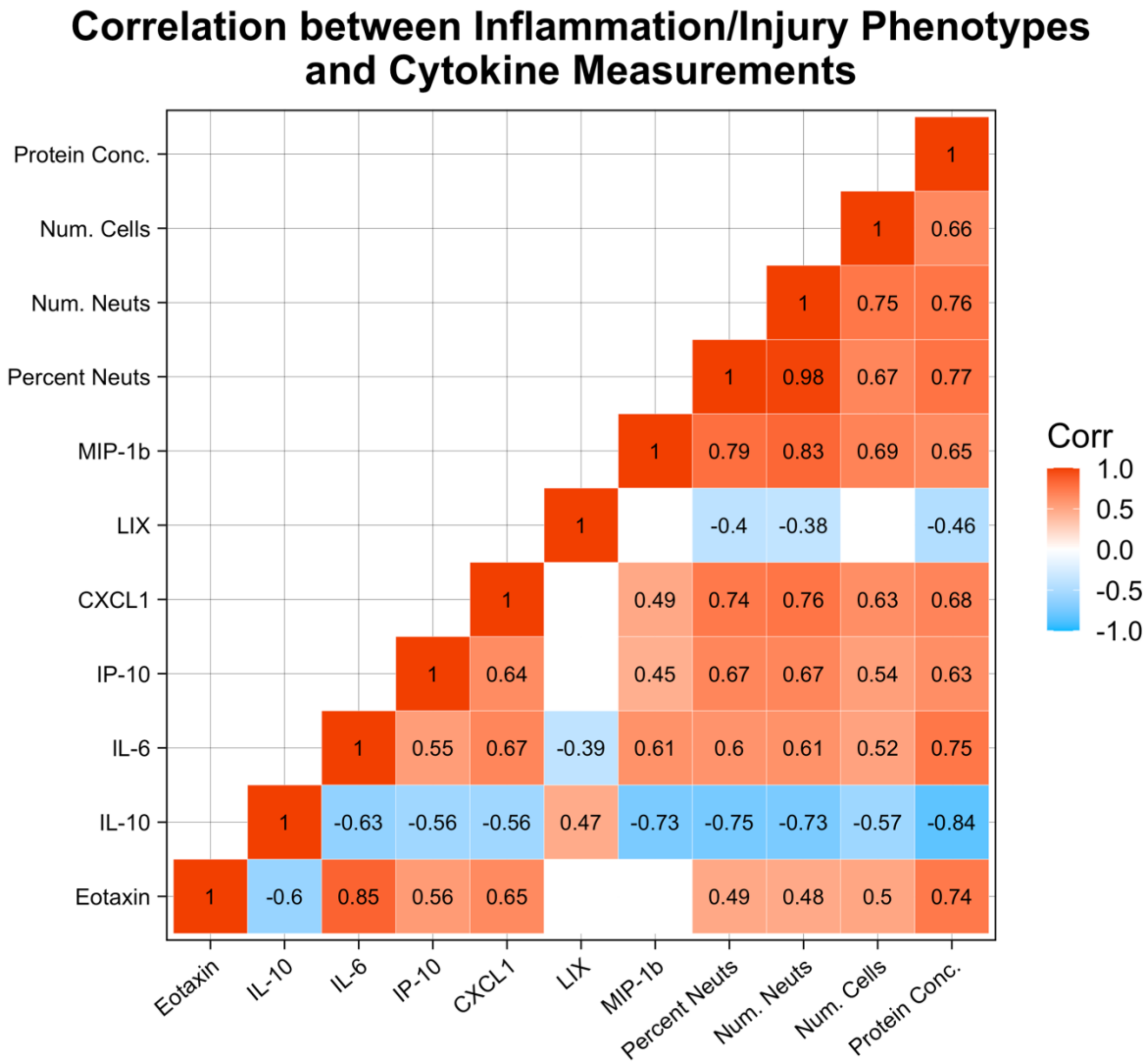


**Supplemental Figure 4. Pairwise correlation between measures of inflammation/injury and BAL cytokines indicate that some aspects of $O_3$ response are regulated in tandem.** Spearman rank-based correlation coefficients were calculated for all samples, for the metrics shown. Correlations were considered statistically significant if $p < 0.05$ (otherwise, a blank square is shown).

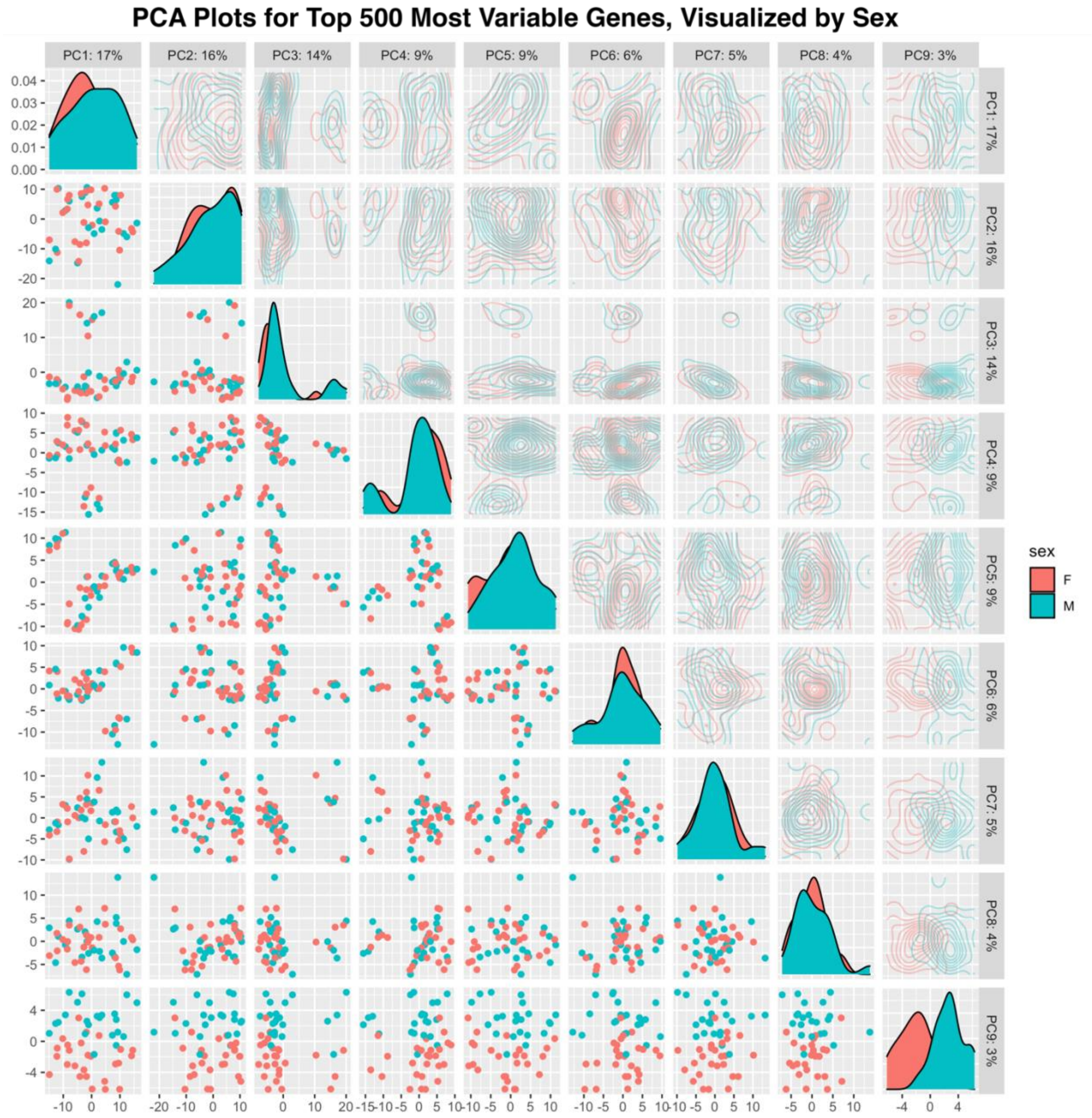


**Supplemental Figure 5. Principal components analysis (PCA) reveals minimal influence of sex on gene expression for the top 500 most variably expressed genes.** PCA was performed, and all pairwise comparisons are shown. Scatterplots in the bottom left-hand corner show individual samples, and indicate that samples don't begin to separate by sex until PC9.

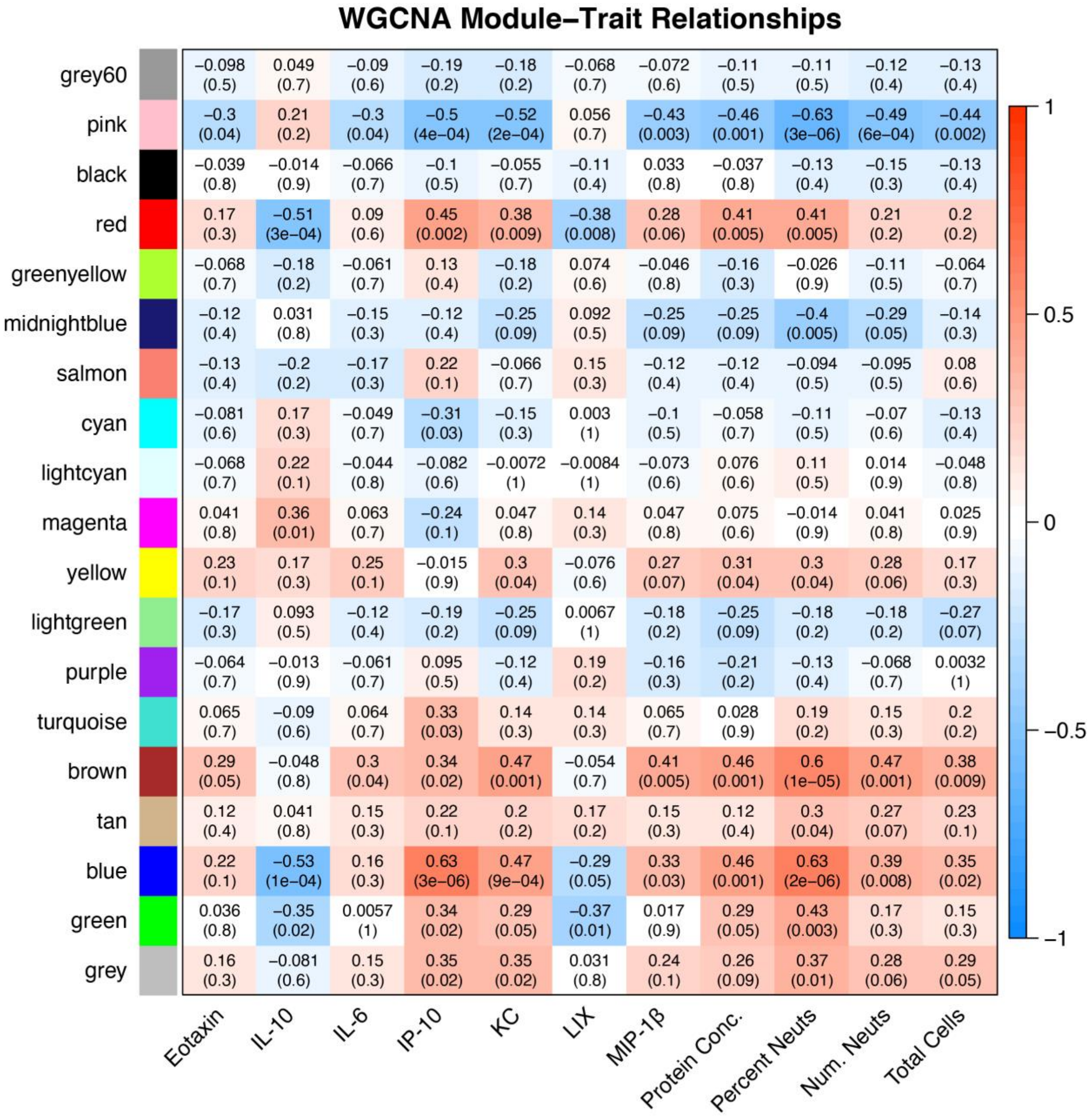


**Supplemental Figure 6. Weighted gene co-expression network analysis (WGCNA) reveals relationships between co-expressed modules of genes and $O_3$-induced inflammatory and injury phenotypes.** WGCNA was performed with all genes that were expressed above a threshold (read counts > samples, 13,594 genes) and grouped into 18 co-expressed modules. “grey60” represents all transcripts that couldn’t be grouped into one of the other 18 modules. Pearson correlation coefficients and corresponding *p*-values were calculated for each pairwise module eigengene and trait relationship (top and bottom number in matrix, respectively)

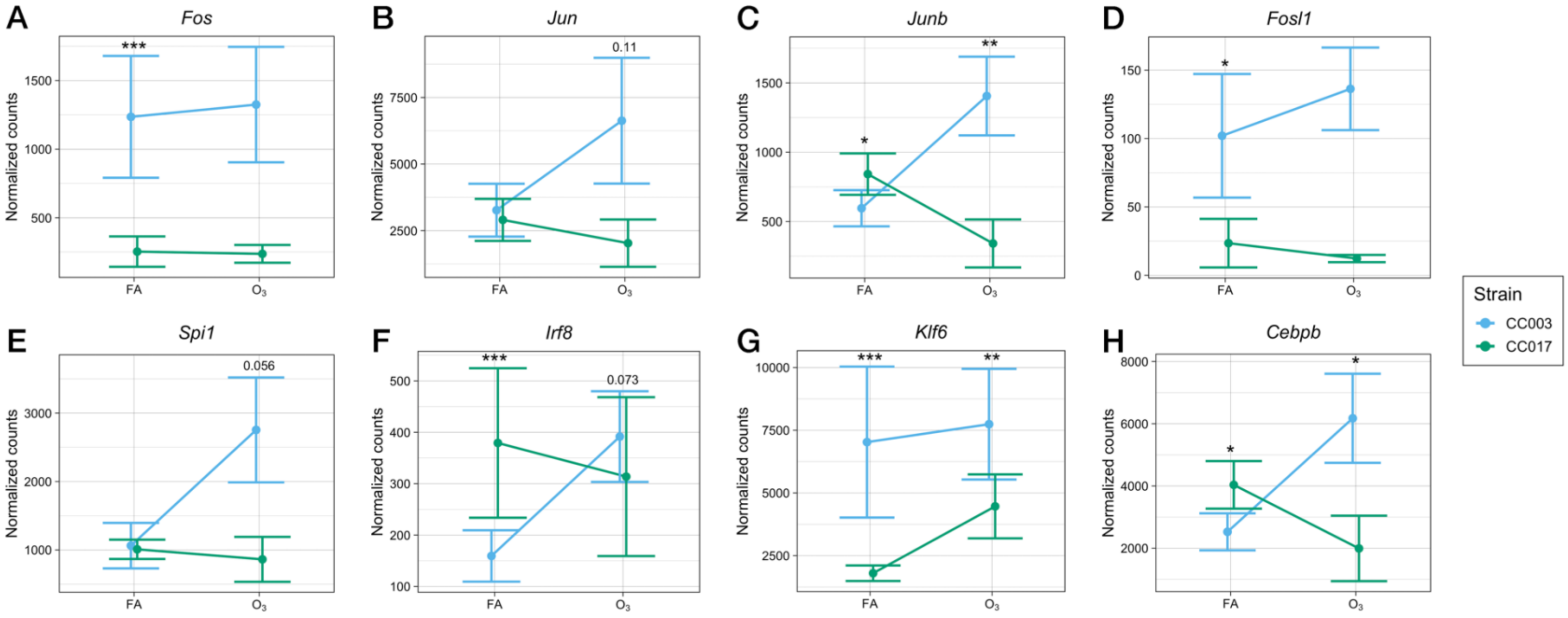


**Supplemental Figure 7. AM gene expression of transcription factors with enriched motifs in sites of differentially accessible chromatin between CC003 and CC017.** We queried gene expression for transcription factors whose binding motifs were enriched within sets of differentially accessible chromatin between CC003 and CC017. In all cases, CC003 had higher expression of the given transcription factor after $O_3$ exposure and, in some cases, at baseline. *Fosl1* is the gene encoding Fra1/FOSL1, and *Spi1* is the gene encoding PU.1. Individual points represent strain means, with standard error. (* $p < 0.05$, ** $p < 0.005$, *** $p < 0.0005$, FDR-adjusted $p$ values for G' effect [asterisk over FA] or GxT effect [asterisk over $O_3$])

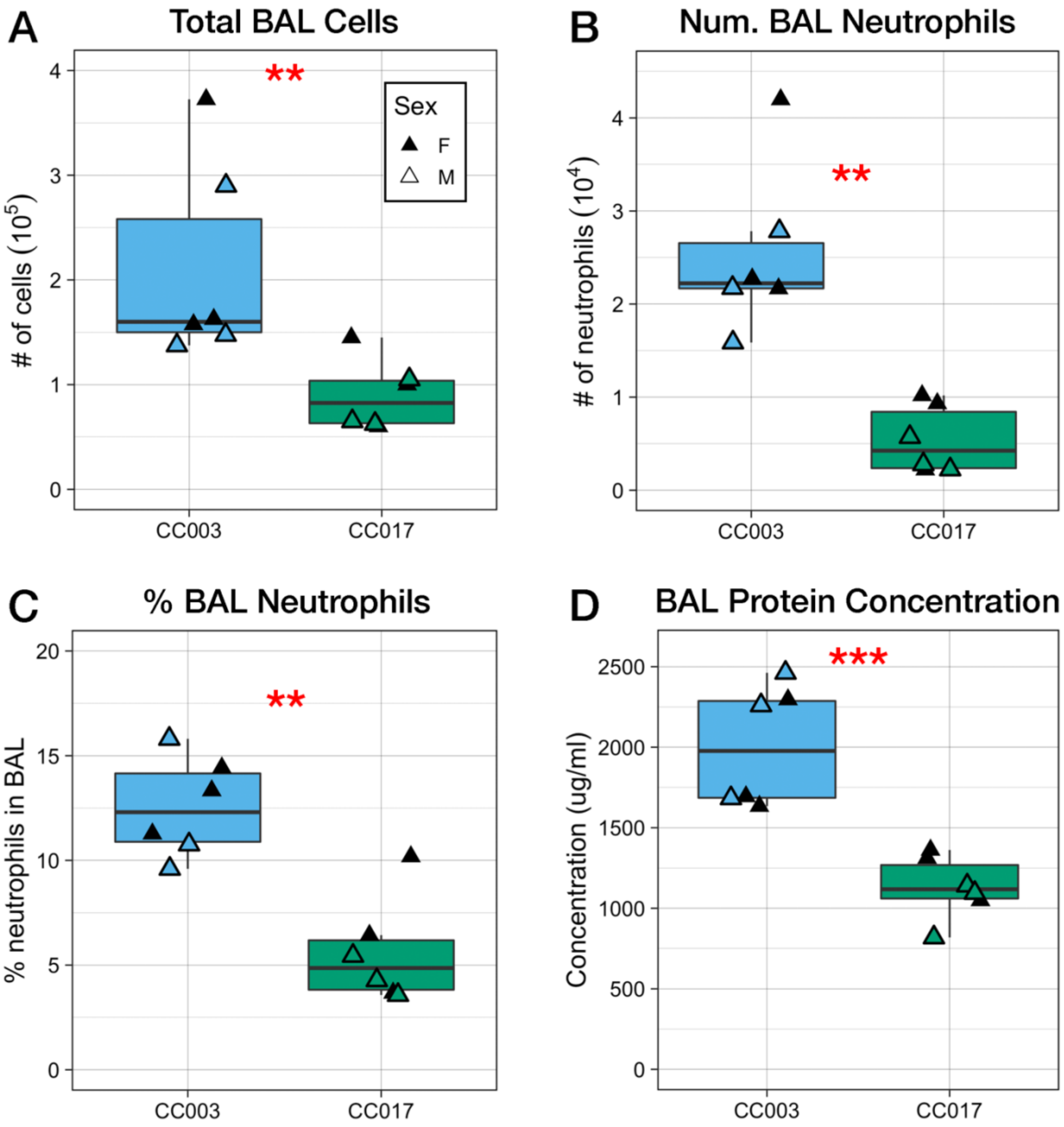


**Supplemental Figure 8. CC003 and CC017 maintain their relative levels of response to a lower concentration of ozone ($O_3$), suggesting that shared mechanisms regulate injury and inflammation along the concentration-response curve.** Female (▲, closed triangle) and male (△, open triangle) CC003 and CC017 mice were exposed to 1 ppm $O_3$ for 3 hours and sacrificed 21 hours later. Bronchoalveolar lavage (BAL) was collected and cellular inflammation (A-C) and injury (D) were measured as described previously in the text. (** $p < 0.005$, *** $p < 0.0005$, for between-strain contrasts performed by t-test)
