## Supplementary material for "Integrative phenotypic and genomic analyses reveal strain-dependent responses to acute ozone exposure and their associations with airway macrophage transcriptional activity": HOMER Motif Analyses: knownResults_G_distal_CC003_up.html

./homer/ATAC\_G\_CC017\_vs\_CC003\_distal\_negative - Homer Known Motif Enrichment Results


### Homer Known Motif Enrichment Results (./homer/ATAC\_G\_CC017\_vs\_CC003\_distal\_negative)

Homer *de novo* Motif Results  
Gene Ontology Enrichment Results  
Known Motif Enrichment Results (txt file)  
Total Target Sequences = 439, Total Background Sequences = 48685

|  |  |  |  |  |  |  |  |  |  |  |  |
| --- | --- | --- | --- | --- | --- | --- | --- | --- | --- | --- | --- |
| Rank | Motif | Name | P-value | log P-pvalue | q-value (Benjamini) | # Target Sequences with Motif | % of Targets Sequences with Motif | # Background Sequences with Motif | % of Background Sequences with Motif | Motif File | SVG |
| 1 | C T A G T C G A G C A T C A T G G C T A T A G C C G A T G T A C C T G A A G C T | JunB(bZIP)/DendriticCells-Junb-ChIP-Seq(GSE36099)/Homer | 1e-48 | -1.107e+02 | 0.0000 | 115.0 | 26.20% | 2458.6 | 5.05% | motif file (matrix) | svg |
| 2 | A C T G C T A G T C G A C G A T C A T G G C T A A T C G C G A T G T A C G C T A A G C T G T A C | Fra1(bZIP)/BT549-Fra1-ChIP-Seq(GSE46166)/Homer | 1e-47 | -1.097e+02 | 0.0000 | 114.0 | 25.97% | 2437.6 | 5.00% | motif file (matrix) | svg |
| 3 | C A T G C T A G T C G A A C G T A C T G C G T A T A G C C G A T T G A C C G T A A G C T G A T C | Fra2(bZIP)/Striatum-Fra2-ChIP-Seq(GSE43429)/Homer | 1e-45 | -1.053e+02 | 0.0000 | 105.0 | 23.92% | 2122.8 | 4.36% | motif file (matrix) | svg |
| 4 | C A G T T G C A A C G T A C T G C G T A A T C G C G A T T G A C C G T A A C G T | BATF(bZIP)/Th17-BATF-ChIP-Seq(GSE39756)/Homer | 1e-45 | -1.038e+02 | 0.0000 | 123.0 | 28.02% | 3052.2 | 6.27% | motif file (matrix) | svg |
| 5 | C T A G T C G A A C G T A C T G C G T A A T G C A C G T G T A C C G T A A G C T G A T C G T A C | Atf3(bZIP)/GBM-ATF3-ChIP-Seq(GSE33912)/Homer | 1e-44 | -1.028e+02 | 0.0000 | 122.0 | 27.79% | 3027.3 | 6.21% | motif file (matrix) | svg |
| 6 | T C G A A C G T C A T G G C T A T A G C C G A T G T A C G C T A A C G T A T G C | AP-1(bZIP)/ThioMac-PU.1-ChIP-Seq(GSE21512)/Homer | 1e-41 | -9.544e+01 | 0.0000 | 126.0 | 28.70% | 3482.9 | 7.15% | motif file (matrix) | svg |
| 7 | C T A G T C G A C G A T A C T G C G T A T A C G A G C T T G A C G C T A A C G T G A T C T A G C | Fosl2(bZIP)/3T3L1-Fosl2-ChIP-Seq(GSE56872)/Homer | 1e-36 | -8.419e+01 | 0.0000 | 81.0 | 18.45% | 1541.3 | 3.16% | motif file (matrix) | svg |
| 8 | C T A G T C G A A C G T A C T G C G T A T A G C C G A T G T A C C G T A A G C T G A T C G T A C | Jun-AP1(bZIP)/K562-cJun-ChIP-Seq(GSE31477)/Homer | 1e-30 | -6.976e+01 | 0.0000 | 63.0 | 14.35% | 1097.4 | 2.25% | motif file (matrix) | svg |
| 9 | G C T A A G T C T A C G T G C A A T C G T C A G G C T A T C G A T C A G A G C T | ELF5(ETS)/T47D-ELF5-ChIP-Seq(GSE30407)/Homer | 1e-21 | -4.968e+01 | 0.0000 | 97.0 | 22.10% | 3609.9 | 7.41% | motif file (matrix) | svg |
| 10 | C T G A A G T C C G A T A G C T A T G C G T A C A C G T A T C G C A G T G C A T | Elf4(ETS)/BMDM-Elf4-ChIP-Seq(GSE88699)/Homer | 1e-21 | -4.911e+01 | 0.0000 | 122.0 | 27.79% | 5388.1 | 11.06% | motif file (matrix) | svg |
| 11 | T C G A C T G A T A G C T G A C T C A G T C A G C G T A C G T A T C A G A G C T | ETV1(ETS)/GIST48-ETV1-ChIP-Seq(GSE22441)/Homer | 1e-18 | -4.253e+01 | 0.0000 | 140.0 | 31.89% | 7270.9 | 14.93% | motif file (matrix) | svg |
| 12 | C G A T T A C G T G A C G A C T C A T G C G T A T A C G A C G T G T A C C T G A | Bach2(bZIP)/OCILy7-Bach2-ChIP-Seq(GSE44420)/Homer | 1e-17 | -4.022e+01 | 0.0000 | 43.0 | 9.79% | 916.7 | 1.88% | motif file (matrix) | svg |
| 13 | A G T C C T G A A G T C C G A T C A G T G A T C A T G C A C T G A T C G G A C T | Fli1(ETS)/CD8-FLI-ChIP-Seq(GSE20898)/Homer | 1e-17 | -3.924e+01 | 0.0000 | 113.0 | 25.74% | 5395.8 | 11.08% | motif file (matrix) | svg |
| 14 | T C G A T A G C T G C A A C T G A C T G C G T A C G T A C T A G G A C T T A C G | ETS1(ETS)/Jurkat-ETS1-ChIP-Seq(GSE17954)/Homer | 1e-16 | -3.851e+01 | 0.0000 | 112.0 | 25.51% | 5373.7 | 11.03% | motif file (matrix) | svg |
| 15 | C G T A T A C G T C G A A C T G A C T G C G T A C G T A T A C G A G C T T A C G | PU.1(ETS)/ThioMac-PU.1-ChIP-Seq(GSE21512)/Homer | 1e-16 | -3.742e+01 | 0.0000 | 74.0 | 16.86% | 2750.3 | 5.65% | motif file (matrix) | svg |
| 16 | T C G A T C G A T A G C G T A C T C A G T A C G C G T A C G T A T C A G A G C T | GABPA(ETS)/Jurkat-GABPa-ChIP-Seq(GSE17954)/Homer | 1e-15 | -3.483e+01 | 0.0000 | 98.0 | 22.32% | 4586.0 | 9.41% | motif file (matrix) | svg |
| 17 | C G T A T G A C T A G C T G C A A C T G A C T G C G T A C G T A T C A G G A C T | ELF3(ETS)/PDAC-ELF3-ChIP-Seq(GSE64557)/Homer | 1e-14 | -3.286e+01 | 0.0000 | 83.0 | 18.91% | 3629.7 | 7.45% | motif file (matrix) | svg |
| 18 | C T G A T A G C T G A C T C A G C T A G G T C A C G T A T C A G A G C T T C A G | ETV4(ETS)/HepG2-ETV4-ChIP-Seq(ENCODE)/Homer | 1e-14 | -3.243e+01 | 0.0000 | 109.0 | 24.83% | 5621.3 | 11.54% | motif file (matrix) | svg |
| 19 | C T G A T G C A T A G C T G A C T A C G T C A G C T G A G C T A T C A G G A C T | ELF1(ETS)/Jurkat-ELF1-ChIP-Seq(SRA014231)/Homer | 1e-14 | -3.241e+01 | 0.0000 | 63.0 | 14.35% | 2307.8 | 4.74% | motif file (matrix) | svg |
| 20 | C G T A T A G C T A G C T G C A A C T G C T A G C G T A C G T A T C A G G A C T | EHF(ETS)/LoVo-EHF-ChIP-Seq(GSE49402)/Homer | 1e-14 | -3.240e+01 | 0.0000 | 120.0 | 27.33% | 6511.6 | 13.37% | motif file (matrix) | svg |
| 21 | C G T A C T G A C G T A C T A G T C G A C T A G A C T G C G T A C G T A T A C G A G C T A T C G | SpiB(ETS)/OCILY3-SPIB-ChIP-Seq(GSE56857)/Homer | 1e-13 | -3.175e+01 | 0.0000 | 42.0 | 9.57% | 1118.4 | 2.30% | motif file (matrix) | svg |
| 22 | A T G C A G T C C T G A A G T C C G A T A C G T A G T C A G T C A C G T A T C G G A C T A C G T | Etv2(ETS)/ES-ER71-ChIP-Seq(GSE59402)/Homer(0.967) | 1e-12 | -2.965e+01 | 0.0000 | 95.0 | 21.64% | 4756.0 | 9.76% | motif file (matrix) | svg |
| 23 | T G C A T C G A T A G C G T A C T C A G C T A G G T C A G C T A T C A G G A C T | ETS(ETS)/Promoter/Homer | 1e-12 | -2.946e+01 | 0.0000 | 48.0 | 10.93% | 1535.8 | 3.15% | motif file (matrix) | svg |
| 24 | T C A G A G C T A T G C C G T A A G C T T C A G C A G T A C T G C T G A A G T C | MITF(bHLH)/MastCells-MITF-ChIP-Seq(GSE48085)/Homer | 1e-10 | -2.452e+01 | 0.0000 | 89.0 | 20.27% | 4727.5 | 9.71% | motif file (matrix) | svg |
| 25 | T A C G T C G A C A G T A C T G G C T A A T G C C G A T G T A C C G T A A C T G T A G C C G T A | NF-E2(bZIP)/K562-NFE2-ChIP-Seq(GSE31477)/Homer | 1e-10 | -2.331e+01 | 0.0000 | 18.0 | 4.10% | 264.7 | 0.54% | motif file (matrix) | svg |
| 26 | T C G A T A G C G T C A A C T G A C T G C G T A C G T A C T A G A G C T T C A G | ERG(ETS)/VCaP-ERG-ChIP-Seq(GSE14097)/Homer | 1e-9 | -2.167e+01 | 0.0000 | 128.0 | 29.16% | 8362.6 | 17.17% | motif file (matrix) | svg |
| 27 | T G C A A G C T A C G T C T A G G A T C C T A G G A T C G T C A C T G A A G T C | CEBP(bZIP)/ThioMac-CEBPb-ChIP-Seq(GSE21512)/Homer | 1e-9 | -2.157e+01 | 0.0000 | 51.0 | 11.62% | 2137.6 | 4.39% | motif file (matrix) | svg |
| 28 | G A T C T C G A A G T C C G A T C G A T A G T C A T G C A C T G A T C G G A C T | Elk1(ETS)/Hela-Elk1-ChIP-Seq(GSE31477)/Homer | 1e-9 | -2.123e+01 | 0.0000 | 55.0 | 12.53% | 2436.7 | 5.00% | motif file (matrix) | svg |
| 29 | T C G A A G C T A C G T A C G T A G T C A G T C A C G T A T C G G A C T A T C G | EWS:ERG-fusion(ETS)/CADO\_ES1-EWS:ERG-ChIP-Seq(SRA014231)/Homer | 1e-9 | -2.074e+01 | 0.0000 | 67.0 | 15.26% | 3350.2 | 6.88% | motif file (matrix) | svg |
| 30 | G T C A G C A T A C T G G T A C G A C T A C T G G C T A A T C G C A G T G T A C C G T A A G C T | Nrf2(bZIP)/Lymphoblast-Nrf2-ChIP-Seq(GSE37589)/Homer | 1e-8 | -2.035e+01 | 0.0000 | 15.0 | 3.42% | 209.4 | 0.43% | motif file (matrix) | svg |
| 31 | T G C A C T G A A T G C G T C A A C T G A C T G C G T A C G T A C T A G A G C T | Ets1-distal(ETS)/CD4+-PolII-ChIP-Seq(Barski\_et\_al.)/Homer | 1e-8 | -1.965e+01 | 0.0000 | 41.0 | 9.34% | 1589.8 | 3.26% | motif file (matrix) | svg |
| 32 | G A T C C T G A A G T C C G A T C G A T G A T C A G T C A C T G A T C G A G C T | Elk4(ETS)/Hela-Elk4-ChIP-Seq(GSE31477)/Homer | 1e-8 | -1.936e+01 | 0.0000 | 51.0 | 11.62% | 2284.5 | 4.69% | motif file (matrix) | svg |
| 33 | C G T A C G T A C G T A G C A T G C A T T A C G G T A C G A C T A C T G C G T A A T C G A C G T G T A C C G T A A G C T | Bach1(bZIP)/K562-Bach1-ChIP-Seq(GSE31477)/Homer | 1e-7 | -1.625e+01 | 0.0000 | 14.0 | 3.19% | 246.1 | 0.51% | motif file (matrix) | svg |
| 34 | T G C A C T G A A G T C G T C A A C T G A C T G C G T A C G T A C T G A A G C T | EWS:FLI1-fusion(ETS)/SK\_N\_MC-EWS:FLI1-ChIP-Seq(SRA014231)/Homer | 1e-6 | -1.386e+01 | 0.0000 | 53.0 | 12.07% | 2889.9 | 5.93% | motif file (matrix) | svg |
| 35 | T C A G T A G C G A C T C A T G C T G A A T C G G C A T G T A C C G T A A C T G T A G C T G C A | MafK(bZIP)/C2C12-MafK-ChIP-Seq(GSE36030)/Homer | 1e-5 | -1.371e+01 | 0.0000 | 32.0 | 7.29% | 1354.5 | 2.78% | motif file (matrix) | svg |
| 36 | T C G A A C G T A C G T C T G A G A T C T C A G G A C T G T C A C G T A A G C T G T C A C T A G A G C T A C G T T C G A | NFIL3(bZIP)/HepG2-NFIL3-ChIP-Seq(Encode)/Homer | 1e-5 | -1.286e+01 | 0.0000 | 40.0 | 9.11% | 1985.4 | 4.08% | motif file (matrix) | svg |
| 37 | T G C A A G C T C T G A A T C G G A C T C T A G G T A C G A T C G T C A A G T C G T A C G A C T C T A G A T C G G C A T C A T G C A T G G A T C G T A C C T G A | CTCF(Zf)/CD4+-CTCF-ChIP-Seq(Barski\_et\_al.)/Homer | 1e-5 | -1.253e+01 | 0.0000 | 19.0 | 4.33% | 602.4 | 1.24% | motif file (matrix) | svg |
| 38 | C T G A A T G C C G T A A C G T A G T C A G T C A C G T A C T G A T C G G C A T | SPDEF(ETS)/VCaP-SPDEF-ChIP-Seq(SRA014231)/Homer | 1e-5 | -1.219e+01 | 0.0001 | 80.0 | 18.22% | 5365.2 | 11.01% | motif file (matrix) | svg |
| 39 | T C A G A G C T A T G C C G T A A G T C T C A G A C G T A T C G T C G A A G T C G A T C T G A C | TFE3(bHLH)/MEF-TFE3-ChIP-Seq(GSE75757)/Homer | 1e-5 | -1.203e+01 | 0.0001 | 14.0 | 3.19% | 354.3 | 0.73% | motif file (matrix) | svg |
| 40 | T A G C C T A G T C G A G A C T A C T G C T G A A G T C T C A G G C A T T G A C C T G A A G C T | Atf7(bZIP)/3T3L1-Atf7-ChIP-Seq(GSE56872)/Homer | 1e-4 | -1.134e+01 | 0.0001 | 40.0 | 9.11% | 2120.1 | 4.35% | motif file (matrix) | svg |
| 41 | A T G C A T G C A T C G T A C G A G C T A G T C G C T A A G T C T C A G G A C T A C T G T C G A | E-box(bHLH)/Promoter/Homer | 1e-4 | -1.115e+01 | 0.0001 | 12.0 | 2.73% | 284.6 | 0.58% | motif file (matrix) | svg |
| 42 | C G T A C G T A C G T A G C A T G C A T A C T G G T A C G A C T C T A G G C T A T A C G G A C T T G A C C G T A A G C T | NFE2L2(bZIP)/HepG2-NFE2L2-ChIP-Seq(Encode)/Homer | 1e-3 | -8.946e+00 | 0.0013 | 11.0 | 2.51% | 305.3 | 0.63% | motif file (matrix) | svg |
| 43 | T A C G T C A G A G C T A T G C C G T A A G T C T C A G A C G T A C T G T C G A | USF1(bHLH)/GM12878-Usf1-ChIP-Seq(GSE32465)/Homer | 1e-3 | -8.826e+00 | 0.0014 | 36.0 | 8.20% | 2058.3 | 4.23% | motif file (matrix) | svg |
| 44 | T A G C A G T C T G A C A G T C C T A G A T C G A G T C C A T G T G A C A G T C G T A C A G T C A G T C G C A T C T A G A T C G G C A T A C T G A T C G G A T C | BORIS(Zf)/K562-CTCFL-ChIP-Seq(GSE32465)/Homer | 1e-3 | -8.468e+00 | 0.0020 | 20.0 | 4.56% | 886.2 | 1.82% | motif file (matrix) | svg |
| 45 | G C T A C T G A T C G A A G T C A G T C C T G A A G T C G T C A C T G A T G C A | RUNX1(Runt)/Jurkat-RUNX1-ChIP-Seq(GSE29180)/Homer | 1e-3 | -8.450e+00 | 0.0020 | 76.0 | 17.31% | 5612.3 | 11.52% | motif file (matrix) | svg |
| 46 | A T G C T C A G T C G A G C A T A C T G C G T A A G T C T C A G G A C T T G A C C G T A A G C T | Atf2(bZIP)/3T3L1-Atf2-ChIP-Seq(GSE56872)/Homer | 1e-3 | -8.385e+00 | 0.0021 | 29.0 | 6.61% | 1553.9 | 3.19% | motif file (matrix) | svg |
| 47 | T C A G G A C T C A G T C T G A A G C T C T A G G A C T T G C A C T G A A G T C | HLF(bZIP)/HSC-HLF.Flag-ChIP-Seq(GSE69817)/Homer | 1e-3 | -8.322e+00 | 0.0021 | 43.0 | 9.79% | 2693.0 | 5.53% | motif file (matrix) | svg |
| 48 | A C T G G A T C G A C T A C T G A C G T C A T G A C T G A C G T A G C T C G A T | RUNX-AML(Runt)/CD4+-PolII-ChIP-Seq(Barski\_et\_al.)/Homer | 1e-3 | -8.165e+00 | 0.0025 | 55.0 | 12.53% | 3753.4 | 7.71% | motif file (matrix) | svg |
| 49 | T C A G A C G T A G T C T C G A A G T C T C A G G C A T C T A G C T A G A G C T | Usf2(bHLH)/C2C12-Usf2-ChIP-Seq(GSE36030)/Homer | 1e-3 | -7.956e+00 | 0.0030 | 27.0 | 6.15% | 1440.3 | 2.96% | motif file (matrix) | svg |
| 50 | T C G A A C G T A C T G C T G A A G T C T C A G A G C T G T A C C G T A A G C T G A T C T C G A | JunD(bZIP)/K562-JunD-ChIP-Seq/Homer | 1e-3 | -7.747e+00 | 0.0036 | 12.0 | 2.73% | 410.5 | 0.84% | motif file (matrix) | svg |
| 51 | T A G C G C T A T C G A C T G A A G T C A G T C C T G A A G T C C G T A C T A G | RUNX(Runt)/HPC7-Runx1-ChIP-Seq(GSE22178)/Homer | 1e-3 | -7.140e+00 | 0.0064 | 55.0 | 12.53% | 3920.0 | 8.05% | motif file (matrix) | svg |
| 52 | T A C G T C G A G A C T A C T G C T G A A G T C T C A G G A C T T G A C C T G A | Atf1(bZIP)/K562-ATF1-ChIP-Seq(GSE31477)/Homer | 1e-2 | -6.518e+00 | 0.0118 | 42.0 | 9.57% | 2861.6 | 5.87% | motif file (matrix) | svg |
| 53 | C T G A C A T G A C T G A C G T A T G C C G T A C A T G T A C G A T G C G C T A T A C G C T G A C T A G A C T G A C G T A T G C C G T A T A G C | RAR:RXR(NR),DR5/ES-RAR-ChIP-Seq(GSE56893)/Homer | 1e-2 | -6.492e+00 | 0.0118 | 6.0 | 1.37% | 134.2 | 0.28% | motif file (matrix) | svg |
| 54 | A T G C G A C T A C T G C A G T G A T C A C G T T A C G T A C G | Smad2(MAD)/ES-SMAD2-ChIP-Seq(GSE29422)/Homer | 1e-2 | -6.330e+00 | 0.0137 | 108.0 | 24.60% | 9196.9 | 18.88% | motif file (matrix) | svg |
| 55 | T A C G A C T G A G C T G T A C C G T A T C G A C T G A A C T G C A T G A C G T A G T C C G T A | COUP-TFII(NR)/K562-NR2F1-ChIP-Seq(Encode)/Homer | 1e-2 | -6.061e+00 | 0.0176 | 98.0 | 22.32% | 8269.3 | 16.98% | motif file (matrix) | svg |
| 56 | A T G C A T C G T A C G A G C T A T C G C T G A A G T C C T A G A G C T A T G C C T G A A T G C | CRE(bZIP)/Promoter/Homer | 1e-2 | -6.045e+00 | 0.0176 | 18.0 | 4.10% | 931.7 | 1.91% | motif file (matrix) | svg |
| 57 | C A G T T C A G G A T C A C T G A C G T C T A G A C T G A C T G G A C T C T A G | Egr1(Zf)/K562-Egr1-ChIP-Seq(GSE32465)/Homer | 1e-2 | -5.456e+00 | 0.0310 | 44.0 | 10.02% | 3219.4 | 6.61% | motif file (matrix) | svg |
| 58 | A G T C A C G T A C T G A G C T A C G T A C G T G T C A A G T C | Foxo1(Forkhead)/RAW-Foxo1-ChIP-Seq(Fan\_et\_al.)/Homer | 1e-2 | -5.381e+00 | 0.0329 | 104.0 | 23.69% | 9066.0 | 18.61% | motif file (matrix) | svg |
| 59 | T G A C G C T A T C G A T G C A A G T C A G T C C G T A A G T C C G T A C T G A G C T A G T A C | RUNX2(Runt)/PCa-RUNX2-ChIP-Seq(GSE33889)/Homer | 1e-2 | -5.312e+00 | 0.0346 | 60.0 | 13.67% | 4744.6 | 9.74% | motif file (matrix) | svg |
| 60 | G A C T C T A G G A T C C A G T A C T G C T G A A T G C G C A T A T G C C T G A | MafA(bZIP)/Islet-MafA-ChIP-Seq(GSE30298)/Homer | 1e-2 | -5.017e+00 | 0.0457 | 60.0 | 13.67% | 4812.7 | 9.88% | motif file (matrix) | svg |
| 61 | C T A G T C G A C G A T C T A G G C A T C A G T C T A G G A T C C G T A G T C A | CEBP:AP1(bZIP)/ThioMac-CEBPb-ChIP-Seq(GSE21512)/Homer | 1e-2 | -4.889e+00 | 0.0511 | 39.0 | 8.88% | 2867.9 | 5.89% | motif file (matrix) | svg |
| 62 | T A C G C T G A C A T G G A T C G T A C G C A T T C A G T A C G A G C T G T C A G A T C G C A T T A C G C G T A C T A G G A T C G A T C C G A T A C T G T C A G | ZNF322(Zf)/HEK293-ZNF322.GFP-ChIP-Seq(GSE58341)/Homer | 1e-2 | -4.840e+00 | 0.0528 | 24.0 | 5.47% | 1547.7 | 3.18% | motif file (matrix) | svg |
| 63 | T G C A G C A T C G A T C G T A C A G T A C T G G T A C C G T A C T G A A G C T G T C A A C T G C T A G G T C A C G A T A C T G G T A C T G C A C G T A A G C T | CEBP:CEBP(bZIP)/MEF-Chop-ChIP-Seq(GSE35681)/Homer | 1e-2 | -4.686e+00 | 0.0606 | 9.0 | 2.05% | 386.2 | 0.79% | motif file (matrix) | svg |
| 64 | C G T A C A T G C A T G A C T G C T A G T C G A G C A T C G A T A G C T A G T C G A T C G T A C | NFkB-p65(RHD)/GM12787-p65-ChIP-Seq(GSE19485)/Homer | 1e-2 | -4.670e+00 | 0.0606 | 30.0 | 6.83% | 2094.5 | 4.30% | motif file (matrix) | svg |
| 65 | T C G A G C A T A C T G C T G A A G T C T C A G G A C T G T A C C G T A A G C T A G T C G A T C | c-Jun-CRE(bZIP)/K562-cJun-ChIP-Seq(GSE31477)/Homer | 1e-2 | -4.642e+00 | 0.0614 | 22.0 | 5.01% | 1404.5 | 2.88% | motif file (matrix) | svg |
