## Supplementary material for "Integrative phenotypic and genomic analyses reveal strain-dependent responses to acute ozone exposure and their associations with airway macrophage transcriptional activity": HOMER Motif Analyses: knownResults_G_distal_CC017_up.html

./homer/ATAC\_G\_CC017\_vs\_CC003\_distal\_positive - Homer Known Motif Enrichment Results


### Homer Known Motif Enrichment Results (./homer/ATAC\_G\_CC017\_vs\_CC003\_distal\_positive)

Homer *de novo* Motif Results  
Gene Ontology Enrichment Results  
Known Motif Enrichment Results (txt file)  
Total Target Sequences = 337, Total Background Sequences = 48986

|  |  |  |  |  |  |  |  |  |  |  |  |
| --- | --- | --- | --- | --- | --- | --- | --- | --- | --- | --- | --- |
| Rank | Motif | Name | P-value | log P-pvalue | q-value (Benjamini) | # Target Sequences with Motif | % of Targets Sequences with Motif | # Background Sequences with Motif | % of Background Sequences with Motif | Motif File | SVG |
| 1 | C G T A C T G A C G T A C T A G T C G A C T A G A C T G C G T A C G T A T A C G A G C T A T C G | SpiB(ETS)/OCILY3-SPIB-ChIP-Seq(GSE56857)/Homer | 1e-12 | -2.849e+01 | 0.0000 | 33.0 | 9.79% | 1023.8 | 2.09% | motif file (matrix) | svg |
| 2 | T C G A A C G T C A T G G C T A T A G C C G A T G T A C G C T A A C G T A T G C | AP-1(bZIP)/ThioMac-PU.1-ChIP-Seq(GSE21512)/Homer | 1e-11 | -2.728e+01 | 0.0000 | 64.0 | 18.99% | 3543.3 | 7.23% | motif file (matrix) | svg |
| 3 | C A T G C T A G T C G A A C G T A C T G C G T A T A G C C G A T T G A C C G T A A G C T G A T C | Fra2(bZIP)/Striatum-Fra2-ChIP-Seq(GSE43429)/Homer | 1e-11 | -2.546e+01 | 0.0000 | 47.0 | 13.95% | 2200.9 | 4.49% | motif file (matrix) | svg |
| 4 | C T A G T C G A G C A T C A T G G C T A T A G C C G A T G T A C C T G A A G C T | JunB(bZIP)/DendriticCells-Junb-ChIP-Seq(GSE36099)/Homer | 1e-10 | -2.499e+01 | 0.0000 | 51.0 | 15.13% | 2567.6 | 5.24% | motif file (matrix) | svg |
| 5 | C T A G T C G A A C G T A C T G C G T A A T G C A C G T G T A C C G T A A G C T G A T C G T A C | Atf3(bZIP)/GBM-ATF3-ChIP-Seq(GSE33912)/Homer | 1e-10 | -2.405e+01 | 0.0000 | 56.0 | 16.62% | 3080.9 | 6.29% | motif file (matrix) | svg |
| 6 | A C T G C T A G T C G A C G A T C A T G G C T A A T C G C G A T G T A C G C T A A G C T G T A C | Fra1(bZIP)/BT549-Fra1-ChIP-Seq(GSE46166)/Homer | 1e-10 | -2.303e+01 | 0.0000 | 49.0 | 14.54% | 2539.1 | 5.18% | motif file (matrix) | svg |
| 7 | C A G T T G C A A C G T A C T G C G T A A T C G C G A T T G A C C G T A A C G T | BATF(bZIP)/Th17-BATF-ChIP-Seq(GSE39756)/Homer | 1e-9 | -2.109e+01 | 0.0000 | 54.0 | 16.02% | 3150.8 | 6.43% | motif file (matrix) | svg |
| 8 | C T A G T C G A C G A T A C T G C G T A T A C G A G C T T G A C G C T A A C G T G A T C T A G C | Fosl2(bZIP)/3T3L1-Fosl2-ChIP-Seq(GSE56872)/Homer | 1e-8 | -2.023e+01 | 0.0000 | 35.0 | 10.39% | 1568.9 | 3.20% | motif file (matrix) | svg |
| 9 | C T A G T C G A A C G T A C T G C G T A T A G C C G A T G T A C C G T A A G C T G A T C G T A C | Jun-AP1(bZIP)/K562-cJun-ChIP-Seq(GSE31477)/Homer | 1e-8 | -1.890e+01 | 0.0000 | 28.0 | 8.31% | 1111.7 | 2.27% | motif file (matrix) | svg |
| 10 | C G T A T A C G T C G A A C T G A C T G C G T A C G T A T A C G A G C T T A C G | PU.1(ETS)/ThioMac-PU.1-ChIP-Seq(GSE21512)/Homer | 1e-5 | -1.340e+01 | 0.0001 | 41.0 | 12.17% | 2661.6 | 5.43% | motif file (matrix) | svg |
| 11 | G C T A A G T C T A C G T G C A A T C G T C A G G C T A T C G A T C A G A G C T | ELF5(ETS)/T47D-ELF5-ChIP-Seq(GSE30407)/Homer | 1e-5 | -1.244e+01 | 0.0001 | 50.0 | 14.84% | 3695.3 | 7.54% | motif file (matrix) | svg |
| 12 | C T G A A G T C C G A T A G C T A T G C G T A C A C G T A T C G C A G T G C A T | Elf4(ETS)/BMDM-Elf4-ChIP-Seq(GSE88699)/Homer | 1e-4 | -1.147e+01 | 0.0004 | 62.0 | 18.40% | 5156.1 | 10.53% | motif file (matrix) | svg |
| 13 | T C A G A G C T A T G C C G T A A G C T T C A G C A G T A C T G C T G A A G T C | MITF(bHLH)/MastCells-MITF-ChIP-Seq(GSE48085)/Homer | 1e-4 | -9.679e+00 | 0.0020 | 54.0 | 16.02% | 4556.2 | 9.30% | motif file (matrix) | svg |
| 14 | C G T A T A G C T A G C T G C A A C T G C T A G C G T A C G T A T C A G G A C T | EHF(ETS)/LoVo-EHF-ChIP-Seq(GSE49402)/Homer | 1e-4 | -9.548e+00 | 0.0021 | 71.0 | 21.07% | 6571.0 | 13.41% | motif file (matrix) | svg |
| 15 | C G T A T G A C T A G C T G C A A C T G A C T G C G T A C G T A T C A G G A C T | ELF3(ETS)/PDAC-ELF3-ChIP-Seq(GSE64557)/Homer | 1e-4 | -9.502e+00 | 0.0021 | 46.0 | 13.65% | 3691.1 | 7.54% | motif file (matrix) | svg |
| 16 | T C G A T A G C G T C A A C T G A C T G C G T A C G T A C T A G A G C T T C A G | ERG(ETS)/VCaP-ERG-ChIP-Seq(GSE14097)/Homer | 1e-3 | -9.005e+00 | 0.0032 | 82.0 | 24.33% | 8037.2 | 16.41% | motif file (matrix) | svg |
| 17 | T C G A C T G A T A G C T G A C T C A G T C A G C G T A C G T A T C A G A G C T | ETV1(ETS)/GIST48-ETV1-ChIP-Seq(GSE22441)/Homer | 1e-3 | -8.453e+00 | 0.0052 | 71.0 | 21.07% | 6811.3 | 13.91% | motif file (matrix) | svg |
| 18 | T C G A T C G A T A G C G T A C T C A G T A C G C G T A C G T A T C A G A G C T | GABPA(ETS)/Jurkat-GABPa-ChIP-Seq(GSE17954)/Homer | 1e-3 | -8.162e+00 | 0.0066 | 49.0 | 14.54% | 4259.5 | 8.70% | motif file (matrix) | svg |
| 19 | T C G A T A G C T G C A A C T G A C T G C G T A C G T A C T A G G A C T T A C G | ETS1(ETS)/Jurkat-ETS1-ChIP-Seq(GSE17954)/Homer | 1e-3 | -8.144e+00 | 0.0066 | 57.0 | 16.91% | 5195.9 | 10.61% | motif file (matrix) | svg |
| 20 | A G T C C T A G C T A G A G T C G A T C G T A C A G T C C T A G A G T C A G T C A G T C G T A C | Sp2(Zf)/HEK293-Sp2.eGFP-ChIP-Seq(Encode)/Homer | 1e-3 | -8.008e+00 | 0.0069 | 59.0 | 17.51% | 5461.7 | 11.15% | motif file (matrix) | svg |
| 21 | T G C A T C G A T A G C G T A C T C A G C T A G G T C A G C T A T C A G G A C T | ETS(ETS)/Promoter/Homer | 1e-3 | -7.257e+00 | 0.0139 | 20.0 | 5.93% | 1284.1 | 2.62% | motif file (matrix) | svg |
| 22 | C G T A C T A G A C T G A C T G G A C T C T A G C A G T C T A G C A T G G A T C | KLF5(Zf)/LoVo-KLF5-ChIP-Seq(GSE49402)/Homer | 1e-2 | -6.823e+00 | 0.0205 | 53.0 | 15.73% | 5004.7 | 10.22% | motif file (matrix) | svg |
| 23 | C T G A T G C A T A G C T G A C T A C G T C A G C T G A G C T A T C A G G A C T | ELF1(ETS)/Jurkat-ELF1-ChIP-Seq(SRA014231)/Homer | 1e-2 | -6.814e+00 | 0.0205 | 27.0 | 8.01% | 2047.2 | 4.18% | motif file (matrix) | svg |
| 24 | T C A G G A C T C A G T C T G A A G C T C T A G G A C T T G C A C T G A A G T C | HLF(bZIP)/HSC-HLF.Flag-ChIP-Seq(GSE69817)/Homer | 1e-2 | -6.681e+00 | 0.0216 | 39.0 | 11.57% | 3392.1 | 6.93% | motif file (matrix) | svg |
| 25 | T G C A A G C T A C G T C T A G G A T C C T A G G A T C G T C A C T G A A G T C | CEBP(bZIP)/ThioMac-CEBPb-ChIP-Seq(GSE21512)/Homer | 1e-2 | -6.570e+00 | 0.0232 | 32.0 | 9.50% | 2625.5 | 5.36% | motif file (matrix) | svg |
| 26 | T A G C C T A G T C G A G A C T A C T G C T G A A G T C T C A G G C A T T G A C C T G A A G C T | Atf7(bZIP)/3T3L1-Atf7-ChIP-Seq(GSE56872)/Homer | 1e-2 | -6.539e+00 | 0.0232 | 27.0 | 8.01% | 2088.0 | 4.26% | motif file (matrix) | svg |
| 27 | A T G C T C A G T C G A G C A T A C T G C G T A A G T C T C A G G A C T T G A C C G T A A G C T | Atf2(bZIP)/3T3L1-Atf2-ChIP-Seq(GSE56872)/Homer | 1e-2 | -5.875e+00 | 0.0431 | 20.0 | 5.93% | 1446.6 | 2.95% | motif file (matrix) | svg |
| 28 | A T G C A T G C A T C G T A C G A G C T A G T C G C T A A G T C T C A G G A C T A C T G T C G A | E-box(bHLH)/Promoter/Homer | 1e-2 | -5.439e+00 | 0.0642 | 6.0 | 1.78% | 218.8 | 0.45% | motif file (matrix) | svg |
| 29 | C T G A T C A G C A G T C T A G A C T G C T A G G A T C A T C G A C T G C T G A T C A G G A T C | Sp5(Zf)/mES-Sp5.Flag-ChIP-Seq(GSE72989)/Homer | 1e-2 | -5.377e+00 | 0.0660 | 37.0 | 10.98% | 3423.6 | 6.99% | motif file (matrix) | svg |
| 30 | C G A T T A C G T G A C G A C T C A T G C G T A T A C G A C G T G T A C C T G A | Bach2(bZIP)/OCILy7-Bach2-ChIP-Seq(GSE44420)/Homer | 1e-2 | -5.316e+00 | 0.0678 | 14.0 | 4.15% | 913.1 | 1.86% | motif file (matrix) | svg |
| 31 | T C A G A G C T A T G C C G T A A G T C T C A G A C G T A T C G T C G A A G T C G A T C T G A C | TFE3(bHLH)/MEF-TFE3-ChIP-Seq(GSE75757)/Homer | 1e-2 | -5.191e+00 | 0.0743 | 7.0 | 2.08% | 304.1 | 0.62% | motif file (matrix) | svg |
| 32 | T A C G T C A G A G C T A T G C C G T A A G T C T C A G A C G T A C T G T C G A | USF1(bHLH)/GM12878-Usf1-ChIP-Seq(GSE32465)/Homer | 1e-2 | -5.028e+00 | 0.0848 | 23.0 | 6.82% | 1890.9 | 3.86% | motif file (matrix) | svg |
| 33 | C T A G T C G A C G A T C T A G G C A T C A G T C T A G G A T C C G T A G T C A | CEBP:AP1(bZIP)/ThioMac-CEBPb-ChIP-Seq(GSE21512)/Homer | 1e-2 | -4.632e+00 | 0.1221 | 33.0 | 9.79% | 3115.5 | 6.36% | motif file (matrix) | svg |
| 34 | C T A G T C A G C A G T T C A G A C T G A C T G G A T C C T A G A C T G C T A G T C A G A T G C | KLF14(Zf)/HEK293-KLF14.GFP-ChIP-Seq(GSE58341)/Homer | 1e-2 | -4.621e+00 | 0.1221 | 63.0 | 18.69% | 6853.8 | 13.99% | motif file (matrix) | svg |
| 35 | T C A G A T C G G A C T A C T G G A C T C A G T C T A G C G T A G T A C C G T A C T A G A T C G | Tbx20(T-box)/Heart-Tbx20-ChIP-Seq(GSE29636)/Homer | 1e-2 | -4.616e+00 | 0.1221 | 15.0 | 4.45% | 1096.6 | 2.24% | motif file (matrix) | svg |
