## Supplementary material for "Integrative phenotypic and genomic analyses reveal strain-dependent responses to acute ozone exposure and their associations with airway macrophage transcriptional activity": HOMER Motif Analyses: knownResults_G_proximal_CC003_up.html

./homer/ATAC\_G\_CC017\_vs\_CC003\_promoter\_negative - Homer Known Motif Enrichment Results


### Homer Known Motif Enrichment Results (./homer/ATAC\_G\_CC017\_vs\_CC003\_promoter\_negative)

Homer *de novo* Motif Results  
Gene Ontology Enrichment Results  
Known Motif Enrichment Results (txt file)  
Total Target Sequences = 92, Total Background Sequences = 48965

|  |  |  |  |  |  |  |  |  |  |  |  |
| --- | --- | --- | --- | --- | --- | --- | --- | --- | --- | --- | --- |
| Rank | Motif | Name | P-value | log P-pvalue | q-value (Benjamini) | # Target Sequences with Motif | % of Targets Sequences with Motif | # Background Sequences with Motif | % of Background Sequences with Motif | Motif File | SVG |
| 1 | A G T C C T G A A G T C C G A T C A G T G A T C A T G C A C T G A T C G G A C T | Fli1(ETS)/CD8-FLI-ChIP-Seq(GSE20898)/Homer | 1e-5 | -1.342e+01 | 0.0006 | 30.0 | 32.61% | 6492.8 | 13.26% | motif file (matrix) | svg |
| 2 | T C G A C T G A T A G C T G A C T C A G T C A G C G T A C G T A T C A G A G C T | ETV1(ETS)/GIST48-ETV1-ChIP-Seq(GSE22441)/Homer | 1e-4 | -9.659e+00 | 0.0132 | 30.0 | 32.61% | 7835.3 | 16.00% | motif file (matrix) | svg |
| 3 | T C G A T A G C T G C A A C T G A C T G C G T A C G T A C T A G G A C T T A C G | ETS1(ETS)/Jurkat-ETS1-ChIP-Seq(GSE17954)/Homer | 1e-4 | -9.217e+00 | 0.0137 | 24.0 | 26.09% | 5678.1 | 11.60% | motif file (matrix) | svg |
| 4 | C T G A A G T C C G A T A G C T A T G C G T A C A C G T A T C G C A G T G C A T | Elf4(ETS)/BMDM-Elf4-ChIP-Seq(GSE88699)/Homer | 1e-3 | -9.110e+00 | 0.0137 | 24.0 | 26.09% | 5718.0 | 11.68% | motif file (matrix) | svg |
| 5 | A T G C A G T C C T G A A G T C C G A T A C G T A G T C A G T C A C G T A T C G G A C T A C G T | Etv2(ETS)/ES-ER71-ChIP-Seq(GSE59402)/Homer(0.967) | 1e-3 | -9.037e+00 | 0.0137 | 21.0 | 22.83% | 4644.1 | 9.49% | motif file (matrix) | svg |
| 6 | C T G A T A G C T G A C T C A G C T A G G T C A C G T A T C A G A G C T T C A G | ETV4(ETS)/HepG2-ETV4-ChIP-Seq(ENCODE)/Homer | 1e-3 | -8.738e+00 | 0.0137 | 27.0 | 29.35% | 7021.7 | 14.34% | motif file (matrix) | svg |
| 7 | C T G A T G C A T A G C T G A C T A C G T C A G C T G A G C T A T C A G G A C T | ELF1(ETS)/Jurkat-ELF1-ChIP-Seq(SRA014231)/Homer | 1e-3 | -8.189e+00 | 0.0164 | 17.0 | 18.48% | 3518.1 | 7.19% | motif file (matrix) | svg |
| 8 | T C G A T A G C G T C A A C T G A C T G C G T A C G T A C T A G A G C T T C A G | ERG(ETS)/VCaP-ERG-ChIP-Seq(GSE14097)/Homer | 1e-3 | -8.018e+00 | 0.0171 | 29.0 | 31.52% | 8154.7 | 16.66% | motif file (matrix) | svg |
| 9 | C G T A C T G A C G T A C T A G T C G A C T A G A C T G C G T A C G T A T A C G A G C T A T C G | SpiB(ETS)/OCILY3-SPIB-ChIP-Seq(GSE56857)/Homer | 1e-2 | -6.629e+00 | 0.0608 | 7.0 | 7.61% | 871.5 | 1.78% | motif file (matrix) | svg |
| 10 | T G A C C T A G T C A G G T C A C G T A T C A G C G A T T C A G T C G A T G C A C T G A T A G C | PU.1-IRF(ETS:IRF)/Bcell-PU.1-ChIP-Seq(GSE21512)/Homer | 1e-2 | -6.382e+00 | 0.0701 | 21.0 | 22.83% | 5671.8 | 11.58% | motif file (matrix) | svg |
| 11 | T C G A T C G A T A G C G T A C T C A G T A C G C G T A C G T A T C A G A G C T | GABPA(ETS)/Jurkat-GABPa-ChIP-Seq(GSE17954)/Homer | 1e-2 | -5.651e+00 | 0.1322 | 19.0 | 20.65% | 5216.7 | 10.65% | motif file (matrix) | svg |
| 12 | G A T C C T G A A G T C C G A T C G A T G A T C A G T C A C T G A T C G A G C T | Elk4(ETS)/Hela-Elk4-ChIP-Seq(GSE31477)/Homer | 1e-2 | -5.470e+00 | 0.1453 | 15.0 | 16.30% | 3750.3 | 7.66% | motif file (matrix) | svg |
| 13 | G C T A A G C T T A C G G T C A C T G A C G A T C T G A G C A T C A G T A G T C | Brn2(POU,Homeobox)/NPC-Brn2-ChIP-Seq(GSE35496)/Homer | 1e-2 | -5.367e+00 | 0.1487 | 3.0 | 3.26% | 177.2 | 0.36% | motif file (matrix) | svg |
| 14 | C G T A C T A G A C T G A C T G G A C T C T A G C A G T C T A G C A T G G A T C | KLF5(Zf)/LoVo-KLF5-ChIP-Seq(GSE49402)/Homer | 1e-2 | -4.924e+00 | 0.2150 | 29.0 | 31.52% | 9923.7 | 20.27% | motif file (matrix) | svg |
| 15 | A G T C G A C T C A G T A C T G C T A G T G A C G C T A A T G C G C A T A T C G C G A T A C T G G A T C G T A C G T C A C T G A | NF1(CTF)/LNCAP-NF1-ChIP-Seq(Unpublished)/Homer | 1e-2 | -4.794e+00 | 0.2284 | 9.0 | 9.78% | 1856.4 | 3.79% | motif file (matrix) | svg |
| 16 | T C G A C T A G A G T C A G T C C G T A C G T A A C G T T A G C T C A G T A C G | NFY(CCAAT)/Promoter/Homer | 1e-2 | -4.698e+00 | 0.2359 | 14.0 | 15.22% | 3698.9 | 7.55% | motif file (matrix) | svg |
| 17 | T G C A T C G A T A G C G T A C T C A G C T A G G T C A G C T A T C A G G A C T | ETS(ETS)/Promoter/Homer | 1e-2 | -4.672e+00 | 0.2359 | 10.0 | 10.87% | 2236.9 | 4.57% | motif file (matrix) | svg |
