## Supplementary material for "Integrative phenotypic and genomic analyses reveal strain-dependent responses to acute ozone exposure and their associations with airway macrophage transcriptional activity": HOMER Motif Analyses: knownResults_G_proximal_CC017_up.html

./homer/ATAC\_G\_CC017\_vs\_CC003\_promoter\_positive - Homer Known Motif Enrichment Results


### Homer Known Motif Enrichment Results (./homer/ATAC\_G\_CC017\_vs\_CC003\_promoter\_positive)

Homer *de novo* Motif Results  
Gene Ontology Enrichment Results  
Known Motif Enrichment Results (txt file)  
Total Target Sequences = 105, Total Background Sequences = 49535

|  |  |  |  |  |  |  |  |  |  |  |  |
| --- | --- | --- | --- | --- | --- | --- | --- | --- | --- | --- | --- |
| Rank | Motif | Name | P-value | log P-pvalue | q-value (Benjamini) | # Target Sequences with Motif | % of Targets Sequences with Motif | # Background Sequences with Motif | % of Background Sequences with Motif | Motif File | SVG |
| 1 | A G T C C T A G C T A G A G T C G A T C G T A C A G T C C T A G A G T C A G T C A G T C G T A C | Sp2(Zf)/HEK293-Sp2.eGFP-ChIP-Seq(Encode)/Homer | 1e-6 | -1.558e+01 | 0.0001 | 49.0 | 46.67% | 11630.9 | 23.48% | motif file (matrix) | svg |
| 2 | T A C G C T A G A T G C G A T C G T A C A G T C C T A G A G T C A G T C A G T C G T A C A G T C | Sp1(Zf)/Promoter/Homer | 1e-4 | -9.938e+00 | 0.0100 | 19.0 | 18.10% | 3238.4 | 6.54% | motif file (matrix) | svg |
| 3 | C T A G G T A C A G T C T G C A A G T C C T G A A G T C A G T C A G T C G C T A | Klf4(Zf)/mES-Klf4-ChIP-Seq(GSE11431)/Homer | 1e-3 | -7.537e+00 | 0.0735 | 15.0 | 14.29% | 2676.1 | 5.40% | motif file (matrix) | svg |
| 4 | T C G A C T A G A G T C A G T C C G T A C G T A A C G T T A G C T C A G T A C G | NFY(CCAAT)/Promoter/Homer | 1e-3 | -7.382e+00 | 0.0735 | 19.0 | 18.10% | 3954.8 | 7.99% | motif file (matrix) | svg |
| 5 | G A C T T C A G C T A G A G T C A G T C G T A C A G T C C T G A A G T C A G T C A G T C G A C T A G T C A C T G A T G C | KLF3(Zf)/MEF-Klf3-ChIP-Seq(GSE44748)/Homer | 1e-2 | -6.526e+00 | 0.1213 | 19.0 | 18.10% | 4250.7 | 8.58% | motif file (matrix) | svg |
| 6 | G C T A A C G T C T A G G T C A C G T A A C G T C G T A C G A T C A G T A G T C C G T A A C G T C T A G C T G A A T C G | OCT:OCT(POU,Homeobox)/NPC-Brn1-ChIP-Seq(GSE35496)/Homer | 1e-2 | -6.461e+00 | 0.1213 | 2.0 | 1.90% | 27.7 | 0.06% | motif file (matrix) | svg |
| 7 | C G T A C T A G A C T G A C T G G A C T C T A G C A G T C T A G C A T G G A T C | KLF5(Zf)/LoVo-KLF5-ChIP-Seq(GSE49402)/Homer | 1e-2 | -5.577e+00 | 0.2237 | 33.0 | 31.43% | 9911.3 | 20.01% | motif file (matrix) | svg |
| 8 | T A G C A G T C T G A C A G T C C T A G A T C G A G T C C A T G T G A C A G T C G T A C A G T C A G T C G C A T C T A G A T C G G C A T A C T G A T C G G A T C | BORIS(Zf)/K562-CTCFL-ChIP-Seq(GSE32465)/Homer | 1e-2 | -5.532e+00 | 0.2237 | 9.0 | 8.57% | 1459.2 | 2.95% | motif file (matrix) | svg |
| 9 | C A T G C T A G T C G A A C G T A C T G C G T A T A G C C G A T T G A C C G T A A G C T G A T C | Fra2(bZIP)/Striatum-Fra2-ChIP-Seq(GSE43429)/Homer | 1e-2 | -5.129e+00 | 0.2725 | 9.0 | 8.57% | 1554.8 | 3.14% | motif file (matrix) | svg |
| 10 | C T G A T C A G C A G T C T A G A C T G C T A G G A T C A T C G A C T G C T G A T C A G G A T C | Sp5(Zf)/mES-Sp5.Flag-ChIP-Seq(GSE72989)/Homer | 1e-2 | -5.056e+00 | 0.2725 | 29.0 | 27.62% | 8638.7 | 17.44% | motif file (matrix) | svg |
| 11 | G T A C C A G T A C T G A C T G A C T G G A T C A C T G A C G T A C T G A C T G A G T C G A T C | KLF6(Zf)/PDAC-KLF6-ChIP-Seq(GSE64557)/Homer | 1e-2 | -4.955e+00 | 0.2725 | 27.0 | 25.71% | 7913.8 | 15.98% | motif file (matrix) | svg |
| 12 | A T G C A G T C C T G A A G T C C G A T A C G T A G T C A G T C A C G T A T C G G A C T A C G T | Etv2(ETS)/ES-ER71-ChIP-Seq(GSE59402)/Homer(0.967) | 1e-2 | -4.793e+00 | 0.2860 | 18.0 | 17.14% | 4618.3 | 9.33% | motif file (matrix) | svg |
| 13 | C G T A A C G T A G C T C G A T C T A G G T A C C G T A A G C T C G T A G C T A | Oct4(POU,Homeobox)/mES-Oct4-ChIP-Seq(GSE11431)/Homer | 1e-2 | -4.787e+00 | 0.2860 | 7.0 | 6.67% | 1080.6 | 2.18% | motif file (matrix) | svg |
| 14 | T G A C C G A T A C T G A C T G A C T G G A C T A C T G A C G T A C T G A C T G G A T C G A T C | EKLF(Zf)/Erythrocyte-Klf1-ChIP-Seq(GSE20478)/Homer | 1e-2 | -4.687e+00 | 0.2860 | 7.0 | 6.67% | 1101.0 | 2.22% | motif file (matrix) | svg |
