## Supplementary material for "Integrative phenotypic and genomic analyses reveal strain-dependent responses to acute ozone exposure and their associations with airway macrophage transcriptional activity": HOMER Motif Analyses: knownResults_GTX_distal_CC003_up.html

./homer/ATAC\_GTX\_CC017\_vs\_CC003\_distal\_negative - Homer Known Motif Enrichment Results


### Homer Known Motif Enrichment Results (./homer/ATAC\_GTX\_CC017\_vs\_CC003\_distal\_negative)

Homer *de novo* Motif Results  
Gene Ontology Enrichment Results  
Known Motif Enrichment Results (txt file)  
Total Target Sequences = 171, Total Background Sequences = 48959

|  |  |  |  |  |  |  |  |  |  |  |  |
| --- | --- | --- | --- | --- | --- | --- | --- | --- | --- | --- | --- |
| Rank | Motif | Name | P-value | log P-pvalue | q-value (Benjamini) | # Target Sequences with Motif | % of Targets Sequences with Motif | # Background Sequences with Motif | % of Background Sequences with Motif | Motif File | SVG |
| 1 | G C T A A G T C T A C G T G C A A T C G T C A G G C T A T C G A T C A G A G C T | ELF5(ETS)/T47D-ELF5-ChIP-Seq(GSE30407)/Homer | 1e-23 | -5.521e+01 | 0.0000 | 60.0 | 35.09% | 3720.9 | 7.60% | motif file (matrix) | svg |
| 2 | C G T A T A C G T C G A A C T G A C T G C G T A C G T A T A C G A G C T T A C G | PU.1(ETS)/ThioMac-PU.1-ChIP-Seq(GSE21512)/Homer | 1e-16 | -3.848e+01 | 0.0000 | 43.0 | 25.15% | 2658.2 | 5.43% | motif file (matrix) | svg |
| 3 | C G T A T G A C T A G C T G C A A C T G A C T G C G T A C G T A T C A G G A C T | ELF3(ETS)/PDAC-ELF3-ChIP-Seq(GSE64557)/Homer | 1e-16 | -3.826e+01 | 0.0000 | 51.0 | 29.82% | 3809.9 | 7.78% | motif file (matrix) | svg |
| 4 | C G T A T A G C T A G C T G C A A C T G C T A G C G T A C G T A T C A G G A C T | EHF(ETS)/LoVo-EHF-ChIP-Seq(GSE49402)/Homer | 1e-16 | -3.706e+01 | 0.0000 | 67.0 | 39.18% | 6637.2 | 13.55% | motif file (matrix) | svg |
| 5 | C T G A A G T C C G A T A G C T A T G C G T A C A C G T A T C G C A G T G C A T | Elf4(ETS)/BMDM-Elf4-ChIP-Seq(GSE88699)/Homer | 1e-14 | -3.297e+01 | 0.0000 | 56.0 | 32.75% | 5182.3 | 10.58% | motif file (matrix) | svg |
| 6 | C G T A C T G A C G T A C T A G T C G A C T A G A C T G C G T A C G T A T A C G A G C T A T C G | SpiB(ETS)/OCILY3-SPIB-ChIP-Seq(GSE56857)/Homer | 1e-13 | -3.071e+01 | 0.0000 | 26.0 | 15.20% | 1136.9 | 2.32% | motif file (matrix) | svg |
| 7 | T C G A T A G C T G C A A C T G A C T G C G T A C G T A C T A G G A C T T A C G | ETS1(ETS)/Jurkat-ETS1-ChIP-Seq(GSE17954)/Homer | 1e-10 | -2.410e+01 | 0.0000 | 49.0 | 28.65% | 5113.9 | 10.44% | motif file (matrix) | svg |
| 8 | T C G A T A G C G T C A A C T G A C T G C G T A C G T A C T A G A G C T T C A G | ERG(ETS)/VCaP-ERG-ChIP-Seq(GSE14097)/Homer | 1e-9 | -2.248e+01 | 0.0000 | 62.0 | 36.26% | 7935.6 | 16.20% | motif file (matrix) | svg |
| 9 | C T A G T C G A A C G T A C T G C G T A A T G C A C G T G T A C C G T A A G C T G A T C G T A C | Atf3(bZIP)/GBM-ATF3-ChIP-Seq(GSE33912)/Homer | 1e-8 | -2.054e+01 | 0.0000 | 36.0 | 21.05% | 3334.7 | 6.81% | motif file (matrix) | svg |
| 10 | C T A G C T A G C G T A C G T A T A C G C G A T C T A G C T G A C T G A C G T A T A C G G A C T | PU.1:IRF8(ETS:IRF)/pDC-Irf8-ChIP-Seq(GSE66899)/Homer | 1e-8 | -2.029e+01 | 0.0000 | 19.0 | 11.11% | 957.6 | 1.96% | motif file (matrix) | svg |
| 11 | T C A G A G C T A T G C C G T A A G C T T C A G C A G T A C T G C T G A A G T C | MITF(bHLH)/MastCells-MITF-ChIP-Seq(GSE48085)/Homer | 1e-8 | -1.993e+01 | 0.0000 | 42.0 | 24.56% | 4459.2 | 9.10% | motif file (matrix) | svg |
| 12 | T C A G T C A G G C T A C G T A T A C G G A C T T C A G T C G A C T G A C G T A T A C G G A C T | IRF8(IRF)/BMDM-IRF8-ChIP-Seq(GSE77884)/Homer | 1e-8 | -1.940e+01 | 0.0000 | 24.0 | 14.04% | 1638.3 | 3.35% | motif file (matrix) | svg |
| 13 | A G T C C T G A A G T C C G A T C A G T G A T C A T G C A C T G A T C G G A C T | Fli1(ETS)/CD8-FLI-ChIP-Seq(GSE20898)/Homer | 1e-8 | -1.857e+01 | 0.0000 | 43.0 | 25.15% | 4858.5 | 9.92% | motif file (matrix) | svg |
| 14 | A C T G C T A G T C G A C G A T C A T G G C T A A T C G C G A T G T A C G C T A A G C T G T A C | Fra1(bZIP)/BT549-Fra1-ChIP-Seq(GSE46166)/Homer | 1e-7 | -1.638e+01 | 0.0000 | 29.0 | 16.96% | 2708.0 | 5.53% | motif file (matrix) | svg |
| 15 | C A G T T G C A A C G T A C T G C G T A A T C G C G A T T G A C C G T A A C G T | BATF(bZIP)/Th17-BATF-ChIP-Seq(GSE39756)/Homer | 1e-7 | -1.620e+01 | 0.0000 | 33.0 | 19.30% | 3413.1 | 6.97% | motif file (matrix) | svg |
| 16 | T C G A T C G A T A G C G T A C T C A G T A C G C G T A C G T A T C A G A G C T | GABPA(ETS)/Jurkat-GABPa-ChIP-Seq(GSE17954)/Homer | 1e-6 | -1.471e+01 | 0.0000 | 36.0 | 21.05% | 4200.2 | 8.58% | motif file (matrix) | svg |
| 17 | T C G A A C G T C A T G G C T A T A G C C G A T G T A C G C T A A C G T A T G C | AP-1(bZIP)/ThioMac-PU.1-ChIP-Seq(GSE21512)/Homer | 1e-6 | -1.402e+01 | 0.0000 | 33.0 | 19.30% | 3759.2 | 7.68% | motif file (matrix) | svg |
| 18 | A T G C A G T C C T G A A G T C C G A T A C G T A G T C A G T C A C G T A T C G G A C T A C G T | Etv2(ETS)/ES-ER71-ChIP-Seq(GSE59402)/Homer(0.967) | 1e-6 | -1.388e+01 | 0.0000 | 37.0 | 21.64% | 4542.6 | 9.28% | motif file (matrix) | svg |
| 19 | T C G A A C G T A C G T C T G A G A T C T C A G G A C T G T C A C G T A A G C T G T C A C T A G A G C T A C G T T C G A | NFIL3(bZIP)/HepG2-NFIL3-ChIP-Seq(Encode)/Homer | 1e-5 | -1.363e+01 | 0.0000 | 26.0 | 15.20% | 2577.7 | 5.26% | motif file (matrix) | svg |
| 20 | T C G A C T G A T A G C T G A C T C A G T C A G C G T A C G T A T C A G A G C T | ETV1(ETS)/GIST48-ETV1-ChIP-Seq(GSE22441)/Homer | 1e-5 | -1.344e+01 | 0.0000 | 47.0 | 27.49% | 6680.8 | 13.64% | motif file (matrix) | svg |
| 21 | T C A G G A C T C A G T C T G A A G C T C T A G G A C T T G C A C T G A A G T C | HLF(bZIP)/HSC-HLF.Flag-ChIP-Seq(GSE69817)/Homer | 1e-5 | -1.242e+01 | 0.0001 | 31.0 | 18.13% | 3667.6 | 7.49% | motif file (matrix) | svg |
| 22 | C T A G T C G A G C A T C A T G G C T A T A G C C G A T G T A C C T G A A G C T | JunB(bZIP)/DendriticCells-Junb-ChIP-Seq(GSE36099)/Homer | 1e-5 | -1.236e+01 | 0.0001 | 26.0 | 15.20% | 2765.4 | 5.65% | motif file (matrix) | svg |
| 23 | C A T G C T A G T C G A A C G T A C T G C G T A T A G C C G A T T G A C C G T A A G C T G A T C | Fra2(bZIP)/Striatum-Fra2-ChIP-Seq(GSE43429)/Homer | 1e-5 | -1.171e+01 | 0.0001 | 23.0 | 13.45% | 2343.3 | 4.78% | motif file (matrix) | svg |
| 24 | C T A G T C G A C G A T C T A G G C A T C A G T C T A G G A T C C G T A G T C A | CEBP:AP1(bZIP)/ThioMac-CEBPb-ChIP-Seq(GSE21512)/Homer | 1e-4 | -1.101e+01 | 0.0003 | 28.0 | 16.37% | 3360.3 | 6.86% | motif file (matrix) | svg |
| 25 | T C A G A C G T A G T C T C G A A G T C T C A G G C A T C T A G C T A G A G C T | Usf2(bHLH)/C2C12-Usf2-ChIP-Seq(GSE36030)/Homer | 1e-4 | -1.024e+01 | 0.0006 | 16.0 | 9.36% | 1395.6 | 2.85% | motif file (matrix) | svg |
| 26 | G C T A C T G A T C G A A G T C A G T C C T G A A G T C G T C A C T G A T G C A | RUNX1(Runt)/Jurkat-RUNX1-ChIP-Seq(GSE29180)/Homer | 1e-4 | -9.980e+00 | 0.0007 | 38.0 | 22.22% | 5612.0 | 11.46% | motif file (matrix) | svg |
| 27 | C T A G T C G A C G A T A C T G C G T A T A C G A G C T T G A C G C T A A C G T G A T C T A G C | Fosl2(bZIP)/3T3L1-Fosl2-ChIP-Seq(GSE56872)/Homer | 1e-4 | -9.972e+00 | 0.0007 | 17.0 | 9.94% | 1586.6 | 3.24% | motif file (matrix) | svg |
| 28 | T G C A T C G A T A G C G T A C T C A G C T A G G T C A G C T A T C A G G A C T | ETS(ETS)/Promoter/Homer | 1e-4 | -9.909e+00 | 0.0007 | 15.0 | 8.77% | 1280.4 | 2.61% | motif file (matrix) | svg |
| 29 | C G A T T A C G T G A C G A C T C A T G C G T A T A C G A C G T G T A C C T G A | Bach2(bZIP)/OCILy7-Bach2-ChIP-Seq(GSE44420)/Homer | 1e-4 | -9.255e+00 | 0.0014 | 12.0 | 7.02% | 908.6 | 1.86% | motif file (matrix) | svg |
| 30 | C T G A T G C A T A G C T G A C T A C G T C A G C T G A G C T A T C A G G A C T | ELF1(ETS)/Jurkat-ELF1-ChIP-Seq(SRA014231)/Homer | 1e-3 | -9.166e+00 | 0.0014 | 19.0 | 11.11% | 2038.1 | 4.16% | motif file (matrix) | svg |
| 31 | A C G T T A C G G A T C A C T G A C G T C T A G A C T G A C T G G A T C C T A G C A T G C T A G | Egr2(Zf)/Thymocytes-Egr2-ChIP-Seq(GSE34254)/Homer | 1e-3 | -8.855e+00 | 0.0019 | 9.0 | 5.26% | 544.0 | 1.11% | motif file (matrix) | svg |
| 32 | T C G A A G C T A C G T A C G T A G T C A G T C A C G T A T C G G A C T A T C G | EWS:ERG-fusion(ETS)/CADO\_ES1-EWS:ERG-ChIP-Seq(SRA014231)/Homer | 1e-3 | -8.505e+00 | 0.0026 | 26.0 | 15.20% | 3481.1 | 7.11% | motif file (matrix) | svg |
| 33 | C T G A T A C G G C A T A G C T A G C T A G T C T C G A A C T G C A G T A G C T A G C T G A T C | IRF3(IRF)/BMDM-Irf3-ChIP-Seq(GSE67343)/Homer | 1e-3 | -8.447e+00 | 0.0027 | 16.0 | 9.36% | 1631.2 | 3.33% | motif file (matrix) | svg |
| 34 | T A G C C T A G T C G A G A C T A C T G C T G A A G T C T C A G G C A T T G A C C T G A A G C T | Atf7(bZIP)/3T3L1-Atf7-ChIP-Seq(GSE56872)/Homer | 1e-3 | -8.146e+00 | 0.0035 | 18.0 | 10.53% | 2028.5 | 4.14% | motif file (matrix) | svg |
| 35 | C T A G T C A G C A G T T C A G A C T G A C T G G A T C C T A G A C T G C T A G T C A G A T G C | KLF14(Zf)/HEK293-KLF14.GFP-ChIP-Seq(GSE58341)/Homer | 1e-3 | -8.076e+00 | 0.0037 | 38.0 | 22.22% | 6151.8 | 12.56% | motif file (matrix) | svg |
| 36 | A T G C A T G C A T C G T A C G A G C T A G T C G C T A A G T C T C A G G A C T A C T G T C G A | E-box(bHLH)/Promoter/Homer | 1e-3 | -8.015e+00 | 0.0038 | 6.0 | 3.51% | 260.0 | 0.53% | motif file (matrix) | svg |
| 37 | C T G A T A G C T G A C T C A G C T A G G T C A C G T A T C A G A G C T T C A G | ETV4(ETS)/HepG2-ETV4-ChIP-Seq(ENCODE)/Homer | 1e-3 | -7.550e+00 | 0.0059 | 32.0 | 18.71% | 4981.0 | 10.17% | motif file (matrix) | svg |
| 38 | T G C A A G C T C A T G C G T A A G C T A C T G G A T C G T C A C G T A A G C T | Atf4(bZIP)/MEF-Atf4-ChIP-Seq(GSE35681)/Homer | 1e-3 | -7.495e+00 | 0.0061 | 12.0 | 7.02% | 1104.0 | 2.25% | motif file (matrix) | svg |
| 39 | T A C G T C G A G A C T A C T G C T G A A G T C T C A G G A C T T G A C C T G A | Atf1(bZIP)/K562-ATF1-ChIP-Seq(GSE31477)/Homer | 1e-3 | -7.254e+00 | 0.0075 | 22.0 | 12.87% | 2964.0 | 6.05% | motif file (matrix) | svg |
| 40 | C T A G T C G A A C G T A C T G C G T A T A G C C G A T G T A C C G T A A G C T G A T C G T A C | Jun-AP1(bZIP)/K562-cJun-ChIP-Seq(GSE31477)/Homer | 1e-3 | -7.248e+00 | 0.0075 | 12.0 | 7.02% | 1136.7 | 2.32% | motif file (matrix) | svg |
| 41 | T C G A A C T G A C T G C G T A C G T A T C G A A G T C C T G A A T C G G T A C G C A T C A T G | ETS:E-box(ETS,bHLH)/HPC7-Scl-ChIP-Seq(GSE22178)/Homer | 1e-3 | -7.233e+00 | 0.0075 | 7.0 | 4.09% | 417.9 | 0.85% | motif file (matrix) | svg |
| 42 | G T A C G A T C C A G T A G T C A G T C A G T C T G C A G A T C C T G A A T G C G T C A A C G T | WT1(Zf)/Kidney-WT1-ChIP-Seq(GSE90016)/Homer | 1e-3 | -7.229e+00 | 0.0075 | 17.0 | 9.94% | 2008.1 | 4.10% | motif file (matrix) | svg |
| 43 | A C T G G A T C G A C T A C T G A C G T C A T G A C T G A C G T A G C T C G A T | RUNX-AML(Runt)/CD4+-PolII-ChIP-Seq(Barski\_et\_al.)/Homer | 1e-3 | -7.088e+00 | 0.0080 | 25.0 | 14.62% | 3615.7 | 7.38% | motif file (matrix) | svg |
| 44 | T A G C G C T A T C G A C T G A A G T C A G T C C T G A A G T C C G T A C T A G | RUNX(Runt)/HPC7-Runx1-ChIP-Seq(GSE22178)/Homer | 1e-3 | -6.975e+00 | 0.0088 | 26.0 | 15.20% | 3854.4 | 7.87% | motif file (matrix) | svg |
| 45 | T C A G A G C T A T G C C G T A A G T C T C A G A C G T A T C G T C G A A G T C G A T C T G A C | TFE3(bHLH)/MEF-TFE3-ChIP-Seq(GSE75757)/Homer | 1e-2 | -6.763e+00 | 0.0106 | 6.0 | 3.51% | 330.8 | 0.68% | motif file (matrix) | svg |
| 46 | T A C G T C A G A G C T A T G C C G T A A G T C T C A G A C G T A C T G T C G A | USF1(bHLH)/GM12878-Usf1-ChIP-Seq(GSE32465)/Homer | 1e-2 | -6.563e+00 | 0.0127 | 16.0 | 9.36% | 1947.1 | 3.98% | motif file (matrix) | svg |
| 47 | T G C A A T G C A C G T A C G T A C G T A T G C C T A G A C G T A C G T A G C T G A T C A G C T | T1ISRE(IRF)/ThioMac-Ifnb-Expression/Homer | 1e-2 | -6.513e+00 | 0.0131 | 3.0 | 1.75% | 63.3 | 0.13% | motif file (matrix) | svg |
| 48 | T G C A C T G A A T G C G T C A A C T G A C T G C G T A C G T A C T A G A G C T | Ets1-distal(ETS)/CD4+-PolII-ChIP-Seq(Barski\_et\_al.)/Homer | 1e-2 | -6.219e+00 | 0.0172 | 13.0 | 7.60% | 1460.0 | 2.98% | motif file (matrix) | svg |
| 49 | T G C A A G C T A C G T C T A G G A T C C T A G G A T C G T C A C T G A A G T C | CEBP(bZIP)/ThioMac-CEBPb-ChIP-Seq(GSE21512)/Homer | 1e-2 | -6.202e+00 | 0.0172 | 20.0 | 11.70% | 2809.0 | 5.74% | motif file (matrix) | svg |
| 50 | C T A G C A G T T G A C C G T A G A T C T C A G G A C T C A T G | BMAL1(bHLH)/Liver-Bmal1-ChIP-Seq(GSE39860)/Homer | 1e-2 | -5.946e+00 | 0.0217 | 53.0 | 30.99% | 10569.4 | 21.58% | motif file (matrix) | svg |
| 51 | C G T A C T A G A C T G A C T G G A C T C T A G C A G T C T A G C A T G G A T C | KLF5(Zf)/LoVo-KLF5-ChIP-Seq(GSE49402)/Homer | 1e-2 | -5.902e+00 | 0.0222 | 28.0 | 16.37% | 4604.8 | 9.40% | motif file (matrix) | svg |
| 52 | T A C G T C G A C A G T A C T G G C T A A T G C C G A T G T A C C G T A A C T G T A G C C G T A | NF-E2(bZIP)/K562-NFE2-ChIP-Seq(GSE31477)/Homer | 1e-2 | -5.698e+00 | 0.0267 | 5.0 | 2.92% | 284.0 | 0.58% | motif file (matrix) | svg |
| 53 | G T C A G C A T G C T A C A G T C T A G G A T C C G T A C T G A C G T A C G A T | Oct2(POU,Homeobox)/Bcell-Oct2-ChIP-Seq(GSE21512)/Homer | 1e-2 | -5.577e+00 | 0.0296 | 11.0 | 6.43% | 1213.0 | 2.48% | motif file (matrix) | svg |
| 54 | G T C A T G C A G C T A A G T C C G T A A C T G T G A C G C A T T C A G C A G T | Ap4(bHLH)/AML-Tfap4-ChIP-Seq(GSE45738)/Homer | 1e-2 | -5.452e+00 | 0.0329 | 32.0 | 18.71% | 5674.1 | 11.59% | motif file (matrix) | svg |
| 55 | T C G A G C A T A C T G C T G A A G T C T C A G G A C T G T A C C G T A A G C T A G T C G A T C | c-Jun-CRE(bZIP)/K562-cJun-ChIP-Seq(GSE31477)/Homer | 1e-2 | -5.033e+00 | 0.0491 | 11.0 | 6.43% | 1306.3 | 2.67% | motif file (matrix) | svg |
| 56 | C A T G T G A C C G T A A G T C T A C G G C A T A C T G G T C A A T G C A G T C | bHLHE41(bHLH)/proB-Bhlhe41-ChIP-Seq(GSE93764)/Homer | 1e-2 | -4.946e+00 | 0.0526 | 28.0 | 16.37% | 4938.5 | 10.08% | motif file (matrix) | svg |
| 57 | A T G C A T C G T A C G A G C T A T C G C T G A A G T C C T A G A G C T A T G C C T G A A T G C | CRE(bZIP)/Promoter/Homer | 1e-2 | -4.896e+00 | 0.0543 | 8.0 | 4.68% | 800.9 | 1.64% | motif file (matrix) | svg |
| 58 | C G A T T A C G T G C A G T A C G A T C G A C T A G C T A C G T A T C G G T A C G A T C G T A C G A T C G T C A | PPARE(NR),DR1/3T3L1-Pparg-ChIP-Seq(GSE13511)/Homer | 1e-2 | -4.808e+00 | 0.0583 | 25.0 | 14.62% | 4301.6 | 8.78% | motif file (matrix) | svg |
| 59 | T G A C C T A G T C A G G T C A C G T A T C A G C G A T T C A G T C G A T G C A C T G A T A G C | PU.1-IRF(ETS:IRF)/Bcell-PU.1-ChIP-Seq(GSE21512)/Homer | 1e-2 | -4.805e+00 | 0.0583 | 38.0 | 22.22% | 7379.8 | 15.07% | motif file (matrix) | svg |
