## Supplementary material for "Integrative phenotypic and genomic analyses reveal strain-dependent responses to acute ozone exposure and their associations with airway macrophage transcriptional activity": HOMER Motif Analyses: knownResults_GTX_distal_CC017_up.html

./homer/ATAC\_GTX\_CC017\_vs\_CC003\_distal\_positive - Homer Known Motif Enrichment Results


### Homer Known Motif Enrichment Results (./homer/ATAC\_GTX\_CC017\_vs\_CC003\_distal\_positive)

Homer *de novo* Motif Results  
Gene Ontology Enrichment Results  
Known Motif Enrichment Results (txt file)  
Total Target Sequences = 13, Total Background Sequences = 47170

|  |  |  |  |  |  |  |  |  |  |  |  |
| --- | --- | --- | --- | --- | --- | --- | --- | --- | --- | --- | --- |
| Rank | Motif | Name | P-value | log P-pvalue | q-value (Benjamini) | # Target Sequences with Motif | % of Targets Sequences with Motif | # Background Sequences with Motif | % of Background Sequences with Motif | Motif File | SVG |
| 1 | C A T G C T A G T C G A A C G T A C T G C G T A T A G C C G A T T G A C C G T A A G C T G A T C | Fra2(bZIP)/Striatum-Fra2-ChIP-Seq(GSE43429)/Homer | 1e-3 | -8.510e+00 | 0.0834 | 4.0 | 30.77% | 1136.8 | 2.41% | motif file (matrix) | svg |
| 2 | A C T G C T A G T C G A C G A T C A T G G C T A A T C G C G A T G T A C G C T A A G C T G T A C | Fra1(bZIP)/BT549-Fra1-ChIP-Seq(GSE46166)/Homer | 1e-3 | -8.329e+00 | 0.0834 | 4.0 | 30.77% | 1191.1 | 2.52% | motif file (matrix) | svg |
| 3 | C T G A A G T C C G A T A G C T A T G C G T A C A C G T A T C G C A G T G C A T | Elf4(ETS)/BMDM-Elf4-ChIP-Seq(GSE88699)/Homer | 1e-3 | -8.304e+00 | 0.0834 | 7.0 | 53.85% | 5446.1 | 11.54% | motif file (matrix) | svg |
| 4 | C A G T T G C A A C G T A C T G C G T A A T C G C G A T T G A C C G T A A C G T | BATF(bZIP)/Th17-BATF-ChIP-Seq(GSE39756)/Homer | 1e-3 | -7.627e+00 | 0.0834 | 4.0 | 30.77% | 1433.3 | 3.04% | motif file (matrix) | svg |
| 5 | C T A G T C G A A C G T A C T G C G T A A T G C A C G T G T A C C G T A A G C T G A T C G T A C | Atf3(bZIP)/GBM-ATF3-ChIP-Seq(GSE33912)/Homer | 1e-3 | -7.528e+00 | 0.0834 | 4.0 | 30.77% | 1471.6 | 3.12% | motif file (matrix) | svg |
| 6 | T C G A A C G T C A T G G C T A T A G C C G A T G T A C G C T A A C G T A T G C | AP-1(bZIP)/ThioMac-PU.1-ChIP-Seq(GSE21512)/Homer | 1e-3 | -6.919e+00 | 0.0834 | 4.0 | 30.77% | 1730.6 | 3.67% | motif file (matrix) | svg |
| 7 | T G C A A G C T C T G A A T C G G A C T C T A G G T A C G A T C G T C A A G T C G T A C G A C T C T A G A T C G G C A T C A T G C A T G G A T C G T A C C T G A | CTCF(Zf)/CD4+-CTCF-ChIP-Seq(Barski\_et\_al.)/Homer | 1e-2 | -6.675e+00 | 0.0834 | 3.0 | 23.08% | 808.0 | 1.71% | motif file (matrix) | svg |
| 8 | C T A G T C G A C G A T A C T G C G T A T A C G A G C T T G A C G C T A A C G T G A T C T A G C | Fosl2(bZIP)/3T3L1-Fosl2-ChIP-Seq(GSE56872)/Homer | 1e-2 | -6.554e+00 | 0.0834 | 3.0 | 23.08% | 843.3 | 1.79% | motif file (matrix) | svg |
| 9 | T C G A T C G A T A G C G T A C T C A G T A C G C G T A C G T A T C A G A G C T | GABPA(ETS)/Jurkat-GABPa-ChIP-Seq(GSE17954)/Homer | 1e-2 | -6.430e+00 | 0.0834 | 6.0 | 46.15% | 5246.4 | 11.12% | motif file (matrix) | svg |
| 10 | T A G C A G T C T G A C A G T C C T A G A T C G A G T C C A T G T G A C A G T C G T A C A G T C A G T C G C A T C T A G A T C G G C A T A C T G A T C G G A T C | BORIS(Zf)/K562-CTCFL-ChIP-Seq(GSE32465)/Homer | 1e-2 | -6.355e+00 | 0.0834 | 4.0 | 30.77% | 2014.8 | 4.27% | motif file (matrix) | svg |
| 11 | T C G A T A G C G T C A A C T G A C T G C G T A C G T A C T A G A G C T T C A G | ERG(ETS)/VCaP-ERG-ChIP-Seq(GSE14097)/Homer | 1e-2 | -6.227e+00 | 0.0834 | 7.0 | 53.85% | 7617.9 | 16.14% | motif file (matrix) | svg |
| 12 | C A T G G A C T G C A T A C T G A G C T A C T G A C T G C G T A G C A T A G C T A T C G T A C G | Foxh1(Forkhead)/hESC-FOXH1-ChIP-Seq(GSE29422)/Homer | 1e-2 | -5.549e+00 | 0.1342 | 3.0 | 23.08% | 1201.1 | 2.54% | motif file (matrix) | svg |
| 13 | C T A G T C G A G C A T C A T G G C T A T A G C C G A T G T A C C T G A A G C T | JunB(bZIP)/DendriticCells-Junb-ChIP-Seq(GSE36099)/Homer | 1e-2 | -5.442e+00 | 0.1380 | 3.0 | 23.08% | 1248.8 | 2.65% | motif file (matrix) | svg |
| 14 | A T G C A G T C C T G A A G T C C G A T A C G T A G T C A G T C A C G T A T C G G A C T A C G T | Etv2(ETS)/ES-ER71-ChIP-Seq(GSE59402)/Homer(0.967) | 1e-2 | -5.349e+00 | 0.1406 | 5.0 | 38.46% | 4396.8 | 9.32% | motif file (matrix) | svg |
| 15 | C G T A C T G A A C T G A G C T A G T C G A T C G A T C G C A T C T G A C T A G C T A G T A C G T C G A T G C A G C A T | EBF2(EBF)/BrownAdipose-EBF2-ChIP-Seq(GSE97114)/Homer | 1e-2 | -5.159e+00 | 0.1586 | 5.0 | 38.46% | 4593.7 | 9.73% | motif file (matrix) | svg |
| 16 | T G C A C T G A A G T C G T C A A C T G A C T G C G T A C G T A C T G A A G C T | EWS:FLI1-fusion(ETS)/SK\_N\_MC-EWS:FLI1-ChIP-Seq(SRA014231)/Homer | 1e-2 | -5.037e+00 | 0.1680 | 4.0 | 30.77% | 2899.3 | 6.14% | motif file (matrix) | svg |
| 17 | G A C T T C A G C T A G A G T C A G T C G T A C A G T C C T G A A G T C A G T C A G T C G A C T A G T C A C T G A T G C | KLF3(Zf)/MEF-Klf3-ChIP-Seq(GSE44748)/Homer | 1e-2 | -4.757e+00 | 0.2093 | 6.0 | 46.15% | 7270.7 | 15.40% | motif file (matrix) | svg |
| 18 | G C T A A G T C T A C G T G C A A T C G T C A G G C T A T C G A T C A G A G C T | ELF5(ETS)/T47D-ELF5-ChIP-Seq(GSE30407)/Homer | 1e-2 | -4.673e+00 | 0.2149 | 4.0 | 30.77% | 3215.5 | 6.81% | motif file (matrix) | svg |
