## Supplementary material for "Integrative phenotypic and genomic analyses reveal strain-dependent responses to acute ozone exposure and their associations with airway macrophage transcriptional activity": HOMER Motif Analyses: knownResults_GTX_proximal_CC003_up.html

./homer/ATAC\_GTX\_CC017\_vs\_CC003\_promoter\_negative - Homer Known Motif Enrichment Results


### Homer Known Motif Enrichment Results (./homer/ATAC\_GTX\_CC017\_vs\_CC003\_promoter\_negative)

Homer *de novo* Motif Results  
Gene Ontology Enrichment Results  
Known Motif Enrichment Results (txt file)  
Total Target Sequences = 19, Total Background Sequences = 47377

|  |  |  |  |  |  |  |  |  |  |  |  |
| --- | --- | --- | --- | --- | --- | --- | --- | --- | --- | --- | --- |
| Rank | Motif | Name | P-value | log P-pvalue | q-value (Benjamini) | # Target Sequences with Motif | % of Targets Sequences with Motif | # Background Sequences with Motif | % of Background Sequences with Motif | Motif File | SVG |
| 1 | T C A G T C A G G C T A C G T A T A C G G A C T T C A G T C G A C T G A C G T A T A C G G A C T | IRF8(IRF)/BMDM-IRF8-ChIP-Seq(GSE77884)/Homer | 1e-4 | -1.066e+01 | 0.0097 | 6.0 | 31.58% | 1558.7 | 3.29% | motif file (matrix) | svg |
| 2 | T C G A A C G T C A T G G C T A T A G C C G A T G T A C G C T A A C G T A T G C | AP-1(bZIP)/ThioMac-PU.1-ChIP-Seq(GSE21512)/Homer | 1e-3 | -8.857e+00 | 0.0295 | 7.0 | 36.84% | 3157.3 | 6.66% | motif file (matrix) | svg |
| 3 | T C A G C T G A C G T A C G T A T A C G G C A T C T A G C T G A C G T A C G T A T A C G G A C T | IRF1(IRF)/PBMC-IRF1-ChIP-Seq(GSE43036)/Homer | 1e-3 | -8.769e+00 | 0.0295 | 4.0 | 21.05% | 702.7 | 1.48% | motif file (matrix) | svg |
| 4 | C T G A T A C G G C A T A G C T A G C T A G T C T C G A A C T G C A G T A G C T A G C T G A T C | IRF3(IRF)/BMDM-Irf3-ChIP-Seq(GSE67343)/Homer | 1e-3 | -8.309e+00 | 0.0295 | 5.0 | 26.32% | 1491.9 | 3.14% | motif file (matrix) | svg |
| 5 | C T A G C T A G C G T A C G T A T A C G C G A T C T A G C T G A C T G A C G T A T A C G G A C T | PU.1:IRF8(ETS:IRF)/pDC-Irf8-ChIP-Seq(GSE66899)/Homer | 1e-3 | -8.001e+00 | 0.0295 | 4.0 | 21.05% | 859.1 | 1.81% | motif file (matrix) | svg |
| 6 | C T A G T C G A G C A T C A T G G C T A T A G C C G A T G T A C C T G A A G C T | JunB(bZIP)/DendriticCells-Junb-ChIP-Seq(GSE36099)/Homer | 1e-2 | -6.420e+00 | 0.1124 | 5.0 | 26.32% | 2261.1 | 4.77% | motif file (matrix) | svg |
| 7 | G C T A A G T C T A C G T G C A A T C G T C A G G C T A T C G A T C A G A G C T | ELF5(ETS)/T47D-ELF5-ChIP-Seq(GSE30407)/Homer | 1e-2 | -6.179e+00 | 0.1226 | 6.0 | 31.58% | 3565.9 | 7.52% | motif file (matrix) | svg |
| 8 | C T G A A G T C C G A T A G C T A T G C G T A C A C G T A T C G C A G T G C A T | Elf4(ETS)/BMDM-Elf4-ChIP-Seq(GSE88699)/Homer | 1e-2 | -5.990e+00 | 0.1296 | 7.0 | 36.84% | 5072.7 | 10.69% | motif file (matrix) | svg |
| 9 | C A G T T G C A A C G T A C T G C G T A A T C G C G A T T G A C C G T A A C G T | BATF(bZIP)/Th17-BATF-ChIP-Seq(GSE39756)/Homer | 1e-2 | -5.384e+00 | 0.2111 | 5.0 | 26.32% | 2868.0 | 6.05% | motif file (matrix) | svg |
| 10 | C A T G G T A C G A C T G C T A C G T A C G T A C G T A G C T A G A C T C T G A T C A G G T A C | Mef2c(MADS)/GM12878-Mef2c-ChIP-Seq(GSE32465)/Homer | 1e-2 | -5.342e+00 | 0.2111 | 4.0 | 21.05% | 1770.4 | 3.73% | motif file (matrix) | svg |
| 11 | G C T A T A G C A G C T A T C G G T C A C G T A G C T A A T G C G A T C C T G A | IRF4(IRF)/GM12878-IRF4-ChIP-Seq(GSE32465)/Homer | 1e-2 | -4.924e+00 | 0.2736 | 4.0 | 21.05% | 1993.0 | 4.20% | motif file (matrix) | svg |
