## Supplementary material for "Integrative phenotypic and genomic analyses reveal strain-dependent responses to acute ozone exposure and their associations with airway macrophage transcriptional activity": HOMER Motif Analyses: knownResults_GTX_proximal_CC017_up.html

./homer/ATAC\_GTX\_CC017\_vs\_CC003\_promoter\_positive - Homer Known Motif Enrichment Results


### Homer Known Motif Enrichment Results (./homer/ATAC\_GTX\_CC017\_vs\_CC003\_promoter\_positive)

Homer *de novo* Motif Results  
Gene Ontology Enrichment Results  
Known Motif Enrichment Results (txt file)  
Total Target Sequences = 132, Total Background Sequences = 45015

|  |  |  |  |  |  |  |  |  |  |  |  |
| --- | --- | --- | --- | --- | --- | --- | --- | --- | --- | --- | --- |
| Rank | Motif | Name | P-value | log P-pvalue | q-value (Benjamini) | # Target Sequences with Motif | % of Targets Sequences with Motif | # Background Sequences with Motif | % of Background Sequences with Motif | Motif File | SVG |
| 1 | A T G C A G C T T C A G T G A C T C A G A T G C T G C A A C G T A T C G G A T C A C T G A G T C | NRF1(NRF)/MCF7-NRF1-ChIP-Seq(Unpublished)/Homer | 1e-8 | -1.857e+01 | 0.0000 | 39.0 | 29.55% | 5037.2 | 11.19% | motif file (matrix) | svg |
| 2 | A T C G A G C T A C T G A G T C A C T G A G T C C G T A A C G T A C T G A G T C A C T G A G T C | NRF(NRF)/Promoter/Homer | 1e-7 | -1.638e+01 | 0.0000 | 34.0 | 25.76% | 4333.1 | 9.63% | motif file (matrix) | svg |
| 3 | C A T G T G A C C G T A A G T C T A C G G C A T A C T G G T C A A T G C A G T C | bHLHE41(bHLH)/proB-Bhlhe41-ChIP-Seq(GSE93764)/Homer | 1e-3 | -8.840e+00 | 0.0200 | 48.0 | 36.36% | 9972.2 | 22.16% | motif file (matrix) | svg |
| 4 | A T G C G A T C G A C T A G C T C G A T C G A T G T C A C G A T T C G A A T C G T A G C T A G C | TATA-Box(TBP)/Promoter/Homer | 1e-3 | -7.636e+00 | 0.0499 | 16.0 | 12.12% | 2109.2 | 4.69% | motif file (matrix) | svg |
| 5 | G A T C T C G A A G T C C G A T C G A T A G T C A T G C A C T G A T C G G A C T | Elk1(ETS)/Hela-Elk1-ChIP-Seq(GSE31477)/Homer | 1e-3 | -7.479e+00 | 0.0499 | 34.0 | 25.76% | 6576.4 | 14.61% | motif file (matrix) | svg |
| 6 | G A T C C T G A A G T C C G A T C G A T G A T C A G T C A C T G A T C G A G C T | Elk4(ETS)/Hela-Elk4-ChIP-Seq(GSE31477)/Homer | 1e-3 | -7.393e+00 | 0.0499 | 34.0 | 25.76% | 6607.4 | 14.68% | motif file (matrix) | svg |
| 7 | C T G A T G C A T A G C T G A C T A C G T C A G C T G A G C T A T C A G G A C T | ELF1(ETS)/Jurkat-ELF1-ChIP-Seq(SRA014231)/Homer | 1e-3 | -6.938e+00 | 0.0574 | 31.0 | 23.48% | 5964.2 | 13.25% | motif file (matrix) | svg |
| 8 | A T G C T C A G T C G A G C A T A C T G C G T A A G T C T C A G G A C T T G A C C G T A A G C T | Atf2(bZIP)/3T3L1-Atf2-ChIP-Seq(GSE56872)/Homer | 1e-2 | -6.708e+00 | 0.0632 | 11.0 | 8.33% | 1252.8 | 2.78% | motif file (matrix) | svg |
| 9 | T C G A T A G C T G C A A C T G A C T G C G T A C G T A C T A G G A C T T A C G | ETS1(ETS)/Jurkat-ETS1-ChIP-Seq(GSE17954)/Homer | 1e-2 | -6.192e+00 | 0.0941 | 29.0 | 21.97% | 5702.6 | 12.67% | motif file (matrix) | svg |
| 10 | T C G A T C G A T A G C G T A C T C A G T A C G C G T A C G T A T C A G A G C T | GABPA(ETS)/Jurkat-GABPa-ChIP-Seq(GSE17954)/Homer | 1e-2 | -6.012e+00 | 0.1014 | 29.0 | 21.97% | 5771.1 | 12.82% | motif file (matrix) | svg |
| 11 | C T G A T A G C T G A C T C A G C T A G G T C A C G T A T C A G A G C T T C A G | ETV4(ETS)/HepG2-ETV4-ChIP-Seq(ENCODE)/Homer | 1e-2 | -6.008e+00 | 0.1014 | 40.0 | 30.30% | 8874.2 | 19.72% | motif file (matrix) | svg |
| 12 | C T A G T C G A G C A T C A T G G C T A T A G C C G A T G T A C C T G A A G C T | JunB(bZIP)/DendriticCells-Junb-ChIP-Seq(GSE36099)/Homer | 1e-2 | -5.979e+00 | 0.1014 | 7.0 | 5.30% | 622.5 | 1.38% | motif file (matrix) | svg |
| 13 | C G T A C T A G A C T G A C T G G A C T C T A G C A G T C T A G C A T G G A T C | KLF5(Zf)/LoVo-KLF5-ChIP-Seq(GSE49402)/Homer | 1e-2 | -5.956e+00 | 0.1014 | 69.0 | 52.27% | 17939.9 | 39.86% | motif file (matrix) | svg |
| 14 | C G T A A C T G C G T A A C G T A T C G C A G T T A G C C G T A T C G A G T A C C T G A T A G C C G T A A C T G C G T A A C G T C G T A C T G A A T C G G C T A | GATA3(Zf),DR8/iTreg-Gata3-ChIP-Seq(GSE20898)/Homer | 1e-2 | -5.685e+00 | 0.1014 | 2.0 | 1.52% | 29.1 | 0.06% | motif file (matrix) | svg |
| 15 | T A C G C T A G A T G C G A T C G T A C A G T C C T A G A G T C A G T C A G T C G T A C A G T C | Sp1(Zf)/Promoter/Homer | 1e-2 | -5.577e+00 | 0.1045 | 42.0 | 31.82% | 9676.3 | 21.50% | motif file (matrix) | svg |
| 16 | A G T C C T G A A G T C C G A T C A G T G A T C A T G C A C T G A T C G G A C T | Fli1(ETS)/CD8-FLI-ChIP-Seq(GSE20898)/Homer | 1e-2 | -5.538e+00 | 0.1045 | 36.0 | 27.27% | 7937.4 | 17.64% | motif file (matrix) | svg |
| 17 | C G T A C T G A C G T A C T A G T C G A C T A G A C T G C G T A C G T A T A C G A G C T A T C G | SpiB(ETS)/OCILY3-SPIB-ChIP-Seq(GSE56857)/Homer | 1e-2 | -5.316e+00 | 0.1196 | 7.0 | 5.30% | 703.6 | 1.56% | motif file (matrix) | svg |
| 18 | T A G C C T A G T C G A G A C T A C T G C T G A A G T C T C A G G C A T T G A C C T G A A G C T | Atf7(bZIP)/3T3L1-Atf7-ChIP-Seq(GSE56872)/Homer | 1e-2 | -5.301e+00 | 0.1196 | 12.0 | 9.09% | 1725.1 | 3.83% | motif file (matrix) | svg |
| 19 | C T A G T C G A A C G T A C T G C G T A A T G C A C G T G T A C C G T A A G C T G A T C G T A C | Atf3(bZIP)/GBM-ATF3-ChIP-Seq(GSE33912)/Homer | 1e-2 | -5.189e+00 | 0.1215 | 7.0 | 5.30% | 720.8 | 1.60% | motif file (matrix) | svg |
| 20 | C T A G T C G A C G A T C T A G G C A T C A G T C T A G G A T C C G T A G T C A | CEBP:AP1(bZIP)/ThioMac-CEBPb-ChIP-Seq(GSE21512)/Homer | 1e-2 | -5.167e+00 | 0.1215 | 7.0 | 5.30% | 723.9 | 1.61% | motif file (matrix) | svg |
| 21 | A G T C G C A T C G T A C G T A G T A C A C G T A C T G G A T C G A T C T C G A | BMYB(HTH)/Hela-BMYB-ChIP-Seq(GSE27030)/Homer | 1e-2 | -5.056e+00 | 0.1256 | 23.0 | 17.42% | 4525.3 | 10.05% | motif file (matrix) | svg |
| 22 | G T A C G C A T C T G A C G T A G A C T A G C T C A T G T G C A C T G A A C G T G A C T C G T A | Prop1(Homeobox)/GHFT1-PROP1.biotin-ChIP-Seq(GSE77302)/Homer | 1e-2 | -5.006e+00 | 0.1261 | 3.0 | 2.27% | 129.0 | 0.29% | motif file (matrix) | svg |
| 23 | T C G A C T G A T A G C T G A C T C A G T C A G C G T A C G T A T C A G A G C T | ETV1(ETS)/GIST48-ETV1-ChIP-Seq(GSE22441)/Homer | 1e-2 | -4.997e+00 | 0.1261 | 34.0 | 25.76% | 7621.9 | 16.93% | motif file (matrix) | svg |
| 24 | A T G C A G T C C T G A A G T C C G A T A C G T A G T C A G T C A C G T A T C G G A C T A C G T | Etv2(ETS)/ES-ER71-ChIP-Seq(GSE59402)/Homer(0.967) | 1e-2 | -4.817e+00 | 0.1396 | 21.0 | 15.91% | 4084.7 | 9.08% | motif file (matrix) | svg |
| 25 | T C G A G C A T A C T G C T G A A G T C T C A G G A C T G T A C C G T A A G C T A G T C G A T C | c-Jun-CRE(bZIP)/K562-cJun-ChIP-Seq(GSE31477)/Homer | 1e-2 | -4.808e+00 | 0.1396 | 8.0 | 6.06% | 971.8 | 2.16% | motif file (matrix) | svg |
| 26 | T A C G T C G A G A C T A C T G C T G A A G T C T C A G G A C T T G A C C T G A | Atf1(bZIP)/K562-ATF1-ChIP-Seq(GSE31477)/Homer | 1e-2 | -4.807e+00 | 0.1396 | 14.0 | 10.61% | 2312.5 | 5.14% | motif file (matrix) | svg |
| 27 | A C T G C T A G T C G A C G A T C A T G G C T A A T C G C G A T G T A C G C T A A G C T G T A C | Fra1(bZIP)/BT549-Fra1-ChIP-Seq(GSE46166)/Homer | 1e-2 | -4.789e+00 | 0.1396 | 6.0 | 4.55% | 592.1 | 1.32% | motif file (matrix) | svg |
| 28 | A G T C C T A G C T A G A G T C G A T C G T A C A G T C C T A G A G T C A G T C A G T C G T A C | Sp2(Zf)/HEK293-Sp2.eGFP-ChIP-Seq(Encode)/Homer | 1e-2 | -4.768e+00 | 0.1396 | 75.0 | 56.82% | 20734.7 | 46.07% | motif file (matrix) | svg |
| 29 | C T A G T C G A C G A T A C T G C G T A T A C G A G C T T G A C G C T A A C G T G A T C T A G C | Fosl2(bZIP)/3T3L1-Fosl2-ChIP-Seq(GSE56872)/Homer | 1e-2 | -4.611e+00 | 0.1419 | 5.0 | 3.79% | 441.0 | 0.98% | motif file (matrix) | svg |
